# Linking collective migration/growth to differentiation boosts global shaping of the transcriptome and exhibits a grasshopper effect for driving maturation

**DOI:** 10.1101/2022.07.24.501313

**Authors:** Ogechi Ogoke, Daniel Guiggey, Alexander Chiang, Sarah Thompson, Tram Hoang Anh Nguyen, Daniel Berke, Cortney Ott, Allison Kalinousky, Claire Shamul, Peter Chen, Shatoni Ross, Zhaowei Chen, Pooja Srivastava, Chris Gaughan, Supriya Mahajan, Ruogang Zhao, Rudyanto Gunawan, Natesh Parashurama

**Affiliations:** Department of Chemical and Biological Engineering, University at Buffalo (State University of New York), Furnas Hall, Buffalo, NY 14260; Clinical and Translation Research Center (CTRC), University at Buffalo (State University of New York), 875 Ellicott St., Buffalo, NY 14203; Department of Biomedical Engineering, University at Buffalo (State University of New York), Furnas Hall, Buffalo, NY 14260; Center for Cell, Gene and Tissue Engineering, University at Buffalo, State University of New York, Furnas Hall Buffalo NY 14260; Khufu Therapeutics, Inc., San Francisco, CA 94123

**Author notes:** Correspondence Author: Natesh Parashurama, 906 Furnas Hall, Buffalo, NY 14260. Contributed equally to this work (Ogechi Ogoke, Daniel Guiggey, Alexander Chiang).

**Keywords:** organogenesis, organoid, hepatic differentiation, liver development, collective migration, hepatic nuclear network, human pluripotent stem cells, liver diverticulum, liver bud, hepatoblasts, FOXA, Hippo

## Abstract

The shift from collective migration to differentiation is a crucial process in epithelial biology but recreating this intricate transition has thus far proved elusive. We provide experimental, mechanistic, *in vivo*, and bioinformatic data supporting an undoubtable link between human pluripotent stem cell (hPSC)- derived collectively migrating hepatoblasts (MHB), and transcriptionally mature, functional hPSC- hepatocytes (HEPs), which incorporates two unrecognized steps. The protocol induces FOXA-dependent induction of HBs, leading to TBX3-positive, YAP-TEAD active MHB’s which provide a transcriptional match with murine liver E9.5 MHBs. Simple cultivation changes trigger MHB’s to rapidly form functional day 18 HEPs, predicted by a deep-learning designed gene circuit, resulting in a ∼236% fold-increase in maturation (PACNet), on par with the highest score, but with enhanced global transcriptional shaping. Overall, incorporating the MHB to HEP transition establishes a new, unrecognized, and highly efficient mechanism for differentiation that can be cumulatively integrated with existing methods to overcome barriers to maturation.

## INTRODUCTION

Human pluripotent stem cell (hPSC) differentiation is a powerful approach for generating an unlimited amount of differentiated cells applicable to disease modeling, drug discovery, and therapy ^1^. Epithelial organs (e.g., liver, lung, pancreas, and thyroid) exhibit transcriptional, structural, and functional complexity ^2, 3^ and therefore are notoriously difficult to differentiate using human pluripotent stem cells (hPSCs). Further, histological analysis of these epithelial organs demonstrates morphogenesis and patterned three dimensional (3D) structural and functional units (e.g., lung alveolus) by integrating migration, growth, and differentiation. Early epithelial organ development typically involves migration before maturation, and some examples at early stages include pancreatic Ngn3 + endocrine progenitors that migrate into the mesenchyme to form clusters or primitive clusters (islets) ^4^, Nkx2.1 +/ Pax 8 + thyroid progenitors migrate and proliferate towards the pre-tracheal position to form thyroid lobes ^5^. Nonetheless, hPSC protocols employing directed differentiation with soluble factor signaling, small molecule addition, transcription factor (TF) programming, and/or organoids, have not focused on migrating epithelial cells or their relation to maturation. Therefore, how hPSCs recreate collective migration, and how this leads to transcriptional and functional maturation are unknown.

The liver is an extremely pertinent model to investigate this link between migration and differentiation, because there are key developmental and physiological processes that suggest a strong connection. In early liver organogenesis ^6^, E9.5 migrating hepatoblasts (MHBs) self-organize into hepatic cords, branching into adherent migrating stands of HBs within surrounding MESC ^7^. These MHB form more mature HBs that lie within small niches bearing endothelial, mesenchymal, and hematopoietic stem cells (HSC), initiating the formation of hepatic sinusoid architecture by E11.5 ^8^. In adults, acute regeneration after surgery (hepatectomy), transplantation, or chemical insult (CCl_4_) ^9–11^ results in a major proliferative response that replaces the entire mass ^12^. Although local HEP proliferation is responsible, acute regeneration in mouse and humans is accompanied by an ANXA2+ migrating population that employ sheet-based wound closure prior to HEP replication ^13^. These recent regeneration studies establishing migration in human and mouse regeneration, together with recent models of human HEP migration in cell lines ^14^ and the associated difficulty in HEP differentiation, suggest that mechanisms of HEP migration and its role in differentiation need re-examination.

hPSC have provided a powerful tool to recreate developmental or physiological processes, and to model and treat disease, but it remains unknown how to induce and maintain collective migration. The conditions that establish liver fate during development have been recreated to induce hPSC-HEPs in directed differentiation protocols in monolayer or organoid formats, employing sequentially added soluble factors or small molecules, that correspond to developmental cues ^15–17^ or forced expression of master transcription factors (TFs) ^18–19^ to instruct cell fate, control lineage decisions, and affect maturation. Historically, murine HEP-directed differentiation protocols ^2, 20–21^ were transitioned into human PSC-HEP protocols ^22–27^ without major changes in paradigm. Overall, hPSC-HEP protocols currently employ three stages; 1) hPSC to gut tube (GT) endoderm induction, 2) HB induction from GT, and, 3) HB maturation towards HEPs. hPSC-HEP studies typically employ molecular analysis, functional analysis (ALB secretion, metabolism and detoxification), and orthotopic transplantation *in vivo* and have generally exhibit some level of functional restoration. Liver organoid studies provide more organotypic efforts that enhance heterotypic and homotypic cellular interactions ^28–31^ and matched or exceeded the previous findings. Organoids do provide avenues for collective migration, and migration pathways have been identified ^30^, but conditions for inducing collective migration are not known. Nonetheless, all hPSC-HEPs, regardless of methods, are considered to be at a fetal stage with a lack of terminal differentiation ^32–35^. The link between migration/growth and maturity, and the observed lack of maturity of hPSC-HEPs suggest a missing connection that requires further investigation.

Despite their successes, current studies exhibit a heavy dependence on soluble factors, small molecule inhibitors, and/or programmed TFs, during hepatic progenitor induction and maturation. Further, they employ the same paradigm for stages of HEP differentiation, lack considerations of migration-related growth, and do not routinely employ measures of global transcriptional maturity as part of their functional analysis. We carefully examined early liver organogenesis and realized there may be at least two missing steps in the traditional protocol, based upon the transition from the GT to the LD, followed by formation of MHB, then formation of early HB, and we designed a hPSC-protocol to highlight these steps. We therefore focused on assessing the transition between migration and differentiation and how this helps shape the hepatic transcriptome, which is not currently well understood. Amongst many novel findings, we identify a new effect which we term grasshopper effect, in which migrating cells, when inhibited, mature by skipping several steps in traditional HEP differentiation, and our approach could be applicable to other epithelial cultivations systems with further refinement. (778 Words)

## RESULTS

### Comparative bioinformatics analysis demonstrates common transcriptional expression patterns in MHBs from early organogenesis and regeneration

hPSC-HEP differentiation protocol normally involves three stages, with the middle stage being HB induction (**Fig. 1A**). However, further examination of this middle stage reveals that there are in fact multiple HB populations. This includes early liver specification, or liver diverticulum (LD)-HBs, (E9.0), ^36^ and MHBs (E9.5) (**Fig. 1A**), both of which are critical ^37^ (**Fig. 1B**). MHBs form HBs which constitutes a growth to differentiation transition, but the role of MHBs has generally been ignored. To determine the significance of MHB and the transcriptional phenotypes from MHB to HB transition, we performed comparative bioinformatic analysis of early liver organogenesis ^38^ (**Fig. 1C-H, Sup. Table 1-3**), the migrating subset during human liver regeneration ^13^ (**Fig. 1K-P, Sup. Table 4**), and fetal liver growth lacking formal migration (**Sup.** Fig. 1A-G). Pie chart analysis of pathways demonstrated a relative increase in increase in signaling, gene expression (transcription), cell cycle in MHBs, whereas HBs had reciprocal decrease of these pathways and increase in metabolism (**Fig. 1C**), and similar trends were in seen in migrating regenerating HEPs (**Fig. 1K**), but not in proliferating Fetal HEPs migration (**Sup.** Fig. 1A-G). This data suggests that MHB to HB transition carry’s a unique phenotype. Liver GRN expression analysis demonstrated MHB, expressed TBX3, HHEX, and PROX1, as expected, and decreased ALB, and a reciprocal downregulation of these factors, with upregulation of HNF4A, HNF1A, FOXA3, and CEPBA in HBs (**Fig. 1D**). TBX3 and PROX1 were also up in the migrating subset, together with a relative downregulation in FOXA1/3 and ALB (**Fig. 1L**). Although E9.0 LD was not measured, the data suggests that LD-HB ALB positive cells downregulate ALB expression by E9.5 when forming MHBs (**Fig. 1D**). PACNet HB genes were clearly distinguished (**Fig. 1E**), and similar data was obtained for HEPS after regeneration (**Fig. 1M**). Radar plots demonstrated a right shift for both E9.5 MHB (**Fig. 1F**) and migration regeneration subset (**Fig. 1N**). We examined signaling and metabolic pathways in more detail, which revealed similar signaling pathways increased in MHB and migrating subset, and some similar metabolic pathways increased in E10.5 HB and the mature HEP (**Figure H and Figure P**). Overall, the data suggests strong similarities between human MHB, and human regenerating migrating subsets, including increased TBX3 and similar signaling pathways in the migrating population, with a suppression of oxidative phosphorylation.

**Figure 1.**
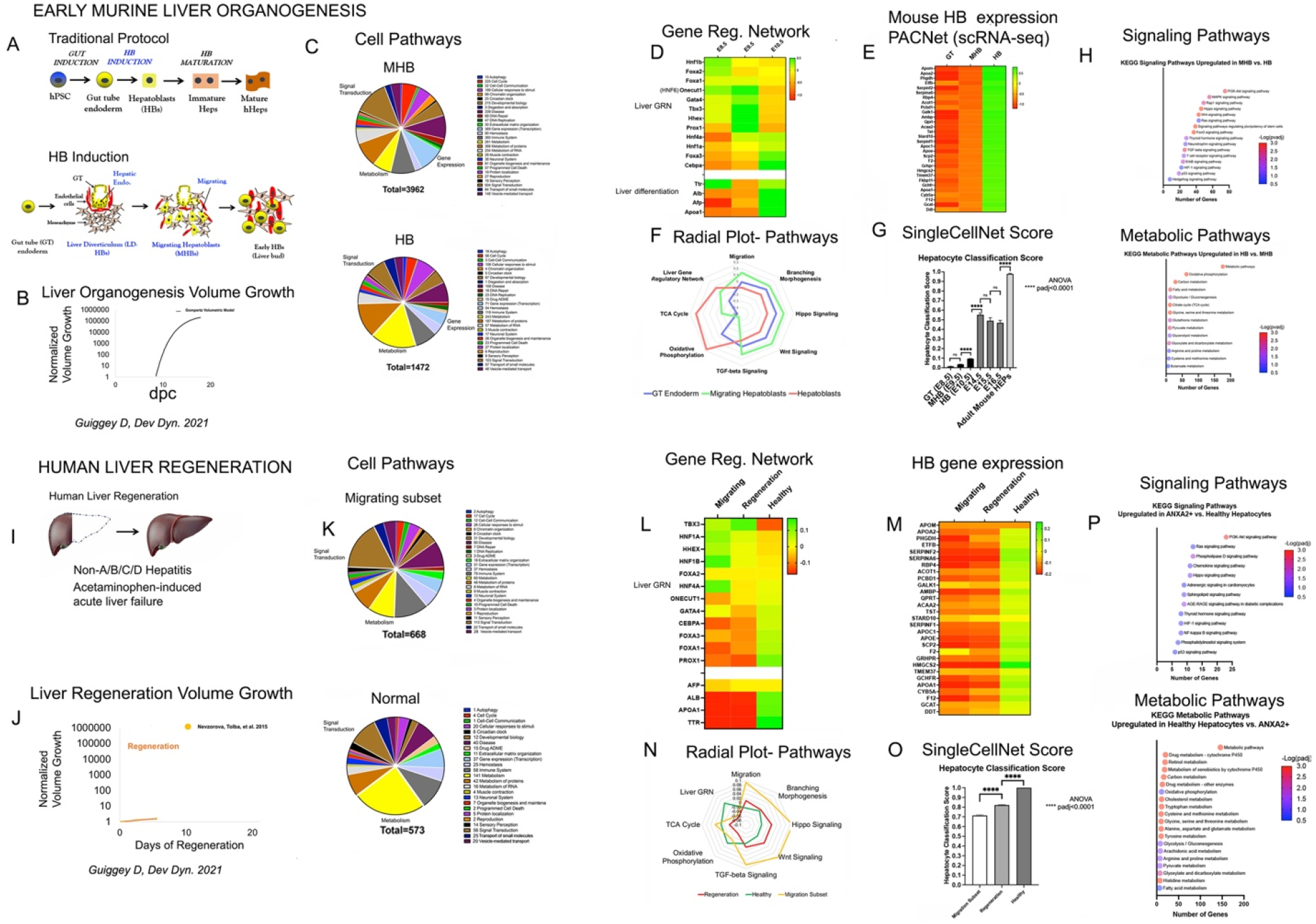
Determination of the migrating hepatoblast (MHB) transcriptional phenotype during murine liver growth and during human liver regeneration A) Schematic of the traditional hepatocyte (HEP) differentiation protocol, alongside proposed missing steps in the gut tube endoderm (GT) and early hepatoblasts (HB) stage. B) Normalized volumetric growth for liver organogenesis for in vivo mouse. C) Pie chart comparison for differentially expressed genes (log2fc > 0.5, padj < 0.05) between migrating hepatoblast (MHB) and HB using Reactome Pathway categories to sort genes. D) Heatmap comparing the average transformed expression values for in vivo mouse hepatic lineage cells (E8.5 (n = 248), E9.5 (n = 668), and E10.5 (n = 1074)) for select important TFs in liver organogenesis (upper) and maturation genes (lower). E) Heatmap comparing the average transformed expression values for in vivo mouse hepatic lineage cells (E8.5 (n = 248), E9.5 (n = 668), and E10.5 (n = 1074)) for top 30 PACNet liver GRN genes that best differentiate MHB (E9.5) and HB (E10.5). F) Radar plot comparing the average transformed expression scores between MHB (n = 668) and HB (n =1074) for select GO pathways based on the differential expression analysis. G) Hepatocyte classification score determined by SingleCellNet and the tabulaMuris classifier. Plotted is mean ± SE. H) Most highly enriched KEGG signaling and metabolic pathways between HB and MHB. I) Diagram of liver regeneration after acute liver injury. J) Normalized volumetric growth for liver regeneration after hepatectomy for adult mouse. K) Same as C) except comparing differentially expressed genes (log2fc > 0.5, padj < 0.05) between migrating subset of cells (ANXA2+) to healthy liver tissue. L) Same as D except comparing cells from the migrating subset (ANXA2+) of regeneration cells, cell populations only found in regenerating liver together and healthy liver cells. M) Same analysis and gene list as E) except comparing cells from the migrating subset (ANXA2+) of regeneration cells, cell populations only found in regenerating liver together and healthy liver cells. N) Same as F) except comparing cells from the migrating subset (ANXA2+) of regeneration cells, cell populations only found in regenerating liver together and healthy liver cells. O) Same as G) except comparing cells from the migrating subset (ANXA2+) of regeneration cells, cell populations only found in regenerating liver together and healthy liver cells P) Same as H) except comparing cells from the migrating subset (ANXA2+) of regeneration cells, cell populations only found in regenerating liver together and healthy liver cells.

**Table 1:**
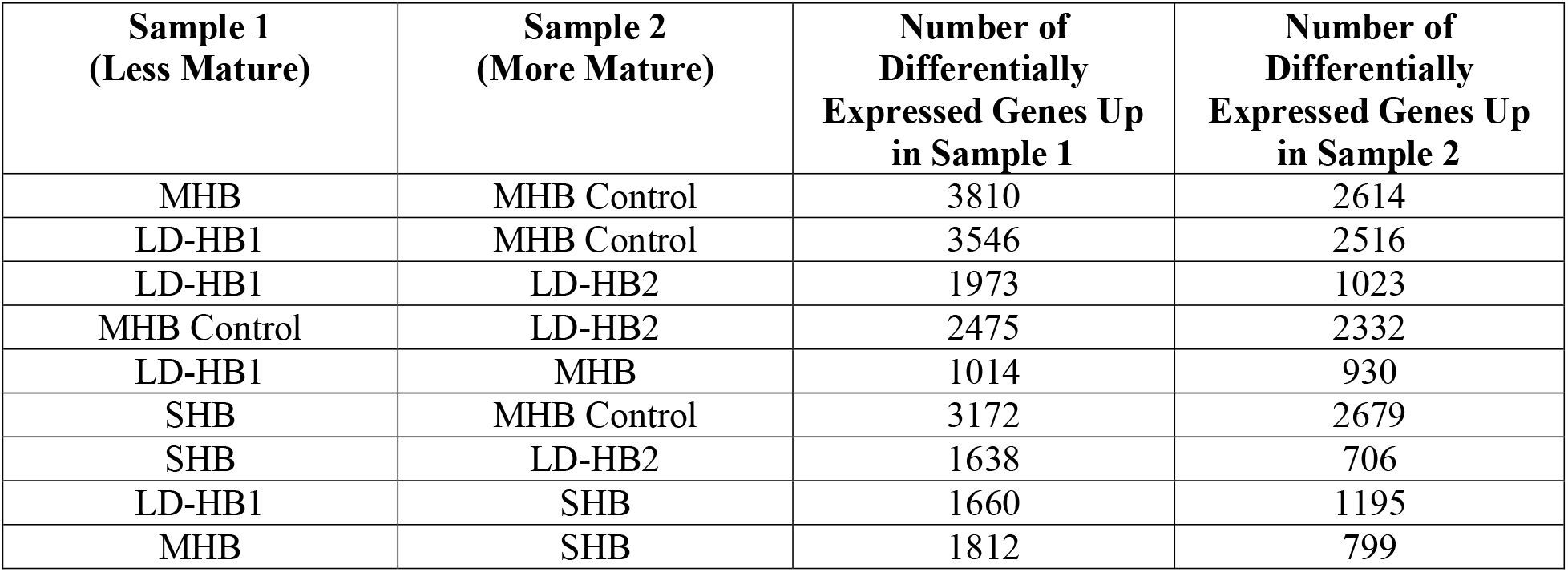
The number of differentiatially expressed genes between different hPSC-HEP sample comparisons.

| **Sample 1**<br>**(Less Mature)** | **Sample 2**<br>**(More Mature)** | **Number of**<br>**Differentially**<br>**Expressed Genes Up**<br>**in Sample 1** | **Number of**<br>**Differentially**<br>**Expressed Genes Up**<br>**in Sample 2** |
| --- | --- | --- | --- |
| MHB | MHB Control | 3810 | 2614 |
| LD-HB1 | MHB Control | 3546 | 2516 |
| LD-HB1 | LD-HB2 | 1973 | 1023 |
| MHB Control | LD-HB2 | 2475 | 2332 |
| LD-HB1 | MHB | 1014 | 930 |
| SHB | MHB Control | 3172 | 2679 |
| SHB | LD-HB2 | 1638 | 706 |
| LD-HB1 | SHB | 1660 | 1195 |
| MHB | SHB | 1812 | 799 |

### SHBs form (monolayer) on Day 14 in H + M medium containing regenerative mediators

We recently developed spontaneously induced HB culture, which we term SHBs in monolayer without instructive factors ^39^. Using this same concept, we added additional mediators of liver regeneration (VEGF, EGF, FGF2, and IGF-1) ^40^ which we identified from ENRICHR pathway analysis in order to prime cells for growth/migration (**Fig. 1A**). These factors also mediate liver differentiation and growth ^24, 30, 41–44^ . The 4 populations formed from day 14 monolayer SHBs are day 18 SHB control (monolayer in H + M medium), day 18 MHBs (organoids in matrigel droplets in H + M medium), day 18 LD-HBs (organoids in suspension in H + M medium), and day 18 MHB Control (same as MHB, but on day 16 medium switched to control medium) (**Fig. 1N**).

To validate the new protocol, we evaluated the day 14 H + M HB protocol and compared it two published protocols, spontaneous differentiation with no addition growth factors, GF - ^39^, and traditional, published methods, GF + ^29^ (**Fig. 2A**), all under hypoxia. Day 4 definitive endoderm (DE) (**Sup.** Fig. 2A**- E**) demonstrated cuboidal cells with bright cell borders indicating an epithelial state and FOXA2 and SOX17 expression (**Fig. 2D, top, left**). In the H + M condition, compared to GF - and GF + conditions, we observed a more elongated morphology by day 12 (**Fig. 2D, top, right**), with epithelial and non-epithelial elements (**Fig. 2D, bottom, left**), whereas we observed more cuboidal epithelial morphology in the GF - and H + M conditions (**Fig. 2D, bottom left, bottom right**). AFP and ALB gene expression were significantly lower in the GF + condition compared to both GF - and H + M conditions, suggesting HBs were low or absent **Fig. 2E**). In H +M compared to GF -, we observed elevated levels of AFP and ALB (**Fig. 2E**), and PROX1 (early liver and bile duct), and CDX2 (hindgut and intestine) was significantly downregulated (**Fig. 2E**). Based on these data, as well as our time course studies (**Fig. 2F**), we employed the day 14 H + M condition (termed SHBs) for further experiments. Functional (ELISA) analysis demonstrated a steady increase in ALB secretion from day 4 to 14, although ALB secretion was still low compared to human functional HEPs ^45^ (**Fig. 2G**). We performed immunoanalysis of the day 14 GF - and H + M conditions (**Fig. 2H-J, Sup.** Fig. 2F-J) and we observed high AFP expression and low/intermediate levels of ALB expression for both conditions (**Fig. 2H**). In the day 6 H + M condition, we observed no ALB expression, low levels of CDX2 expression (mid and hindgut), and SOX2 (foregut), consistent with ventral foregut endoderm (**Fig. 2I**). In the day 14 H + M condition, we observed heterogeneous CD31 (endothelial) expression, but not in the GF (-) condition (**Fig. 2I**), consistent with transient endothelial expression during HEP differentiation ^46^. Accordingly, we observed nuclear expression of FOXA2 and HNF4A in both the day 14 GF - and H + M conditions (**Fig. 2J**). CDX2 expression was lower in the H + M condition, as expected (**Fig. 2J**). We conclude that H + M treated cells were an early HB population. We performed long term culture in H + M medium, which showed an AFP +, ALB +, and TBX3 + cell population (**Sup.** Fig. 2K). Day 14 SHBs were stable in hypoxic culture up to day 24 and presumably beyond. The data suggests the H + M protocol results in a robust, early, SHB population. Importantly, this protocol was tested on multiple iPSC and ESC cell lines with similar results from gene and protein analysis (**Sup.** Fig. 3).

**Figure 2.**
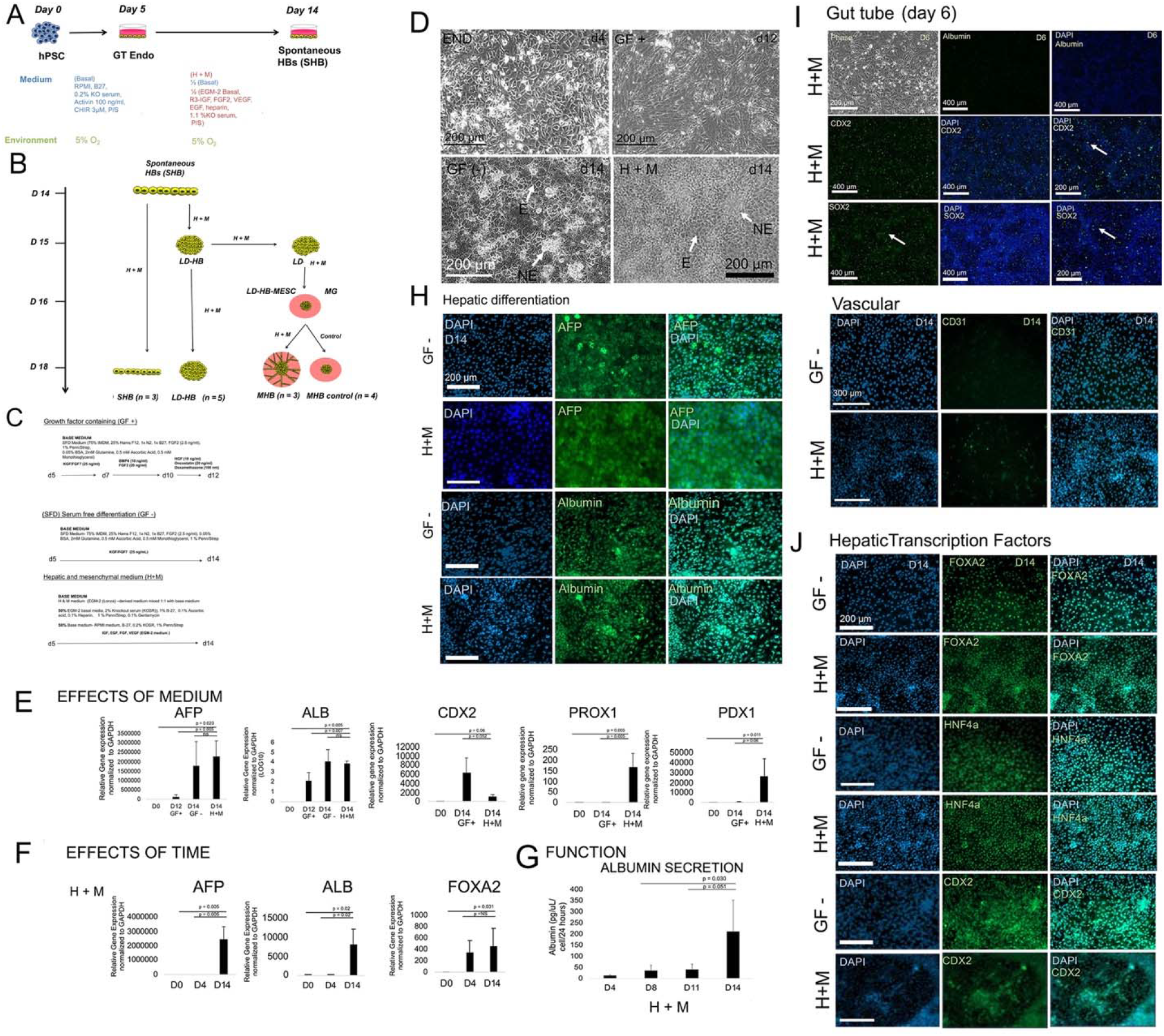
*In vitro* human pluripotent stem cell (hPSC) protocol for inducing MHB and analyzing growth to differentiation switching A) Schematic summarizing early hepatoblast (HB) differentiation in monolayer for the H + M differentiation protocol. B) Overall schematic summarizing the protocols to achieve the four conditions (SHB, LD-HB, MHB, MHB Control) C) Three protocols for hPSC-HB induction (5% O_2_); Growth factor (GF (+)) protocol based upon published work ^39^; GF (-) protocol with serum-free SFD medium, H + M protocol EGM-2 modified medium. D) Morphological analysis during HB differentiation. Endoderm (END)-cuboidal; GF (+): elongated; GF (-): cuboidal; epithelial (E) and non-epithelial (NE) elements (arrows); H + M condition: cuboidal, and NE elements. E) Gene expression analysis (qRT-PCR) of the effects of medium on hepatic differentiation; GF (+) (n = 3), GF (-) (n =3), and H + M (n = 4); mean ± SD. F) Same as K except effects of time: day 0 (n = 3), day 4 (n = 3), and day 14 (n = 3); mean ± SD. G) Enzyme-linked immunoabsorbance assay (ELISA) analysis for ALB secretion; day 8 (n = 4), day 11 (n = 5), day 14 (n = 3). mean ± SD. H) Immunocytochemistry of day 14 hPSC-derived HBs in H + M and GF (-) conditions. DAPI (UV filter), FITC, and merged (UV and FITC) are shown. HBs are stained for AFP (above) and ALB (below). I) Same methods as in H) except hPSC-derived GT endoderm (day 6 cells) were stained by immunocytochemistry, and ALB (liver), CDX2 (hindgut), and SOX2 (foregut) were targeted. J) Same methods as in H) except CD31 (vascular differentiation) was targeted. K) Same methods as in H) except hepatic TFs FOXA2, HN4A, and gut TFs were assessed targeted, as was the intestinal marker CDX2.

### Day 15 liver organoids submerged in MG demonstrate collective migration and growth due to H + M treatment

To recreate the interactions between the LD and surrounding mesoderm tissue complex (MES), we employed day 15 liver organoids submerged in MG in H + M medium which initiated outward collective migration MHBs. We first elucidated factors which enable organoid compaction, or condensation (**Sup.** Fig. 4A). We found MES-derived HUVEC cells cultured with EGM-2 medium (**Sup.** Fig. 4B) and H + M medium alone (**Sup.** Fig. 4C) enabled compaction of day 6 GT cells, although many other approaches failed. Day 14 hPSC-HBs in H + M medium resulted in compaction compared to controls (**Fig. 3A-B**), confirmed by dye-labeling studies (**Sup.** Fig. 4D). Tissue sectioning demonstrated well-organized compact tissue with clusters of epithelial cells and cystic regions (**Fig. 3C**). Immunostaining of organoids demonstrated evidence of ALB (**Fig. 3D**) and AFP expression (**Sup.** Fig. 4E), with minimal cell death (**Sup.** Fig. 4F). After continuous treatment of H + M medium, on day 15, we observed extensive growth and collective migration in all cases, but not in control medium (**Fig. 3E, top**). 96-well systems submerged with MG showed similar results (**Sup.** Fig. 4G). Control medium demonstrated no outward growth/ migration from days 16-18 (**Fig. 3F, top two panels, inset**), whereas H + M medium resulted in radial finger-like migrating protrusions with evidence of branching and webbing (**Fig. 3F, bottom two panels, inset)**, shown at higher magnification (**Fig. 3G**). In some cases, we observed miniature, peripheral cystic structures in control-treated organoids (**Fig. 3G, left)**, while in the H + M condition, we observed migrating, branching cell strands that extend well over 100 µm (**Fig. 3G, right**). The filtering of out-of-focus cells clearly highlights this phenotype (**Fig. 3H)**. For adherent organoids, we again observed minimal collective migration in the control, but conversely, extensive migration in the H + M treated condition (**Sup.** Fig. 4H). and the data demonstrated that collective migration and growth were linked (**Sup.** Fig. 4H). We also evaluated collagen gels (CG) droplets instead of MG, as a model of stiff (2 mg/ml) matrix (**Sup.** Fig. 4I-K) and in contrast to the MG condition, observed sheet-like outward growth. We hypothesized MHBs co-expresses liver and mesodermal protein expression. This was based upon evidence of mesoderm emergence from the LD, as well as the potential presence of hepato-mesenchymal cells ^38^. Organoid immunostaining showed AFP was expressed in both the control and the H + M condition (**Fig. 3I, left, Sup.** Fig. 4L) and H+M treated organoids expressed CD31 (**Fig. 3I, middle**) and SMA (**Fig. 3I, (right)**), and TBX (**Sup.** Fig. 4L). Further, we found migrating strands were indeed AFP (**Sup.** Fig. 4L) and ALB positive (**Sup.** Fig. 4N). We performed ELISA for ALB secretion (**Fig. 3J, left**). The SHB condition showed low ALB secretion. ALB secretion was higher in MHB Control vs. MHB condition, and all conditions were significantly lower than HepG2 cell secretion (**Fig. 3J, left)**. Urea secretion in SHB culture showed an increase but then a significant decrease from day 14 to day 18. (**Fig. 3J, right)**. The MHB condition secreted significantly more urea than the MHB Control and NHDF condition and was significantly lower than HepG2 (**Fig. 3J, right)**. Thus, the MHB condition demonstrated lower ALB secretion, but higher urea secretion when compared to the MHB Control treated organoids. We also performed long term culture of day 18 MHBs until day 30 (**Fig. 3K**). Morphological analysis of day 30 MHBs shows progressive rapid and irregular growth accompanied by migration (**Fig. 3K**), and immunoanalysis demonstrates stable ALB expression (**Fig. 3K**, **middle and bottom rows**). This suggests the cells are stable and robust in long culture to day 30. Overall, our data suggests that the MHBs display an AFP positive, ALB positive, SMA positive phenotype, suggesting a partial mesenchymal phenotype, and secreted higher urea than LD-HB organoids, and demonstrating robust culture through at least Day 30.

**Figure 3.**
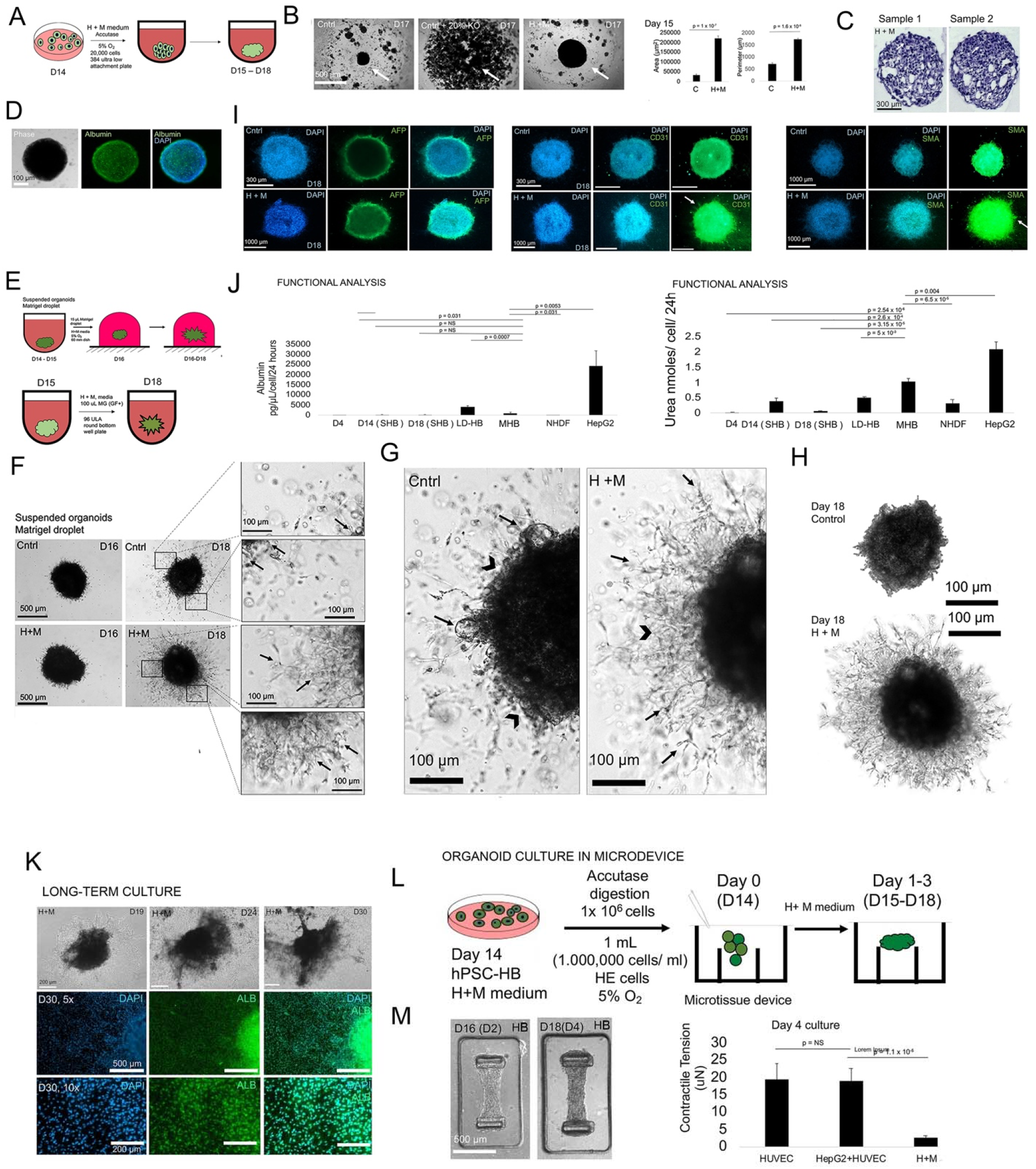
Induction of collectively migrating liver organoids from hPSC-HB A) Schematic of day 14 hPSC-HB organoid formation B) Day 17 images hPSC-HB organoid compaction. Left: control RPMI basal; Middle: RPMI basal medium + 20% KOSR; Right: H + M; Right: Bar graph quantitation: Area (n = 4), perimeter (n = 4); mean ± SD. C) H + E images of day 17 hPSC-HB organoids; above arrows- uniform epithelium; below arrows- non-uniform- cystic like structure. D) Immunofluorescence staining of ALB (middle row), on day 17 hPSC-HB whole organoids. E) Schematic of day hPSC-HB organoid suspended in Matrigel (MG) droplet culture (60 mm dish) or 96-well. F) Phase contrast images of day 18 migrating hPSC-HBs treated in control and H + M medium. G) Same as F except larger; Control: cyst like structures (arrow), minimal CCM (arrowhead). H + M organoids (right) demonstrate CCM. H) Same as F except filtered images to remove out cells that were out of focus. I) Immunocytochemistry of Control (top) and H + M treated (Middle and lower) day 18 whole organoids for AFP, CD31, SMA; counterstained with DAPI and FITC. J) ELISA analysis ALB and urea secretion of day 4 (n = 3), day 14 MONO) (n = 3), day 18 MONO (n = 3), day 18 (SUSP), day 18 organoids (n =3), NHDF cells (n = 3), HepG2 liver hepatoblastoma (n = 3); p-values listed; mean ± SD. K) Phase contrast morphology of replated D19 (left), D24 (center), D30 (right) H+M derived hepatoblast (HB) cells. Immunostaining of iPSC D30 H+M derived HB cells cultured in monolayer for DAPI (left), ALB (center), merged DAPI and ALB (right). L) Schematic for organoid culture in microdevice. M) Phase contrast images of microtissue formed (top) from H+M D14 hPSC-HB cells at D16 (D2) (left) and D18 (D4) (right). Contractile tension comparison (bottom) of HUVEC (left), HepG2+HUVEC (center), H+M hPSC-HB (right). Plotted is mean ± SD.

### Day 18 MHBs display a mesenchymal phenotype in a bioengineered device

Numerous studies demonstrate a hepato-mesenchymal (HM) hybrid phenotype arises during early liver organogenesis ^38^, human fetal liver ^47–48^, and mouse fetal liver ^49^. We tested whether day 18 H + M treated MHBs exhibited a functional, mesenchymal, HM phenotype, by testing their ability to form a microtissue. We used an established bioengineered device that evaluates tissue tension in mesenchymal-derived microtissues ^50^. The device is a microfabricated pillar culture system predicated upon forming a microtissue only in the presence of mesenchymal properties. Preliminary experiments with HepG2, HUVEC, and HepG2/HUVEC cell lines were used to optimize conditions (**Sup Figs. 3O-U**). The functional, mesenchymal potential of day 18 hPSC-HBs was revealed when they formed cell sheets (**Fig. 3L-M**), and contractile tension analysis demonstrated the presence of tension, albeit at significantly lower levels than with cell lines (**Fig. 3M**). These data support the premise that day 18 HBs bear a HM phenotype.

### Human MHB’s express TBX3^+^, FOXA low phenotype with transcriptional upregulation of mesodermal and neuroectodermal TF

To determine the global transcriptional changes, we performed RNA-seq analysis. Principal component (PC) analysis with hPSC-derived populations (MHB, SHB, MHB control, LD-HB), controls, and data from other studies (Li et al. ^51^, Velazquez et al. ^18^), revealed distinct migration/growth versus maturation phenotypes (**Fig. 4A**, **Table 1, Sup. Tables 5-8**). Interestingly, the LD-HB1 group clustered near the migratory MHB population, even though the organoids were in suspension, and there was and could not be, any outward migration. Moreover, even though all LD-HB samples separated into two distinct phenotypes, with LD-HB1 and LD-HB2, as judged by liver GRN expression and radar plot (**Sup.** Fig. 5A). The LD-HB1 demonstrated evidence of migratory population (TBX3 + and shift to right on radar plot), while LD-HB2 demonstrated evidence of maturity (CEBPA, shift to left on the radar plot) (**Sup.** Fig. 5A). SHB represent an intermediate phenotype with partial GRN TF expression and partial overlay with migration on the radar plot (**Sup.** Fig. 5B). These populations were clearly distinguished on the heatmap based upon a migration/growth and differentiation gene lists (**Sup.** Fig. 5C-D). Next, we focused on MHB (migrating) and MHB-Control (differentiated) conditions, and we determined global differences in pathways (**Sup. Table 5**). Pie chart analysis demonstrated a clear switching mechanism; Reactome demonstrated relatively large fractions of Signal Transduction, and Transcription, in the MHB population, while Metabolism and Immune Response was higher in MHB control (**Fig. 4B**), similar to E9.5 MHB (**Fig. 1B**). We noted that both migrating populations (MHB and LD-HB1) had drastically different GRN TF expression than non-migrating populations (**Fig. 4C, Sup. Table 6**). MHB exhibited a FOXA low, TBX3 +, MHB phenotype, and while MHB control exhibited a FOXA1/2 +, TBX3 low, respectively **(Fig. 4C**). Other than TBX3, nearly all hepatic GRN expression were repressed in MHB. We mapped the data on a radar plot and demonstrated that MHB was shifted to the right **(Fig. 4D**), as expected **(Fig. 1F**). We note that Velazquez et. al. and Li. et. al., which provide RNA-seq data over time, do not generate a population that is shifted to the right, suggesting that they may lack a migrating population (**Sup.** Fig. 5F-G). Since a recent study showed FOXA1/2 knockdown results in counter-activation of mesoderm and neural gene expression ^39^, we hypothesized that decreased FOXA factors in MHB causes a similar effect. Interestingly, we identified increased mesodermal and neural TF in MHB, which validated the upregulation of these TF **(Fig. 4E**) and their downstream **(Fig. 4E-F**, **Sup.** Fig. 5E) neural and mesodermal genes. We analyzed FOXA1/2 knockdown data from the Warren et al. study, and FOXA1/2 KD cells were similar to MHB **(Fig. 4G-J**). We observed a similar trend in human migrating regenerating cells (**Sup.** Fig. 5H). To summarize our findings, we generated a schematic of the lineage changes from the MHB to the HB population **(Fig. 4K**) using a corresponding map of GRN TFs in each lineage **(Fig. 4L**). The schematic outlines the observed changes in FOXA1/2, TBX3, and CEBPA. To further validate this, we performed an addition PC analysis based solely upon upregulated neural and mesodermal genes for all populations, and we showed that mature MHB control and migration MHB can be completed separated, as can the related hPSC-derived cell populations **(Fig. 4M**). We further confirm these findings during murine MHB to HB transition **(Fig. 4N**), and we perform analysis of ChIP-Seq for FOXA2 in human liver (HEPG2) cells **(Fig. 4O**), which confirms downstream binding of FOXA2 to mesoderm and neural genes, and similar data was seen for TBX3 and CEBPA **(Sup.** Fig. 4I-J). These findings strongly suggest that the downregulation of FOXA and upregulation of TBX3 correlates with neural TF and neural genes, and mesodermal TF and mesoderm genes in MHB.

**Figure 4.**
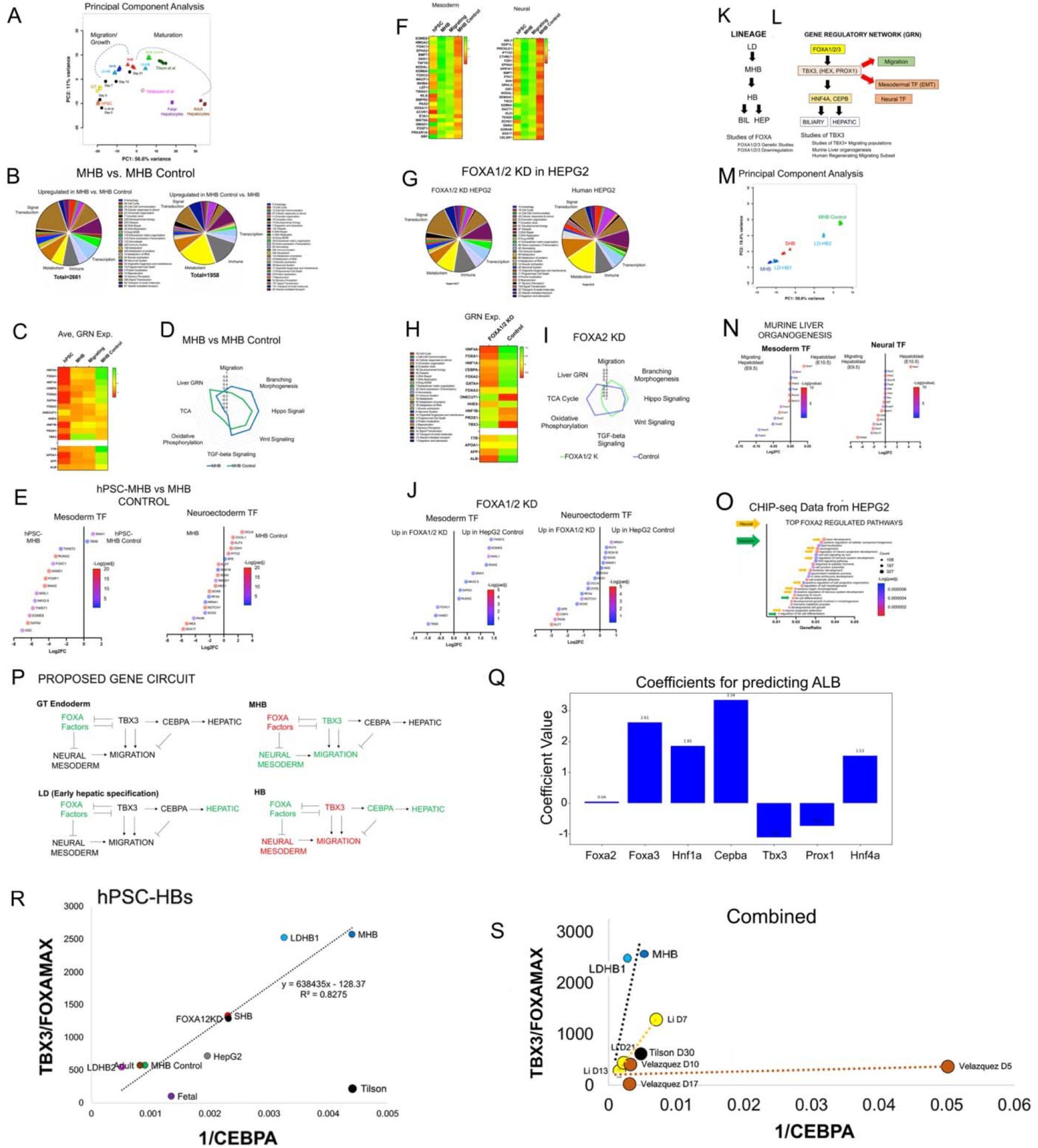
Transcriptomic analysis and modeling of migrating hepatoblasts A) PCA plot for liver and gut genes from Mu et al. analysis. Our five conditions were plotted (hPSC (orange), LD-HB (light blue), MHB (blue), SHB (red), MHB Control (green)) alongside time course data from Li et al. (black), and most mature conditions for Velazquez et al. (pink) and Tilson et al. (green). Additionally controls for human GT (yellow), 12 week fetal hepatocytes (purple), and adult hepatocytes (brown) were also included. B) Pie chart comparison for differentially expressed genes (log2fc >1.5, padj < 0.05) between MHB and MHB Control using Reactome Pathway categories to sort genes. C) Heatmap comparing the average transformed expression values in our conditions (hPSC (n = 2), MHB (n = 3), Migrating (n = 7), and MHB Control (n = 4) for select important TFs in liver organogenesis (upper) and maturation genes (lower). D) Radar plot comparing the average transformed expression scores between MHB (n = 3) and MHB Control (n = 4) for select GO pathways based on the differential expression analysis. E) Gene expression comparisons between MHB (n = 3) and MHB control (n = 4) for select mesoderm and neuroectoderm TFs. F) Lineage diagram with hypothesize role of the GRN based on FOXA and TBX3 studies. G) Same as B) except comparing FOXA1/2 knockdown data (n=3) in HepG2 to control HepG2 (n = 3). H) Same as C) except comparing FOXA1/2 knockdown data (n=3) in HepG2 to control HepG2 (n = 3). I) Same as D) except comparing FOXA1/2 knockdown data (n=3) in HepG2 to control HepG2 (n = 3). J) Same as E) except comparing FOXA1/2 knockdown data (n=3) in HepG2 to control HepG2 (n = 3). K) Heatmap of transformed expression values for top 30 differential expressing mesoderm development genes (GO:0007498) between MHB (n = 3) from MHB Control (n = 4). Control hPSC (n = 2) and migrating condition (n = 7) also shown. L) Same as K except for neuron development genes (GO:0048666) M) Principal component analysis for combined mesoderm (GO:0007498) and neuron GO:0048666) development gene lists with our five conditions (MHB, LD-HB1, SHB, LD-HB2, MHB Control). N) Same as E) except comparing E9.5 MHB cells (n = 668) to E10.5 hepatoblast cells (n = 1074) in mouse development. O) ChIP-seq gene pathway enrichment analysis for FOXA2. Neural (yellow) and mesoderm (green) related pathways are labeled. P) Proposed gene circuit connecting TFs FOXA, TBX3, CEBPA to hepatic development, alternative lineages (neural, mesoderm), and migration based on previous analysis. Q) Correlation analysis for the ratio between TBX3 and largest normalized FOXA gene (FOXA1, FOXA2, FOXA3) with the inverse of CEBPA. Conditions (LD-HB, MHB, SHB, MHB Control) are shown along with control conditions (12 week Fetal and Adult hepatocytes), knockdown data (Control HepG2 and FOXA1/2 KD HepG2), and Tilson et al. data. Time course data for Velazquez et al. and Li et al. plotted separately alongside our data with slopes.

### A three-component gene circuit model composed of TBX3, FOXA, and CEBPA predicts the transition from migration to maturation

We sought to develop a gene circuit that predicts the data in our study. Based on the data (**Fig. 4A-J**), we included FOXA factors (downregulation associated with migration) and TBX3 (upregulation associated with migration) as potential TF. We used HNF4A and CEBPA as maturating TF. A theoretical schematic for the proposed gene circuit describing the GT, the LD, the MHB, and the HB **(Fig. 4P**). FOXA2 is high in GT and assumed to be high in the LD to activate ALB. In MHB, FOXA is downregulated, and TBX3 upregulated, which together activates both migration and alternate mesoderm and neural TFs. Next, TBX3 is inactivated, FOXA (FOXA3) is activated, and CEBPA is activated, which blocks migration and stimulates hepatic maturation **(Fig. 4P**). To validate these TF, we employed a deep learning algorithm called Tensorflow ^52^ modified to identify which TF’s were best correlated with differentiation, since migration is negatively correlated with ALB transcription. Developmental TF networks are believed to be conserved between species, and so we used the early mouse liver organogenesis scRNA-seq data set ^38^. The deep learning analysis found that TBX3 most negatively correlated with ALB expression, while CEBPA and FOXA3 most positively correlated with ALB **(Fig. 4Q**). We first tested the gene circuit with human migrating regenerating data, and a lineage correlation was observed **(Sup.** Fig. 5K). This Migrating Regenerating subset was validated to be migrating using the radar plot and the heatmap validating TBX3 upregulation **(Sup.** Fig. 5L-M). We hypothesized that this conserved gene circuit connecting FOXA, TBX3, and CEPBA would be present in murine FOXA KO models ^53–54^. We first performed heatmap analysis, which reveals de-differentiation, and coordinated changes in FOXA1/2/3 levels, TBX3, and CEBPA levels **(Sup.** Fig. 5N). We then examined the gene circuit correlation, which was strong **(Sup.** Fig. 5O) except for FOXA1/2/3 KO. To demonstrate why FOXA1/2/3 KO cells were not on the curve, we demonstrate that they do not exhibit a migratory phenotype **(Sup.** Fig. 5P-Q). This confirmed that they were not migratory phenotypically, and their GRN expression ensured that they were not on the curve. We then examined the hPSC-HEP populations, including the immature (MHB) and mature MHB Control populations and FOXA1/2 KD data using the gene circuit model. We identified a strong linear correlation for all the data tested **(Fig. 4R**). We also note that Tilson et al. study did not lie upon this line. While most studies did not provide time series data, we examined literature and we found that Velazquez et al. and Li et. al. provided time series data. We plotted these as correlations on the same correlation as the hPSC-HEP data, demonstrating that these examples all lie on different lines **(Fig. 4S**). Overall, studies that display a migrating phenotype seem to follow a similar gene circuit based on correlations, but studies that do not exhibit a migratory phenotype do not, suggesting they have different mechanisms of HEP differentiation.

### FOXA factors control hepatic commitment in human liver cell lines and induce a global de- differentiation phenotype with morphological changes

Given the prominent role of FOXA1/2 in our proposed gene circuit, and a study demonstrating that FOXA3 compensates for FOXA1/2 in murine liver ^53^, we aimed to determine if FOXA1/2/3 KO results in further de-differentiation and activation of a migration phenotype via TBX3 upregulation. We first targeted FOXA1/2/3 to generate a FOXA1/2/3 triple knockdown (TKD) in human liver (HepG2) cells. We observed a morphological change in cells, less circular and larger (**Fig. 5A-B**) compared to scrambled control, with no differences in cell viability (**Sup.** Fig. 7A-B). We found that in FOXA1/2/3 KD cells, FOXA factors were significantly downregulated, in addition to ALB and AFP (**Fig. 5C**), indicating hepatic de-differentiation. Supporting this, we observed significant downregulation of many liver GRN TFs, including HNF4A and HEX, with TBX3 and HNF1B trending downwards (**Fig. 5D**), although we observed no significant changes in HNF6, HNF1A, and PROX1 (**Fig. 5E**). As further evidence of de- differentiation, we observed significant upregulation of pluripotency factors NANOG, and SOX2, and significant downregulation of OCT4 (**Fig. 5F**). We observed no significant changes in definitive endoderm markers CXCR4 or SOX17, but we identified significant GSC and GATA4 downregulation (**Fig. 5G**). Immunostaining confirmed liver de-differentiation (**Sup.** Fig. 7C-D). Western blotting demonstrated downregulation of FOXA1/2/3, HNF4A, and ALB, with upregulation of AFP, CDX2 (gut/intestine), and SOX 17 (DE) (**Fig. 5H-I**). We also analyzed markers for alternate lineages (**Sup.** Fig. 7E) and demonstrated downregulation of major metabolic genes (**Sup.** Fig. 7F). Finally, we observed an increase in lipid droplet accumulation and a decrease in urea production (**Sup.** Fig. 7G-H) consistent with de-differentiation and loss of function. This data suggests that FOXA1/2/3 downregulation did not cause an increase in TBX3 transcription, but did cause a morphology change and strong liver de-differentiation in human liver cells.

**Figure 5.**
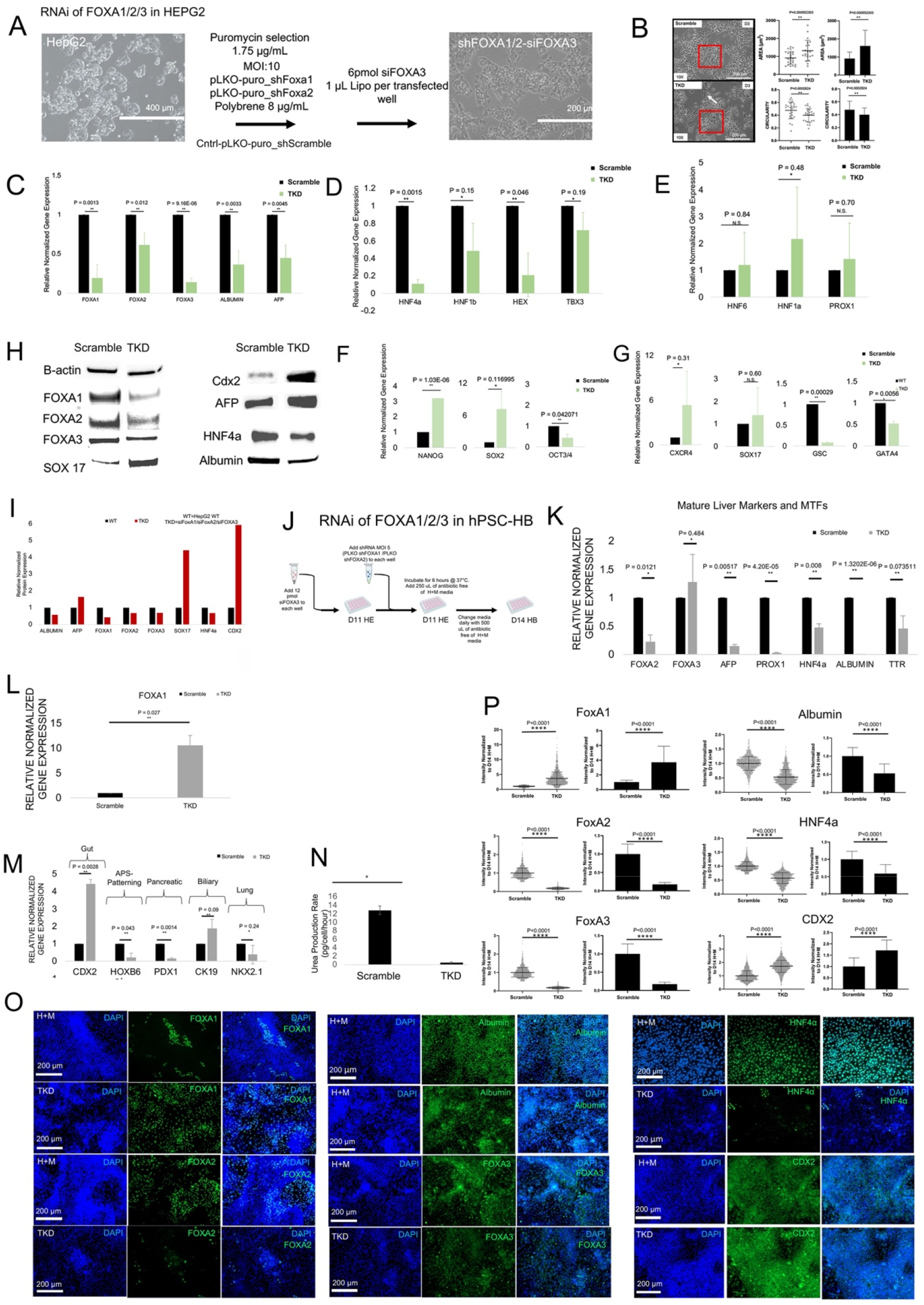
RNAi of FOXA1/2/3 in liver cells A) Schematic for RNA interference for FOXA1/2/3 in HEPG2 cells B) Phase contrast images between shScrambled (top) and FOXA1/2/3 KD (bottom). Average area (top) and circularity (down) for shScrambled and triple knockdown (TKD). C) Bar graph of gene expression kinetics (qRT-PCR) of FOXA1/2/3 KD (TKD) HepG2 compared to shScrambled HepG2 for FOXA1, FOXA2, FOXA3, ALBUMIN, AFP. n = 3 for each group compared. Plotted is mean ± SD. Significance (**) defined as P ≤ 0.05. D) Same as C) except for additional liver developmental TFs that were downregulated in TKD compared to shScrambled, HNF4A, HNF1B, HEX, and TBX3. E) Same as C) except for additional liver development TFs that were upregulated in TKD compared to shScrambled, HNF6, HNF1A, PROX1. F) Same as C) except for neural/ectoderm development markers, NANOG, SOX2, OCT3/4. G) Same as C) except for definitive endoderm markers, CXCR4, SOX17, GSC, GATA4. H) Western blot between shScramble (left) and TKD (right) for beta-actin, FOXA1, FOXA2, FOXA3, SOX17, CDX2, AFP, HNF4A, ALB I) Quantitative analysis of data in H). Significance (*) defined as P ≤ 0.05 (n = 3). J) Schematic for shFOXA1/2 and siFOXA3 knockdown approach on day 11 in the H + M protocol, and assessed cells at day 14. K) Bar graph of gene expression kinetics (qRT-PCR) of FOXA1/2/3 KD (TKD) hPSC-derived D14 HB compared to shScrambled hPSC-derived D14 HB for mature liver markers and TFs FOXA2, FOXA3, AFP, PROX1, HNF4A, ALBUMIN, TTR. N = 3 for each group compared. Plotted is mean ± SD. Significance (*) defined as P ≤ 0.05. L) Same as K) except for FOXA1. M) Same as K) except for alternative lineage liver markers, gut (CDX2), APS-patterning (HOXB6), pancreatic (PDX1), biliary (CK19), lung (NKX2.1). N) Bar graph of urea production rate for FOXA1/2/3 KD (TKD) hPSC-derived D14 HB compared to shScrambled hPSC-derived D14 HB. Plotted is mean ± SD. Significance (*) defined as P ≤ 0.05. O) Immunocytochemistry of FOXA1/2/3 KD (TKD) (bottom) hPSC-derived D14 HB compared to control shScrambled hPSC-derived D14 HB (HM) (top). DAPI (UV filter), FITC, and merged (UV and FITC) are shown. Cells are stained for FOXA1, FOXA2, FOXA3, ALBUMIN, HNF4A, CDX2. P) Quantitative analysis of data in O). Significance (*) defined as P ≤ 0.0001 (****) .

### FOXA factors control hepatic commitment hPSC-HBs but do not induce a partial migratory phenotype

Given this strong phenotype, we wanted to test whether FOXA/1/2/3 KD phenotype in our hPSC- HB induction model in H + M medium results in a de-differentiated, migration phenotype. We established shFOXA1/2 and siFOXA3 knockdown approach on day 11 in the H + M protocol, compared to shScramble control, and assessed cells on day 14 (**Fig. 5J**). Gene expression analysis demonstrated de- differentiation of liver factors, including significant downregulation of FOXA2, AFP, PROX1, HNF4A, ALB, and TTR, indicating hepatic de-differentiation (**Fig. 5K)**. Interestingly, we observed no change in FOXA3 despite siRNA targeting, and we observed an increase in FOXA1 (**Fig. 5L**). In terms of alternate lineages, we observed an upregulation of CDX2 (intestine), and CK19 (biliary) markers, and a downregulation of HOXB6 (gut patterning), PDX1 (pancreatic) and NKX2.1 (lung/thyroid) differentiation (**Fig. 5M**). We again observed a downregulation in urea secretion in the TKD condition (**Fig. 5N)**. Immunostaining demonstrated downregulation of FOXA2, FOXA3, ALB, HNF4A and increases in FOXA1 and CDX2 (**Fig. 5O-P)**. These data establish a hepatic de-differentiation in response to TKD in H + M medium, indicating FOXA factors play a large role in initiating hepatic induction in hPSC-HBs, but we observed no evidence of a TBX3 + migration phenotype.

### Growth-to-differentiation transition *in vivo* leads to extensive tissue growth in a novel ectopic model

The concept of growth to differentiation transition can potentially be used to build tissues *in vivo*. We wanted to establish that *in vivo* growth to differentiation transition is a feasible method for building liver tissue from hPSC, and to do this we developed a simple ectopic transplant model in subcutaneous tissues (**Figs. 6A-B**). A subcutaneous site is advantageous here because it is more likely hypoxic than commonly used ectopic transplant sites, and hypoxia supports early liver growth ^55^. We injected day 5 DE/GT cells to model the LD cell which resulted in a palpable mass after at least two weeks in all conditions tested (**Figs. 6C**). As a control, hPSC were injected subcutaneously to form a teratoma after 4 weeks (**Figs. 6D, top row**). Importantly, we observed various clusters of high ALB expression (**Fig. 6E**). Next, we wanted to demonstrate that HEP-committed cells can thrive ectopically. We implanted HepG2 liver cells and after 4 weeks, a large tissue mass with strands of tumor cells surrounded by cell-free regions was present (**Fig. 6F**). The data demonstrated ALB-low expressing cells with pools of ALB (+) liquid between strands of cells suggesting ALB was secreted into pools of venous blood (**Fig. 6G**), when compared to control (**Sup.** Fig. 8A-C). No formal vascular structures were identified. Next, we tested our hypothesis that day 5 DE/GT transplantation can undergo a growth to differentiation transition and result liver tissue growth and differentiation. We transplanted day 5 DE/GT resulting in an oval-shaped mass with extensive homogeneous tissue growth after 2 weeks. Higher magnification analysis demonstrated evidence of organization, with nests of cellular dense regions (**Fig. 6H**). ALB immunohistochemistry for evidence of liver differentiation demonstrated a large part of the mass (> 50%) and was positive for ALB, mainly focused on the center of the mass, which is noted in dual FITC-DAPI image (**Fig. 6I, DAPI vs. Green**). Clustered cellular networks expressed even higher ALB expression levels (**Fig. 6I, arrows**). The data demonstrates *in vivo* liver differentiation occurs from DE/GT transplants without blood vessels.

**Figure 6.**
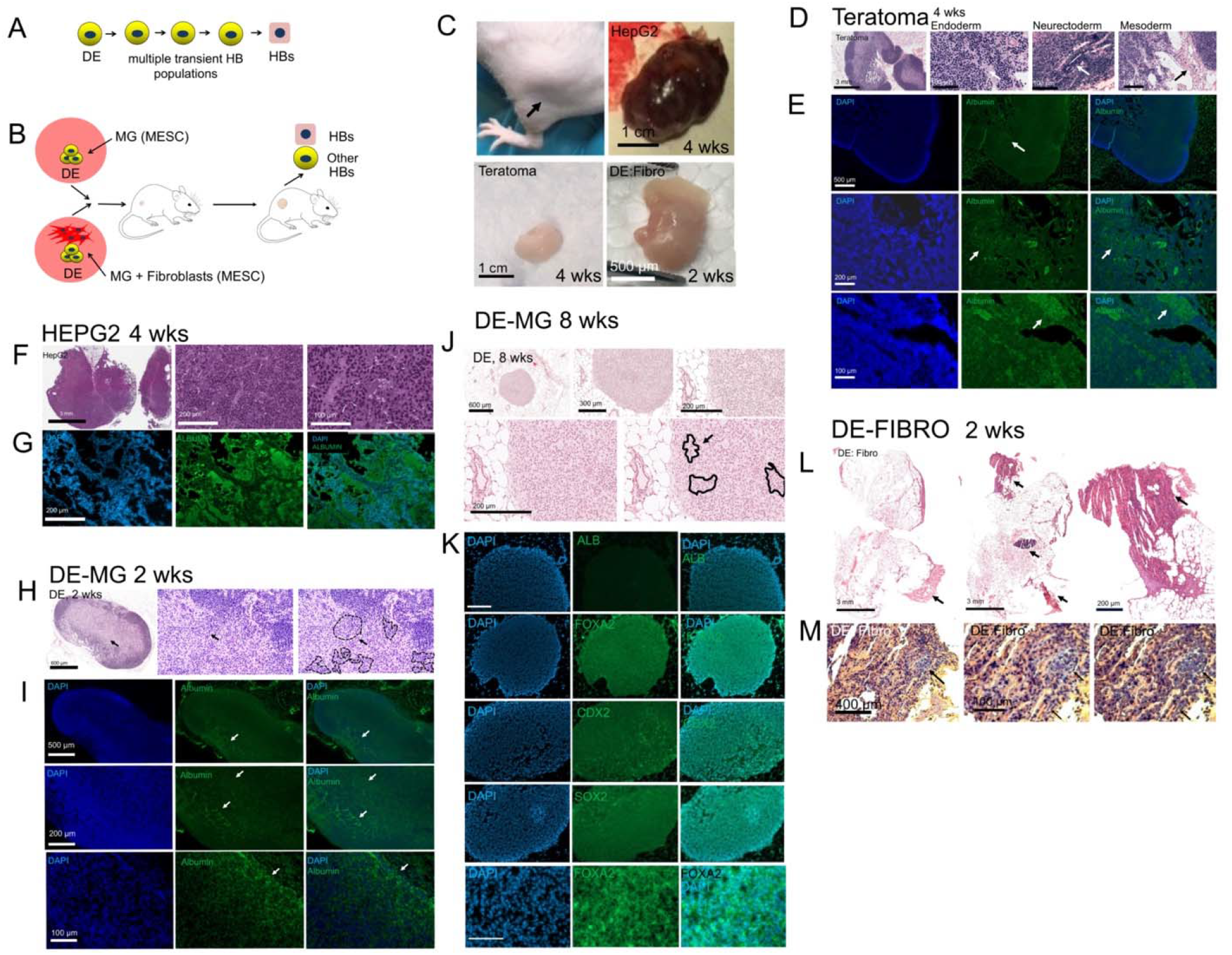
In vivo transplantation to determine effects of growth to differentiation switching A) Schematic of early liver cell differentiation between definitive endoderm (DE) to hepatoblasts (HB). B) Schematic for our mouse in vivo ectopic transplant model. C) Gross images of teratoma, hPSC-DE:HFF, 4 weeks post-transplantation HEPG2 tumor, and DE:Fibro. D) Histological analysis (Hematoxylin and Eosin) of teratoma (neuro-tubular structures (arrow), hPSC-DE, Neurectoderm, and Mesoderm tissue. E) Immunocytochemistry of teratoma in H + M and GF (-) conditions. DAPI (UV filter), FITC, and merged (UV and FITC) are shown. Cells are stained for ALB. F) Gross images of 4 weeks post-transplantation HepG2 tumor. G) Same as E) except for 4 weeks post transplantation HepG2. H) Gross images of DE:MG, 2 weeks post-transplantation. I) Same as E) except for 2 weeks post-transplantation DE:MG J) Tissue section for DE:MG, Arrows show morphological clustering. K) Same as E) except for DE:MG, 8 weeks post-transplantation. L) Tissue section for DE:Fibro 2 weeks post-transplantation. Arrows show scattered growth small areas in fibro-fatty tissue. M) Histological analysis (Hematoxylin and Eosin) of hPSC-DE:HFF (DE-derived (blue) (arow), HFF (orange))

Overall, the data demonstrates day 5 DE/GT can appreciably expand *in vivo* over a two-week time period, demonstrating evidence of growth to differentiation transition. To test the long-term potential of this approach, to we performed long term experiments for 8 weeks. H + E staining demonstrated a round, homogenous mass of cells, with some evidence of morphological clustering (**Fig. 6J**). Surprisingly, we no longer detected ALB at 8 weeks (**Fig. 6K, top row**). We stained for other relevant gut markers including FOXA2, CDX2, SOX2, all of which were positive (**Fig. 6K**). Given that the liver arises from the foregut/midgut junction, the data suggested that cells lost their liver fate, but retained their gut fate. We examined FOXA2 at higher magnification, and we found that FOXA2 staining demonstrated nuclear exclusion (**Fig. 6K, bottom row**). This data suggests that the loss of ALB was due to the loss of nuclear FOXA2 expression *in vivo*, and that the cells potentially took on a foregut (SOX2) or hindgut (CDX2) fate.

In the final experimental model, we hypothesized that supporting cells may enable further growth and differentiation. We employed HFF since we have previously shown that they induce hepatic endoderm migration *in vitro* ^55^. After 4 weeks, the data shows scattered growth of small areas in fibro-fatty tissue, (**Fig. 6L**), with small areas with dense tissue that was clearly due to transplantation (**Fig. 6L, arrows**). Further analysis confirmed a dense 1000 µm (1 mm) wide area, rich with HFF which was particularly stiff upon tissue section (**Fig. 6L, right, arrow**). H + E analysis revealed that the HFF were aligned and colored orange, while the DE-derived cells were colored blue (**Fig. 6M, left, arrow**). The data demonstrates a clear lack of growth. Further, we were unable to assess differentiation because the extremely stiff environment prevented further sectioning However, these data suggest the possibility of collective migration of hPSC-derived (blue) DE/GT strands within sheets of HFF cells (**Fig. 6M, middle, arrow**), and we performed image segmentation, which clearly demonstrates blue, migrating hPSC-DE-derived strands (**Fig. 6M, right arrow**). In summary, HFF mixed with DE/GT enables collective migration, but a growth phase was not reached, perhaps due to excessive stiffness. Overall, we demonstrate evidence of growth and differentiation at 2 weeks post-transplant, with evidence of liver differentiation, growth to differentiation transition, and loss of hepatic differentiation by 8 weeks, and intercellular migration but loss of growth in stiff environments.

### MHB Control and LD-HB2 display comparable liver scores (PACNet) to published studies

We wished to better elucidate the growth to differentiation transition from MHB to MHB Control. In the Fetal to Adult HEP transition, we determined that increased expression of liver GRN (**Sup.** Fig. 9A) correlates well with large increased in transcription between 12-week Fetal HEPs and Adult HEPs (**Sup.** Fig. 9B). Rather than examine all genes, we employed methods to quantitatively measure transcription (PACNet) ^56^, and we performed several analyses to demonstrate PACNet is robust and sensitive (**Fig. 1G and 1O, Sup.** Fig. 9C-F). We hypothesized that hPSC-HEPs are transcriptionally immature, and we analyzed all recent published hPSC-HEP studies that employed RNA-seq (n = 20) (**Table 2**) after eliminating a few for low scores or sample size, and compared with hPSC-HEP populations, with Tilson et. al. study being the highest scoring study (**Fig. 7A**). All published scores were reduced compared to 12- week Fetal HEPs and adult HEPs, and MHB Control and LD-HB2 conditions compared favorably with all published studies (**Fig. 7A**). MHB Control was a result of initiating migration in H + M medium and stopping the process with medium removal. We assumed that LD-HB2 was the result of maturation from an immature LD-HB1 population, without external migration, and solely in the presence of H + M medium. We first compared MHB Control and LH-HB2 to understand their similarities and differences (**Sup. Table 7)**. Comparison of GRN expression demonstrated differences in TF expressed, suggesting that there are multiple potential HEP differentiation pathways (**Sup.** Fig. 9G). Pie chart analysis demonstrated both populations exhibited elevated signaling, but MHB Control displayed elevated Metabolism (**Sup.** Fig. 9H-I). Radar plot analysis demonstrate a strong overlap between these two populations, but with distinctions in signaling and liver GRN (**Sup.** Fig. 9J). When compared directly, MHB Control and LD-HB2 were not significantly different than trans-differentiation using TF reprogramming (6-7 TF) of fibroblasts ^19, 57^ (**Fig. 7B, left**). Compared to the highest scoring organoid ^58^ and monolayer ^15^ studies, both LD-HB2 and MHB-Control compared favorably to Koido et. al., but Tilson et. al. scored higher than both (**Fig. 7B, right**). We observed a variety in culture time, and therefore, we divided the PACNet score by the total number of days of culture, to obtain a normalized scoring system that measures efficiency of differentiation per day of culture. The normalized score demonstrated that our approaches demonstrated excellent efficiency, and our populations were two of top 5 scoring studies in the field, and two of the top 4 without TF programing (**Fig. 7C**).

**Figure 7.**
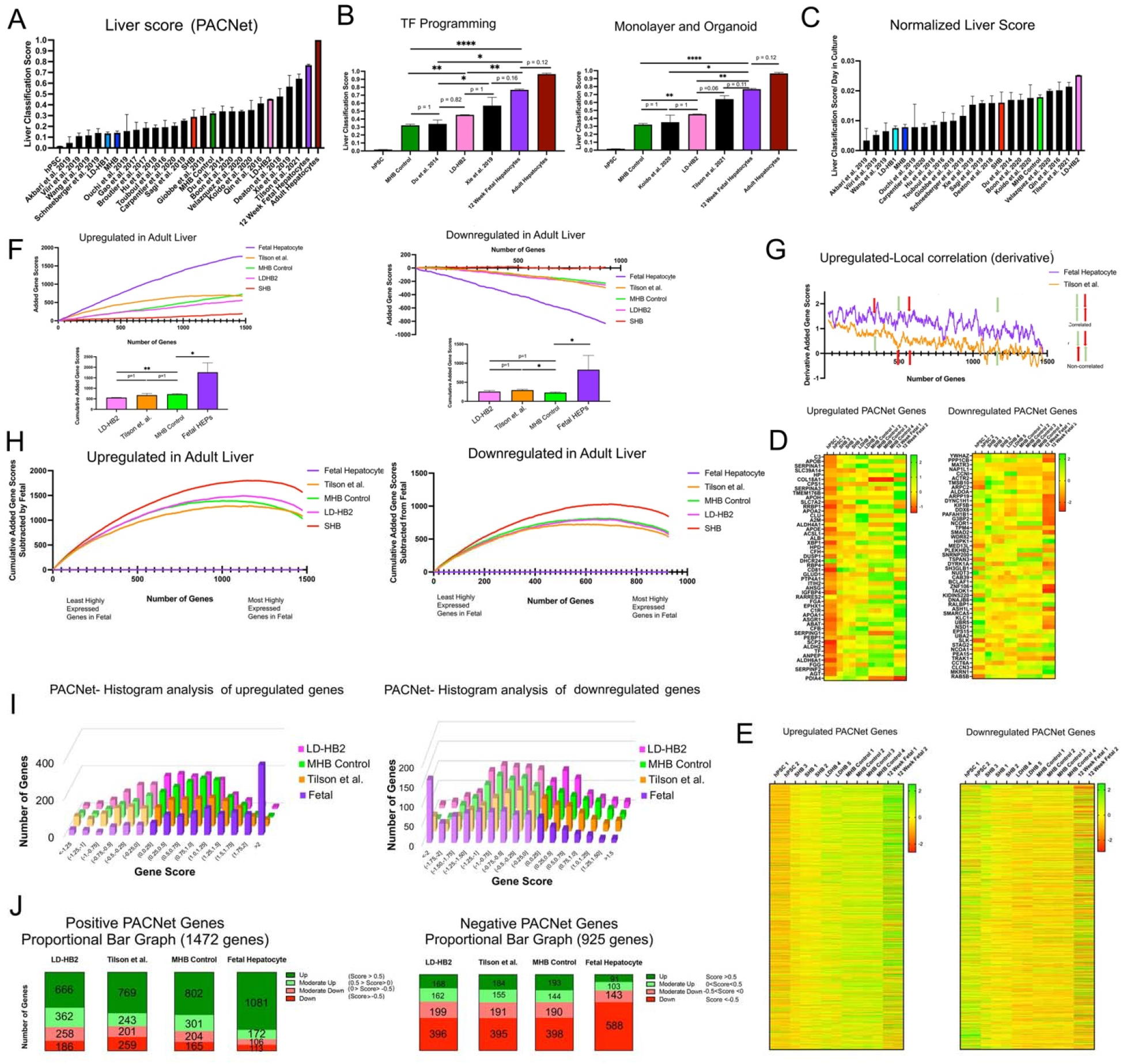
Transcriptional analysis of differentiated populations after growth to differentiation switching A) Comparison of PACNet Liver Classification scores for our samples (colored), controls (hPSC, 12 week fetal HEPs, adult HEPs), and 20 protocols from the literature (black). B) Comparison of PACNet Liver Classification scores for our highest scoring samples (LD-HB2 and MHB Control), with highest scoring samples from the literature that used TF reprogramming (Du et al., Xie et al.) and controls (adult hepatocytes and 12 week fetal hepatocytes). Also showing a comparison with the highest scoring samples from the literature for monolayer culture (Tilson et al.) and organoid (Koido et al.) C) Same as A) except classification scores were normalized to days in culture. D) Transformed expression heatmap for top 50 most highly expressed positive and negative PACNet liver GRN genes sorted by training data E) Transformed expression heatmap for upregulated (1564 genes) and downregulated (932 genes) PACNet liver GRN genes sorted by most highly expressed in the training data F) Added average transformed gene expression scores for upregulated (1564) and downregulated (932) PACNet liver GRN genes sorted by highest expressing in the PACNet training data for our conditions (MHB control (n = 4), LDHB2 (n = 2), SHB (n = 3)), samples from Tilson et al. (n = 4), and fetal hepatocytes (n = 2). Sum of all liver GRN genes was used to statistically compare conditions. G) Derivative of F) showing only fetal hepatocyte and Tilson et al. Select correlated and non-correlated genes were further labeled. H) Same as F) except cumulative gene expression scores were subtracted from fetal for each additional gene added. Order of the genes was also changed to the most highly expressed genes and the least highly expressed genes in fetal hepatocytes for the upregulated PACNet and downregulated PACNet gene lists respectively. I) Histogram for the transformed gene expression scores for the upregulated (1564) and downregulated (932) liver GRN PACNet genes J) Proportional bar graph summarizing the histogram results from I) into four groups (Up, Moderate Up, Moderate Down, Down based on score).

**Table 2:** Studies used for PACNet analysis of liver classification/maturity.

| Source | Journal | PMID | Sample Used |
| --- | --- | --- | --- |
| Akbari et al. 2019 | Stem Cell Reports | 31522975 | endoderm derived hepatic organoids, differentiation medium, WT |
| Boon et al. 2020 | Nature Communications | 32170132 | ESC derived hepatic like cells D20 |
| Broutier et al. 2017 | Nature Medicine | 29131160 | primary liver derived organoid, hepatocyte specification |
| Carpentier et al. 2020 | Gut | 32114504 | D21 ESC-derived HLCs |
| Deaton et al. 2018 | Nature Communications | 30271901 | D20 iPSC derived hepatocytes WT |
| Du et al. 2014 | Cell Stem Cell | 24582926 | ESC-derived hepatocytes |
| Du et al. 2014 | Cell Stem Cell | 24582926 | D30 fibroblast derived hepatocytes |
| Gao et al. 2017 | Stem Cell Reports | 28216149 | iPSC derived hepatocyte like cells |
| Giobbe et al. 2019 | Nature Communications | 31827102 | P30 hepatocyte organoids, standard hydrogel |
| Guan et al. 2017 | Journal of Hepatology | 31760071 | D62 hepatic organoid, differentiation medium |
| Hu et al. 2018 | Cell | 30500538 | primary fetal hepatocyte-derived organoids, P22 |
| Ouchi et al. 2019 | Cell Metabolism | 31155493 | D25 iPSC-derived liver organoids |
| Qin et al. 2016 | Scientific Reports | 27874032 | D18 Derived hepatocytes |
| Sagi et al. 2019 | Cell Stem Cell | 31491396 | Androgenetic ESC-derived WT hepatic cells |
| Schneeberger et al. 2019 | Hepatology | 31715015 | D12 liver organoids, spinner flask |
| Tilson et al. 2021 | Hepatology | 34288010 | iPSC-derived hepatocyte-like cells, WT Control |
| Touboul et al. 2016 | Journal of Hepatology | 26921690 | ESC-derived hepatocytes |
| Velazquez et al. 2020 | Cell Systems | 33290741 | D17 DesLO |
| Viiri et al. 2019 | Scientific Reports | 30837492 | hepatocyte like cell |
| Wang et al. 2019 | Cell Research | 31628434 | hPSC-derived expandable hepatic organoids |
| Xie et al. 2019 | Cell Research | 31270412 | HPLC-derived induced hepatocytes |
| Koido et al. 2020 | Nature Medicine | 32895570 | human liver bud organoids |
| Li et al. 2017 | Nature Communications | 28466868 | Day 21 |

Since the PACNet score, which employs only 100 upregulated and 100 downregulated genes, may represent either a measure of gene activation (whether gene is expressed or not), or transcriptional up- or down-regulation (whether transcripts are expressed or repressed at appropriate levels), we wanted to more systematically measure transcription. When we performed heat map analysis of the top 100 upregulated and top 100 downregulated genes, certain PACNet adult HEP upregulated genes were often not upregulated in the 12-week fetal HBs, like APOB or COL18A1 (**Fig. 7D)**, and the mis-regulation of genes was also observed for the PACNet adult HEP downregulated genes. There were also discrepancies between hPSC-HEP samples and Fetal HEPs for both expected upregulated and downregulated genes. We visualized these differences across all PACNet genes, which enabled visualization of the non-correlation between Fetal and Adult HEPs, and between the hPSC-HB populations and Fetal HEPs (**Fig. 7E)**. Overall, we demonstrate two of the top five scores for measurement of differentiation efficiency, but we observe mis-regulation or lack of matching between Fetal and adult HEP genes expression trends, and between hPSC-HEPs and Fetal HEPs trends.

### MHB Control demonstrate advantages in transcriptional shaping

We defined transcriptional shaping as the ability for the sample to match upregulation of Fetal HEP upregulated genes, and downregulation of Fetal HEP downregulated genes. The PACNet gene lists contain 1477 upregulated adult liver genes and 925 downregulated adult liver genes. We show that genes beyond the first 100 also can contribute to score, suggesting all genes are important (**Sup.** Fig. 7K). To account for this, we first plotted cumulative, transformed gene expression data (Rlog) (see methods) for scoring upregulated and downregulated scores (**Fig. 7F**). These plots, based upon PACNet adult liver genes from highest significance to lowest significance, demonstrate Fetal HEPs have highest cumulative scores (reflecting increased gene expression). Surprisingly, the MHB population cumulative score passes that of the Tilson et. al., the highest scoring study, but not to the level of fetal HEPs (**Fig. 7F, left**). Accordingly, analysis demonstrates no significant difference between MHB and Tilson (**Fig. 7F, bottom left**). On the other hand, for the downregulated adult HEP PACNet genes, the cumulative (negative) gene scores of SHB was near zero, and Tilson et al. ended up lowest among experimental samples, but the experimental groups were much closer to each other (within 30 points), and all were therefore nearly equally lower than Fetal HEPs by approximately 600 points (**Fig. 7F, right**). The MHB Control was significantly lower than Tilson et al. (**Fig. 7F, bottom right**). However, our approach did not take into account gene-by-gene differences, as illustrated with an instantaneous (surface) derivative of the cumulative scores in **Fig. 7G**, which enables visualization of mismatching or lack of correlation between scores in Fetal HEPs and Tilson et. al., experimental conditions (**Fig. 7G)**. Therefore, we sorted PACNet gene list based upon fetal HEP expression and replotted differences in scores from experimental samples and Fetal HEPs (**Fig. 7H, left**). For upregulated genes, MHB control retained its advantage compared to Tilson et. al. (**Fig. 7H, left**). For downregulated genes, both Tilson et. al. and experimental conditions grouped together and did not match Fetal HEPs (**Fig. 7H, right**). To further understand differences in scores, we grouped gene expression levels across genes into batches of expression (**Fig. 7I**). On a 3D plot, we see all conditions and their expression levels from least expressed gene (left) to most expressed genes (right) for the upregulated gene list (**Fig. 7I, left**) and the downregulated gene list (**Fig. 7I, right**). The upregulated gene list confirmed that fetal HEPs displayed the highest batched scores from 1.5-2, whereas from 0 to 1.5, MHB-Control had the highest levels of batched scores. Interestingly, Tilson et al. had the highest levels of batched gene expression for the least highly expressed fetal genes (**Fig. 7I, left**). In the downregulated gene list, fetal HEPs had the highest negative scores for downregulated genes from < -2 to -0.75 (**Fig. 7I, right**). The other three experimental groups had similar levels of negative scores from -0.75 to 0 (**Fig. 7I, right**). Further, these groups had higher scores than fetal HEPs in the upregulated genes in the downregulated gene list (**Fig. 7I, right**), and only minor differences were present between groups. Our analysis suggested that MHB Control was comparable to Fetal HEPs. We also visualized proportional adult HEP gene expression in fetal HEPs and experimental groups (**Fig. 7J**). Overall, fetal HEPs exhibited high scores for upregulated PACNet genes as expected, while MHB-Control was next, and ahead of Tilson et al. in terms of both Up and Moderately Up genes (**Fig. 7J, left**). Similarly, for downregulated fetal genes, MHB-Control was ahead of the other conditions for the smallest number of downregulated genes. Therefore, in terms of transcriptional shaping of hPSC-HBs, MHB Control outperformed other conditions for 3 out of 4 categories (Up, Mod. Up, Down) (**Fig. 7J, left**). For the downregulated PACNet genes, for two categories, the conditions only varied only by nine genes, leaving only two categories. MHB Control had the least number of Moderate-Up genes, and LD-HB2 had the least number of Up genes. These data strongly suggest that MHB Control was highly successful at transcriptional shaping towards Fetal HEPs than LD-HB2 and Tilson et al. populations, despite a large initial discrepancy in PACNet gene scores.

### The migration/growth step is characterized by increased transcription of signaling pathways

We hypothesized that incorporating the growth stage may increase global transcriptional shaping in part by modulating signaling pathways. We first examined murine E9.5 MHB signaling pathways, by comparing to E8.5 GT and E10.5 HB, and we identified 17 candidate receptors and 16 signaling mediators (**Sup.** Fig. 10A), validated with ENRICHR (**Sup. Table 8**). We observed many positively enriched cell stress pathways, including energy (AMPK), oxidative (FOXO), hypoxia (HIF1A), nutrient (mTOR), and DNA damage (p53), which also are listed in our comparative analysis (**Fig. 8A**). Additional signaling pathways included Wnt, TGF-Beta, Hedgehog, Hippo Signaling (Hippo positive and Hippo negative), and the mediators Ras, PI3K and AKT signaling mediators (**Fig. 8A**). In murine HB, blood (blood coagulation, hemostasis positive), and metabolic pathways (Long chain fatty acid metabolic process, PPAR-gamma, HIF1A signaling, Oxidation reduction, TCA cycle, Ox. Phosp.) were upregulated (**Fig. 8A**). A very similar list of signaling and metabolic pathways was obtained when we compared the MHB and MHB Control and the LD-HB1 and LD-HB2 conditions, respectively (**Fig. 8B**). Interestingly, ChIP-seq analysis of TBX3 in human liver (HEPG2) cells demonstrated that Signal Transduction pathways and Metabolism were two of the most commonly TBX3-regulated pathways (**Fig. 8C-D**), and many of the signaling pathways observed correlate with those upregulated in the MHB murine and human populations (**Fig. 8D**). ChIP-seq analysis of FOXA2 and CEBPA also demonstrated their role in cell signaling and metabolism (**Sup.** Fig. 10B-E). Heat map analysis for p53, Hippo, Neural migration, TCA, and Ox. Phosp., confirmed the large differences between MHB and MHB control (**Sup.** Fig. 10F).

**Figure 8.**
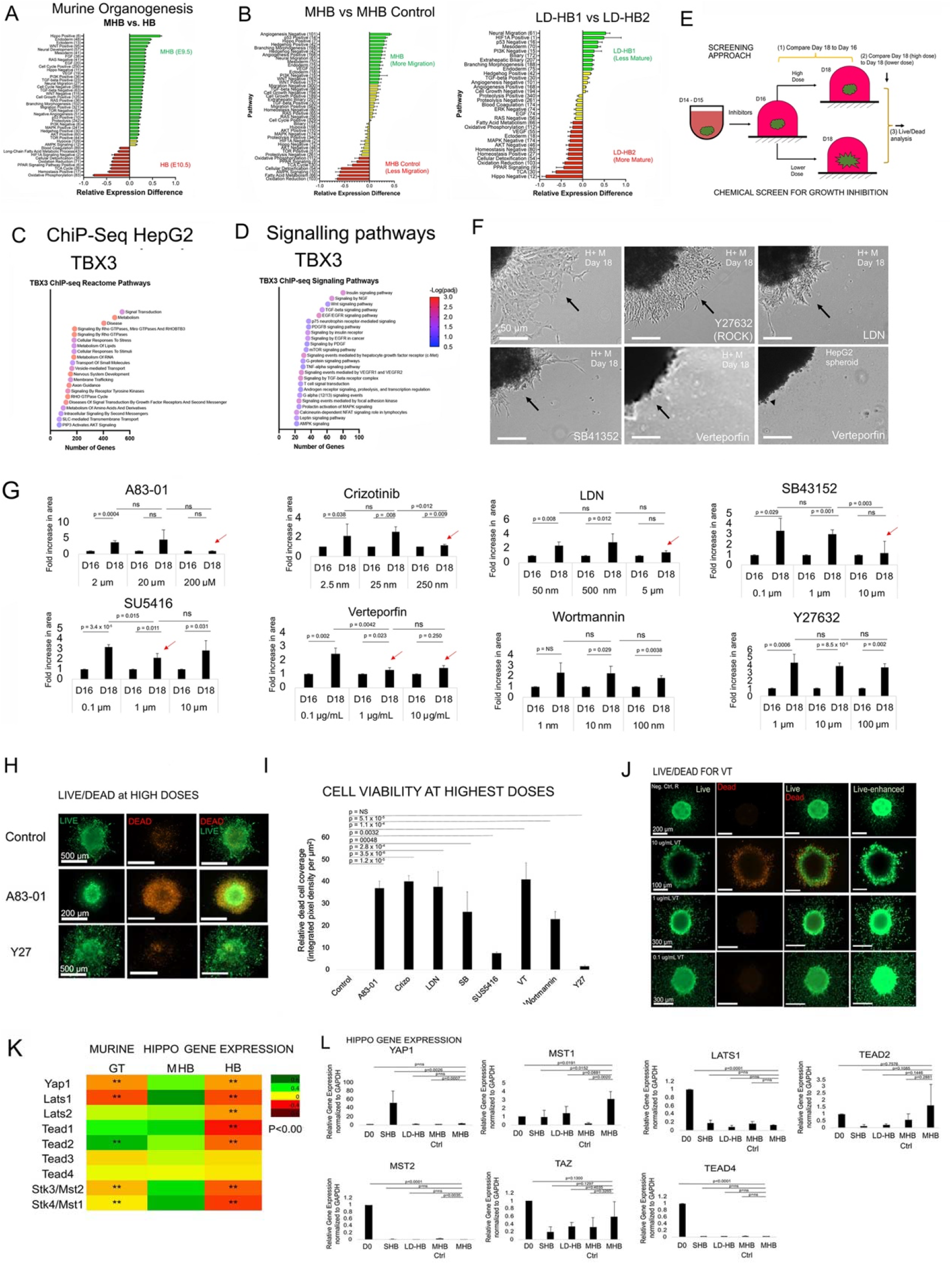
Chemical screen of migrating organoids with cell signaling inhibitors A) Average transformed expression differences for select development, signaling, and metabolic GO pathways between MHB (E9.5) and HB (E10.5) in mouse development. Plotted is mean ± SE. B) Same as A) except comparing our hPSC derived MHB and MHB Control conditions as well as our LD-HB1 and LD-HB2 conditions C) ChIP-seq results for most highly enriched Reactome Pathways D) Same a D) except filtering for significantly enriched (p<0.05) signaling pathways E) Schematic of functional chemical screen of signaling pathways that effect organoid growth/migration. F) Images of treated organoids. Top- untreated control, ROCK treated, LDN treated; Bottom- SB41352 treated, Verteporfin (VT) treated, and HepG2 spheroids + V treated; Arrows show inhibition. G) Data for each inhibitor screened; Fold-increase in area (growth) with 3 concentrations per chemical inhibitor; Red arrows indicate positive hits; P-values listed; n = 3 for each condition. Plotted is mean ± SD. H) Images of live (green) /dead (red) assay for cell viability after chemical treatment. I) Quantitation of experiments in D); P-values shown; N =3 per condition; mean ± SD. J) Same as D but focused on VT treatment; enhanced images (green) shown. K) Heatmap analysis of averaged hippo pathway mediator gene expression from mouse scRNA-seq data (Lotto et al.) ** p < 0.01. L) Hippo gene expression analysis in day 18 organoids; Monolayer (MONO), Suspension (SUSP), Control (CNTRL), MG (Migrating). One-way ANOVA using Tukey’s multiple comparison test; mean ± SD. (2582 Words)

Overall, we present common signaling pathways in all three migrating populations, and connect these signaling pathways with the three member gene circuit described in **Fig. 4**.

### *In vitro* chemical screen of pathway inhibitors demonstrates that Hippo pathway (YAP-TEAD inhibitors) mediates collective migration/growth in day 18 MHBs

We aimed to perform a functional screen to determine which pathways inhibit migration. We used the data to design and perform an *in vitro* chemical screen from day 16 organoid to day 18 migrating organoid, with three criteria to recover potential positive hits (**Fig. 8E**). Initially, we specifically observed significant reduction in growth/collective migration in response to the VT treatment (YAP-TAZ/TEAD inhibitor) when compared to ROCK inhibitor, LDN, and SB41352 treatment (**Fig. 8F**). These results demonstrated feasibility and hence, we expanded our screen to twenty-four inhibitor conditions with eight candidates (three concentrations per candidate) (**Table 3**). We performed dose responses up to three orders of magnitude (**Sup. Table 10**). Our functional screen initially demonstrated a loss of measurable migration/growth in several conditions, including A83-01 (high), Cristozinib (high), LDN (high), SB43152 (high), SU5416 (intermediate), Verteporfin (VT) (high, intermediate) (**Fig. 8G, red arrows**). Interestingly, at intermediate doses, the only inhibitors that significantly reduced migration/growth were VT and SU5416 (**Fig. 8G, red arrows**). We hypothesized that the lack of migration/growth at the highest concentrations was due to a loss of cell viability rather than migration. Indeed, we found that all inhibitors except SU5416 and Y27632 caused cell death at high doses (**Fig. 8H-I, Sup.** Fig. 10G-H). The two remaining inhibition conditions at intermediate doses were SU5416 and VT, which we found decreased migration/growth but did not cause cell death (**Fig. 8J, Sup.** Fig. 10H), and thus were positive hits recovered by the screen. We focused on the effects of VT, since SU546 did not cause inhibition at high doses. Our data for E9.5 MHB, MHB, and LD-HB, all demonstrate transcription of Hippo pathway is involved, and it is known that Hippo signaling controls liver cell fate, regeneration, and growth through complex signaling cascade with several mediators ^59^. We conclude that YAP-TEAD signaling plays a key role in promoting migration in MHB population, and that inhibition of YAP-TEAD with VT inhibits migration. scRNA-seq analysis of Hippo mediators demonstrated upregulation in the MHB population (**Fig. 8K, Sup. Table 4, 7**). We evaluated gene expression of Hippo mediators hPSC-HEPS and the data showed that YAP1 and MST1 was significantly upregulated in MHB and MST2 was significantly downregulated in MHB control (**Fig. 8L**).

**Table 3.** Table of small molecule inhibitors and molecular targets.

| **Small Molecule Pathway Inhibitors** | **Molecular Targets** |
| --- | --- |
| A83-01 | TGF-β type 1 activin receptor like kinase (ALK-5), activin type 1B receptor (ALK4), nodal type 1 receptor (ALK7) |
| LDN-193189 | BMP type 1 receptors: activin receptor like kinase (ALK 1,2,3 and 6) |
| Crizotinib | c-MET receptor |
| SB43152 | TGF-β type 1 activin receptor like kinase (ALK-5) |
| SU5416 | Vascular endothelial growth factor receptor (VEGFR2; FLK1/KDR) |
| Verteporfin | Yes associated protein (YAP) and TEAD interactions |
| Wortmannin | PI3K enzymes |
| Y-27632 | Rho-associated protein kinases (ROCKs) e.g. p160ROCK, ROCK2 |

In summary, our chemical screen recovered VT as a positive hit, and transcription of Hippo mediators in MHB populations suggest collective migration is controlled by YAP-TEAD (**Fig. 8L**). (6339 Words)

## DISCUSSION

Epithelial tissues transition from migration to differentiation in several instances, in organ development, and in wound healing/tissue regeneration. However, hPSC studies generally have not investigated this process or linked it to maturation, while maturation of transcriptionally complex epithelial cells like liver, pancreas, and lung remains a key challenge. We elucidated the missing connection between these two impediments to HEP maturation. We highlighted two missing steps in current hPSC-HEP protocols. Using this concept, we engineer one of the first protocols for generating the elusive human hPSC-MHBs, and we demonstrate these MHBs share common signatures with murine MHB and migrating, regenerating HEPs. We perform simple cultivation changes that lead to striking, global transition from MHBs to HEPs. Our analysis of the efficiency indicates that two hPSC-HEP populations, MHB and LD- HB2, are among the top 5 most efficient protocols. The MHB to HEP transition also leads to improved transcriptional shaping towards an adult HEPs population. Finally, we develop a predictive gene circuit in which three markers predict the transition from MHB to HBs. Because our protocol does not employ additional soluble factors, small molecule inhibitors, it can be potentially integrated into any platform. To accomplish this, we develop several bioinformatic tools that can easily be adopted.

Human epithelial collective migration is essential for epithelial morphogenesis ^60^, and our study should provide cell biologists avenues for investigating collective cell migration, Our study details similarities between MHBs, murine E9.5 MHBs, and regenerating migrating HEPs, and we develop a mapping and analysis approach to distinguish these migrating cells and their phenotype from their mature HEP counterparts. The data suggests that this process was accompanied by not only TBX3+ upregulation, but also increased Hippo, Wnt, TGF-Beta, hypoxia, mTOR, AMPK signaling, and a reduction in Ox. Phos., and TCA cycle. We also demonstrate the MHB’s are characterized by upregulation of mesoderm and neural TF expression, which was not known. To induce MHBs, we performed several years of trial-and-error experiments, and we found that day 14 organoids embedded in Matrigel for 2 days with H+ M medium, initiated collective migration on day 16. Since removal of the H + M coincided with maturation of MHB, the removal of H + M (FGF2, EGF, IGF-1, VEGF and Heparin) results in maturation. We speculate that loss of EGF signaling may reduce FOXA2 ^61^ or loss of VEGF may inhibit hepatic migration ^62^. We also speculate that differentiation enhancers of HEPs employ TBX3 and FOXA to interact with YAP-TEAD to promote signaling in MHB ^63^. Our chemical inhibition studies indeed predict that Hippo pathway signaling (YAP-TEAD) is critical to migration, and TBX3 binds to the TEAD promoter, TBX3 has a complex relationship with Hippo pathway in the liver ^64^. The decrease in oxidative phosphorylation and TCA cycle genes can be explained by the global downregulation of factors like FOXA2 and HNF4A which control metabolic genes during HEP differentiation ^39^. We perturbed FOXA1/2/3 in effort to elucidate gene regulation, but also to determine if could re-create the MHB phenotype. Our data suggests that MHB cells can be generated moving forward, but not in the reverse direction, by targeting FOXA1/2/3. Overall, the induction and characterization of MHBs provides a new avenue for studying epithelial and liver biology, and clinically translatable protocol for maturation or other cell and tissue therapy applications.

PSC-HEP differentiation protocols are over 20 years old, but our study demonstrates that a re-examination of rules and assumptions may be needed. Most protocols include 5-10 additional factors form the GT stage to the HEP stage ^16^, co-cultivation with supporting cells and 7 additional factors ^58^, and programming with 4-8 TFs ^19, 57, 65^. The development of PACNet, ^56^ amongst other tools, enables determination of transcriptional fidelity compared to adult HEP counterparts, and using this tool, we found that regardless of approach, all current studies remain significantly less transcriptionally mature than 12-week Fetal HEPs. We develop a new differentiation approach that may have the capacity to reach Fetal HEPs, that does not require additional soluble factors, TF programming, or co-cultivation. Employing migration to differentiation transition results in two of the five highest normalized PACNet scores. Our data suggests that this results in a downregulation of TBX3, upregulation of liver GRN including CEBPA, increase in metabolism, loss of signaling pathways, all resulting a complete change in cell state. Interestingly, CEBPA functions as TF that induces differentiation in the skin during regeneration ^66–68^. It is important to note that H + M medium does have a single instructive factor, FGF2, and Heparin which may improve its function the cell surface sequestration. Nevertheless, no other known maturating factors were used. Our data suggests that: 1) this transition approach appears to skip several steps in differentiation and, 2) there is tremendous scope to mature MHB control by integrating into other protocols, since only one instructive factor (FGF2) was used, with no additional instructive or maturating factors, small molecules, or programmed TFs. 3) Spontaneous gene activation with minimal GFs may be important to allow cells to coordinate gene modules for differentiation and maturation. Interestingly, the Tilson et. al. study, the highest scoring study, employed continuous hypoxia and only 3 factors (FGF2, and later HGF, Oncostatin) which is as few GFs as was used in any traditional protocol, similar to our study. Overall, we present a new way of maturing cells, which appears to skip several stages in maturation which we term “grasshopper effect.”. We also present transcriptional shaping as a method for determining how far we are towards generating 12-week Fetal HEPs, and the MHB to MHB Control transition presents several advantages in matching Fetal HEPs more than the Tilson et al. study. These data suggest transcriptional activation of Fetal HEP genes can occur via growth to differentiation transition, and that that maturation after transcriptional activation may raise the levels transcriptional maturation. In fact, our data actually implies that changes take place over two days of culture, and if we calculate efficiency based on that, our approach would be far and away the most efficient approach for differentiation.

There are several limitations to our study. Specifically, apart from FGF, we intentionally refrained from introducing conventional instructive factors such as BMP2, and maturation factors like HGF, retinoic acid, dexamethasone, and oncostatin. Although we obtained transcriptionally mature hPSC-HEPs, and we have performed urea and ALB secretion studies, we did not evaluation metabolic functions, and the cells can be matured further.by the addition of maturating factors towards the end of our study, or in additional stage. A second limitation is that we used commercially available medium, EGM-2 (Endothelial growth medium 2 (Lonza)), which has proprietary amounts of FGF2, EGF, VEGF, IGF-1, and Heparin. Future studies will use known concentrations of each of these factors to better understand the role of these individual and combined factors. A third limitation is that we performed conventional RNA-seq instead of scRNA-seq, which would have helped determine whether markers were co-expressed, whether separate lineages were formed, and how lineages changes over time. The advantage of RNA-seq, however, is the improved depth of sequencing that we obtain, particularly for TF expression. However, there is a possibility there are differentiated cell subtypes that are present, potentially impacting our conclusions.

In summary, we were motivated to model liver organogenesis between E8.5-10.5, and based on this data and our analysis, we have developed a highly efficient hPSC-HEP protocol that employs an entirely new technique to differentiate cells, using a migration/growth to differentiation transition. The first new stage involves induction of MHBs, which can be used to understand how human cells collectively migrate and grow to form epithelial tissues. We determine the phenotype of these cells using comparisons to MHB’s in murine development and human regeneration, this leads us to a common TBX3 + high mechanism which is accompanied by increase signaling, increased expression of alternate lineage TFs, and decreased oxidative metabolism. By cultivation change, we observe a rapid change to mature cells which results in loss of signaling, increase metabolism, and activation of liver GRN TF, and this switch results in two of the five most transcriptionally mature cells in the field, based on both transcriptional scores and transcriptional shaping. These studies highlight the potency of this new unrecognized stage in hPSC differentiation. (1353 Words)

## METHODS

For Reagents, Cell Lines, Antibodies, Culture of cell lines (HepG2, HUVEC, HFF), *In vivo* **transplantation assay**

Please see Sup. Methods.

### Feeder-free culture (maintenance), harvesting, and collecting of hPSC

Methods are as described previously ^39^.

### Design of culture system

Since our unique transplantation model supported the hLD-MESC hypothesis, we wanted to develop an *in vitro* system based on this (**Fig. 2A-C**). Our focus on early liver organogenesis precluded the use of traditional instructive/maturating factors like HGF, oncostatin, dexamethasone, and retinoic acid. Further, we noted that sequential instructive factors can provide mixed signals, and thus we chose to eliminate sequential factor design. Instead, we chose a single medium formulation, without any changes, which was employed for the entirety of differentiation under hypoxic conditions. In the first condition, we chose a nutrient rich medium containing, SFD previously used for maintaining hPSC-derived endoderm progenitor cells, the precursor to HBs, with minimal growth factors ^39^. We retained KGF (FGF7) in the medium, since KGF has been shown to induce hPSC-differentiation of endoderm towards GT ^39^. For the second strategy, we used data from ENRICHR pathway analysis to identify that VEGF, EGF, FGF2, and IGF-1 signaling were upregulated in the MH population. Each of these factors has been linked to liver differentiation and growth ^44^,. We found that EGM-2 medium, a commercially available medium employed VEGF, EGF, IGF-1, and FGF (**Fig. 2A-C**), and therefore we employed EGM-2 medium, but removed both hydrocortisone and serum (**Fig. 2A-C**).

### Preparation of SFD medium

Methods are as described previously ^39^.

### Preparation of H + M medium

Base (control) medium contained 50% RPMI, 1% B27, 0.2% Knockout serum (KOSR) (no FBS), and 1% P/S. The modified EGM-2 medium contained EGM-2 basal medium (Lonza), 2% KOSR, 0.5% B-27, 0.05% long R3 insulin growth factor (R3-IGF), 0.05% epidermal growth factor (EGF), 0.2% fibroblast growth factor 2 (FGF2), 0.05% vascular endothelial growth factor (VEGF), 0.05% ascorbic acid, 0.05% heparin, 0.05% gentamycin, 1% P/S. These were mixed at 50:50 conditions.

### DE induction from human stem cells under hypoxic (5% O_2_) conditions

Methods are as described previously ^39^.

### For HB induction using growth factor GF (+) differentiation protocol, and HB cell induction using GF (-) containing SFD medium

Please see Sup. Methods.

### HB cell induction using hepatic and mesenchymal (H + M) medium

hPSC-derived DE were differentiated towards HBs from day 4 to day 14 using H + M medium (defined above), under hypoxic conditions (5% O_2_). Please see Sup. Methods for further details.

### Fluorescent dye-labeling of stem cell-derived progenitors

Please see Sup. Methods.

### Live/dead assay for thin cell sheet and liver organoids

Organoids were assayed for cell viability using a live/dead reagent kit (Biotium, Catalog #30002-T). Please see Sup. Methods.

### GT organoid formation, including GT endoderm organoid formation, GT endoderm/HUVEC organoid formation, GT organoids in EGM-2 medium

Please see Sup. Methods.

### Generation of microfabricated device (micropillar device) and Generation of thin organoids (microtissues) in device

Please see Sup. Methods.

.Preparation of agarose-coated microwells

Methods are as described previously ^39^.

### Collagen type I gel formation, Organoid culture in MG droplets and Organoid culture in collagen droplets

Please see Sup. Methods.

### Harvesting spheroids embedded in MG or collagen

Spheroids embedded in gel were harvested and placed in 15 mL tubes. Accutase was then added to the tubes. Please see Sup. Methods.

### Histology assay, Histological analysis of organoid and tissues, Microscopy of organoids, Immunofluorescence microscopy, Whole organoid fixation and immunofluorescence

Please see Sup. Methods.

### Urea assay

Urea metabolite concentration analysis was done via a spectrophotometric urea assay kit (Bioassay systems, Catalog No: DIUR-100). Please see Sup. Methods.

### ELISA assay

Please see Sup. Methods.

### Reverse transcriptase reaction and real-time polymerase chain reaction (RT-PCR)

Please see Sup. Methods.

### Analysis of organoid migration

ImageJ was used to identify relative characteristics of the migrating organoids. Please see Sup. Methods.

### Filtering of lateral 3D migration for improved image visualization

Please see Sup. Methods.

### siRNA and shRNA knockdown of day 12 H + M treated hPSC-HEP cells

Day 12 H + M cells were treated with shRNA FOXA1/2 and siFOXA3 using vectors, lentiviral production, transfection reagents and procedures described recently. ^39^ Please see Sup. Methods.

### Chemical screening of growth inhibition in MG culture

Eight small molecule inhibitors, with three conditions each, for a total of 24 conditions were employed (**Sup. Table 1**). As mentioned above, D15 H + M (20 organoids, 300-400 µm in diameter) were collected from suspension culture and seeded onto 60 mm tissue culture treated dishes as done in the MG organoid droplet culture (one organoid per droplet, 20 droplets). After the MG gelling step (droplets were incubated at 37°C, 5% CO_2_, 5% O_2_ for 30 minutes at to allow the MG to solidify) the organoid droplets were cultured in 5 mL of H + M media supplemented with a pre-determined concentration of a small molecule inhibitor. Please see Sup. Methods.

### Live/dead assay for biologically inhibited organoids in MG culture

Please see Sup. Methods.

### Statistics

For statistical assessment between two groups, Student’s t-test was done (type II with two tails), and p-values equal to or less than 0.05 was considered to have statistical significance and are defined within the text and figures/figure legends. ANOVA was employed for comparison of multiple groups.

### RNA-sequencing (RNA-seq). and Bulk RNA-seq Analysis

Please see Sup. Methods.

### scRNA-seq Data Collection, Normalization and Filtering

Please see Sup. Methods.

### snRNA-seq Analysis

Data from 72,262 human nuclei was analyzed with snRNA-seq for the liver regeneration data from healthy samples (n = 9), Acetaminophen (APAP) -Acute liver failure (ALF) samples (n = 10), and Non-A, B, C, D, E (NAE)-ALF samples (n = 12). Please see Sup. Methods for further details.

#### Cell Cluster Regrouping

Please see Sup. Methods for further details.

#### T-distributed stochastic neighbor embedding (TSNE)

Please see Sup. Methods for further details.

#### Pathway Heatmaps

Please see Sup. Methods for further details.

#### Pathway validation

To validate pathways in the MH cluster, we examined the expression of five liver differentiation genes (ALB, AFP, HEX, PROX1, TBX3), and these correlated in all three databases (DAVID, REACTOME, ENRICHR).

*Differential Expression Analysis-DAVID, Differential Expression-ENRICHR, Comparison of DAVID and ENRICHR, scRNA-seq Gene expression PCA plots, Developmental Mouse Pathway Ranking* Please see Sup. Methods for further details.

### Average Pathway Score

Average pathway scores were determined by running the ScaleData function with a negative binomial model to transform the expression values for all the genes within the dataset. Please see Sup. Methods for further details.

### Pie Chart analysis

The FindMarkers function in Seurat was used with scRNA-seq data to find differentially expressed genes (log2fc>0.5,padj<0.01) between different cell types. For bulk RNA-seq, the results function in DESeq2 was used to find differentially expressed genes (log2fc > 1.5,padj < 0.05). Please see Sup. Methods for further details.

### PACNet Classification

The PACNet Classification Score was first run using the CellNet PACNET web client (http://ec2-3-89-75-200.compute-1.amazonaws.com/cl_apps/agnosticCellNet_web/), using normalized gene expression data for our samples from DESeq2. Please see Sup. Methods for further details.

### Pvalue Adjustment and Multiple Comparison Test

To adjust pvalues between PACNet classification scores we calculated the pvalues for all comparisons and adjusted the pvalues utilizing the Bonferoni adjustment with an alpha equal to 0.05.

### Alternative PACNet GRN Analysis

Since PACNet only utilizes 100 gene pairs to classify derived hepatocytes, we desired to further analyze all the genes within the PACNet GRN list. We went around accomplishing this by creating a combined dataset of samples with our conditions (n = 17), gut tube (n = 3), 12 week fetal liver (n = 2), as well as from several different derived hepatocyte protocols including Li et al. (n = 5), Velazquez et al. (n = 10), and Tilson et al. (n = 4). Please see Sup. Methods for further details.

### Transcription Factor (TF) Correlation Analysis

Please see Sup. Methods.

### Deep Learning Model

For our analysis, we selected specific transcription factors (Foxa2, Foxq3, Hnf1a, Cebpa, Tbx3, Prox1, Hnf4a) from the murine scRNA-seq data as predictors, aiming to predict the RNAseq time points (mentioned earlier) variable. Please see Sup. Methods for further details. (1322 Words)

### Disclosure, Intellectual Property

NP has received a formal waiver R-030-7485 from UB for this work, and the work is currently submitted by Khufu Therapeutics to US Patent office US2370292. NP is founder of Khufu Therapeutics and CG is a consultant for Khufu Therapeutics.

## Acknowledgements

NP was supported by the UB CBE startup funds, a Mark Diamond Fellowship, the New York State Stem Cell Science C024316, the UB the Stem cells in regenerative medicine (ScIRM) center, the University at Buffalo Center for Cell, Gene, Tissue Engineering (CGTE). OO was supported by the Western NY Prosperity Fellowship. We would like to thank Drs. Jeremy Lotto and Pamela Hoodless (University of British Columbia), for their work regarding the publication of their manuscript and associated data which was used for the murine liver organogenesis bioinformatics figure. We would like to thank Neil Henderson PhD, FRCP, FAASLD, FMedSci, Centre for Inflammation Research, Queen’s Medical Research Institute, University of Edinburgh, UK. We would like to thank Dr. Imtiaz Mohammed (Lead histologist) and Dr. Amber Worral (PAS Microscopy and Histology Core Labs Director) for their assistance with tissue preparation, processing and sectioning of our tissue samples within the histology core facility at the University at Buffalo. We would also like to thank Dr. Wade Sigurdson (Confocal Microscope and Flow Cytometry Facility Director) for his advice on methods of imaging tissues and whole-mount organoids.

## AUTHOR CONTRIBUTIONS

**OO, DG, ST, AC, TN, DB, CO, AK, CS, PC, SR, ZC, PJ, SM, RZ, RG, NP**: Supervising the work, Responsible for all data, figures, and text, Ensuring the authorship is granted appropriately, Ensuring that the authors approve the content and submission of the paper, Ensuring that all authors approve the content and submission of the paper, as well as edits made through the revision and production processes, Ensuring adherence to all editorial and submission policies, Identifying and declaring competing interests on behalf of all authors, Identifying and disclosing related work by any co-authors under consideration elsewhere, Archiving unprocessed data and ensuring that figures accurately present the original data. **NP**: Arbitrating decisions and disputes and ensuring communication with the journal (before and after publication), sharing of any relevant information or updates to co-authors, and accountability for fulfillment of requests for reagents and resources.

## ABBREVIATIONS

AFP: alpha-fetoprotein
ALB: albumin
BMP4: Bone Morphogenetic Protein
BSA: Bovine serum albumin
CCM: Collective cell migration
CHIR: Wnt pathway agonist
DMEM: Dulbecco’s Modified Eagle’s Medium
EDTA: Ethylenediaminetetraacetic acid
EG: Epidermal growth factor
EHT: epithelial to hepatic transition
FBS: fetal bovine serum
FGF2: Fibroblast growth factor-2 GT Gut Tube
iPSC: Induced pluripotent stem cells
HEP: Hepatocyte
H + M: hepatic and mesenchymal
hESC: human embryonic stem cells
hPSC: human pluripotent stem cells
HBs: hepatoblasts
HSCs: hematopoietic stem cells
HE: hepatic endoderm
iPSC: induced pluripotent stem cells
LD: liver diverticulum
LD-HB: liver diverticulum hepatoblasts
MES: Mesoderm
MG: Matrigel (growth-factor free)
MHB: Migrating Hepatoblast
MHB: Control Migrating Hepatoblast exposed to Control medium
RT-PCR: Real-time polymerase chain reaction
R3 IGF-1 R: Insulin growth factor-1
SFD: Serum free-differentiation
STM: Septum transversum mesenchyme
TF: Transcription factor
VEGF: Vascular endothelial growth factor
VT: Verteporfin
YAP: Yes associated Protein

## AUTHOR CONTRIBUTIONS

OO, DG, ST, AC, TN, DB, CO, AK, CS, PC, SR, ZC, PJ, CG, SM, RZ, RG, NP: Supervising the work, Responsible for all data, figures, and text, Ensuring the authorship is granted appropriately, Ensuring that the authors approve the content and submission of the paper, Ensuring that all authors approve the content and submission of the paper, as well as edits made through the revision and production processes, Ensuring adherence to all editorial and submission policies, Identifying and declaring competing interests on behalf of all authors, Identifying and disclosing related work by any co-authors under consideration elsewhere, Archiving unprocessed data and ensuring that figures accurately present the original data. NP: Arbitrating decisions and disputes and ensuring communication with the journal (before and after publication), sharing of any relevant information or updates to co-authors, and accountability for fulfillment of requests for reagents and resources.

