## Supplemental Figures, Tables, and Methods for "Linking collective migration/growth to differentiation boosts global shaping of the transcriptome and exhibits a grasshopper effect for driving maturation"

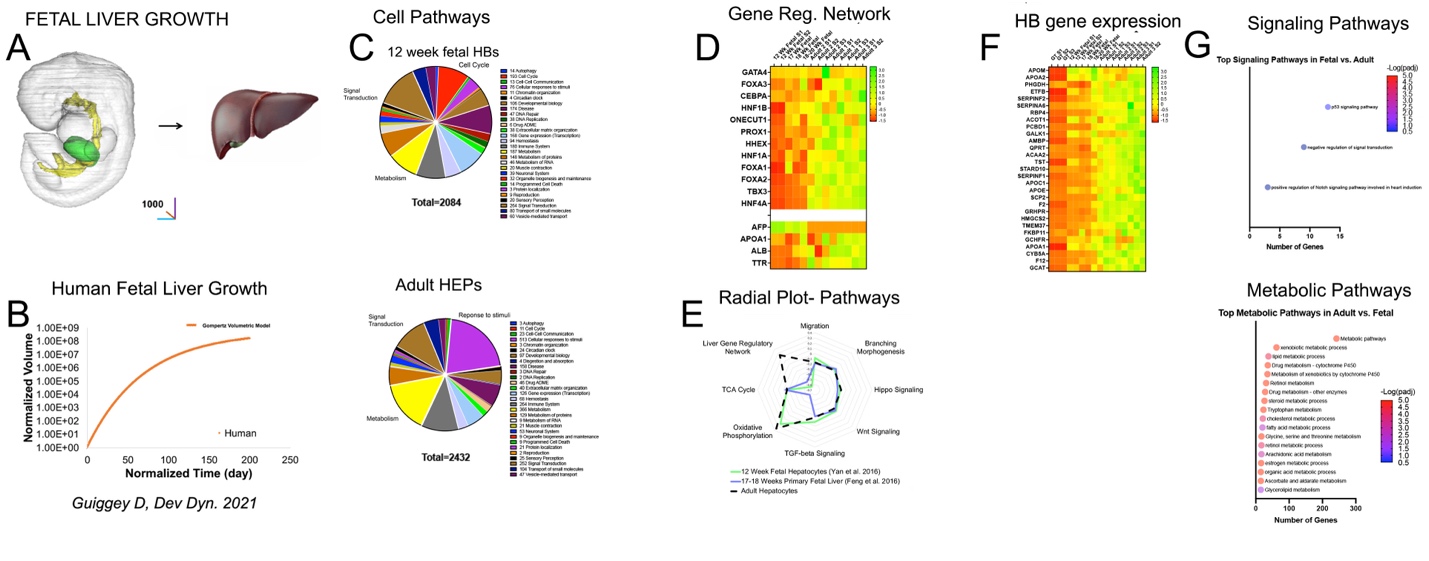

**Supplementary Figure 1.**

A) Diagram of human fetal liver (Ogoke, Yousef et al.) transitioning to adult fetal liver by proliferation.
B) Gompertz model for human volumetric growth normalized to Day 7 embryonic liver (Ogoke, Yousef et al.)
C) Pie chart comparison for differentially expressed genes (log2fc > 1.5, padj < 0.05) between 12 week fetal liver (n = 2) and Adult (n = 3) using Reactome Pathway categories to sort genes.
D) Heatmap comparing the average transformed expression values for human fetal liver (Yan et al. (12 week, n = 2), Feng et al. 2016 (17-18 week, n = 2), Ehrlich et al. (18-20 week, n = 2)) and human adult hepatocytes (University of Cambridge (n=2), Du et al. 2014 (n = 3), Boon et al. 2020 (n = 2)) for liver GRNS TFs in liver organogenesis (upper) and maturation genes (lower).
E) Radar plot comparing the average transformed expression scores between human fetal 12 week liver (n = 2), 17-18 week fetal liver (Feng et al. 2016, n = 2), and human adult hepatocytes (n = 7) for select GO pathways.
F) Heatmap comparing the average transformed expression values for in vivo human hepatic lineage samples (Yan et al. (12 week, n = 2), Feng et al. 2016 (17-18 week, n = 2), Ehrlich et al. (18-20 week, n = 2)) and human adult hepatocytes (University of Cambridge (n = 2), Du et al. 2014 (n = 3), Boon et al. 2020 (n = 2)) for top 30 PACNet liver GRN genes that best differentiate mouse MHB (E9.5) and mouse HB (E10.5) shown in Figure 1E.
G) Most highly enriched KEGG signaling and metabolic pathways between 12 week fetal liver (n = 2) and adult hepatocytes (n = 7).

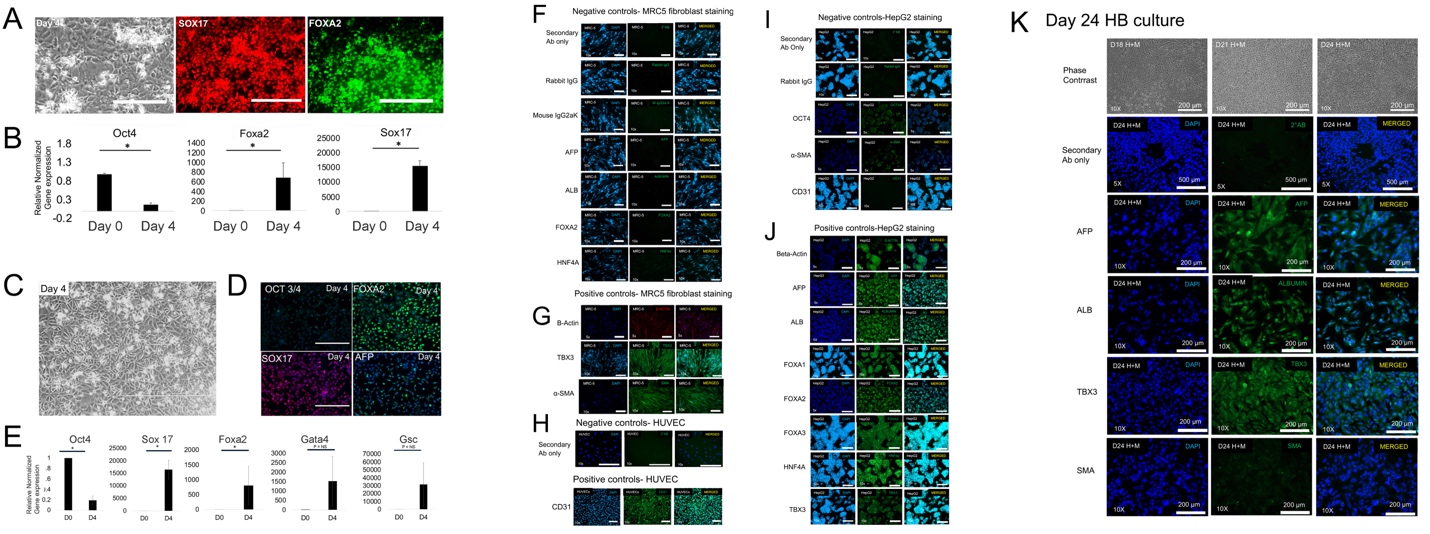

**Supplementary Figure 2. Characterization of hPSC-HB culture.**

A) Morphology and immunochemistry of endoderm and hepatic progenitor cells. Left panel- Phase contrast image of day 4 endoderm morphology. Bar = 200 µm. Middle panels- Sox-17 (red) and Foxa2 (green) staining of endoderm progenitor cells on day 4. Observable protein expression detected.
B) Bar graph of gene expression kinetics (qRT-PCR) of endoderm induction from hPSC. n = 3 for each group compared. All comparisons between day 0 and day 4. OCT 4, P = 0.000067, FOXA2, P = 0.037, SOX 17, P = 0.00029. Plotted is mean ± SD. Significance (*) defined as P ≤ 0.05.
C) Phase contrast images of cells during endoderm induction under hypoxic conditions. Day 4 cell shown.
D) Immunofluorescent staining. Top row - day 4 cells immunestained for OCT4/ DAPI (left) and FOXA2/ DAPI (right), Bar = 200 µm. Bottom -day 4 cells immunostained for SOX17/DAPI (left), and AFP/ DAPI (right), Bar = 200 µm. Observable protein expression detected.
E) Bar graph of gene expression kinetics (qRT-PCR) of endoderm transcription factors (TFs) during endoderm induction from human stem cells, on day 0 and day 4 of culture. Cells transfected on day 1 with siRNA. In control (shScramble) and siFoxa1/2 conditions. N = 3 for control and siFOXA1/2 for all conditions. OCT4, P =0.000063, SOX17, P = 0.0015, FOXA2, P = 0.012, GATA4, P = NS (P = 0.011), GSC, P = NS. Plotted is mean ± SD. Significance (*) defined as P ≤ 0.05.
F) Immunostaining of MRC-5 fibroblast cells cultured in monolayer using same methods described above for negative control targets specifically: 2Ab°, Rabbit IgG, Mouse IgG2aK, AFP, ALB, FOXA2, and HNF4α. Immunostaining of MRC-5 fibroblast cells using Secondary Antibody (goat anti-rabbit) alone. Images are taken at 10x. From left to right: left (DAPI), middle (2 Ab°), right (Merged images). No observable protein expression detected in cells. Scale bar = 200 µm. Immunostaining of MRC-5 fibroblast cells using Rabbit IgG. Images are taken at 10x. From left to right: left (DAPI), middle (Rabbit IgG), right (Merged images). No observable protein expression detected in cells. Scale bar = 200 µm. Immunostaining of MRC-5 fibroblast cells using Mouse IG2a Kappa. Images are taken at 10x. From left to right: left (DAPI), middle (Mouse IgG2aK), right (Merged images). No observable protein expression detected in cells. Scale bar = 200 µm. Immunostaining of MRC-5 fibroblast cells for alpha-fetoprotein (AFP). Images are taken at 10x. From left to right: left (DAPI), middle (AFP), right (Merged images). No observable expression of protein detected in cells. Scale bar = 200 µm. Immunostaining of MRC-5 fibroblast cells for albumin (ALB). Images are taken at 10x. From left to right: left (DAPI), middle (ALB), right (Merged images). No observable expression of protein detected in cells. Scale bar = 200 µm. Immunostaining of MRC-5 fibroblast cells for Forkhead Box A2 (FOXA2). Images are taken at 10x. From left to right: left (DAPI), middle (FOXA2), right (Merged images). No observable expression of protein detected in cells. Scale bar = 200 µm. Immunostaining of MRC-5 fibroblast cells for hepatocyte nuclear factor 4 alpha (HNF4A). Images are taken at 10x. From left to right: left (DAPI), middle (HNF4A), right (Merged images). No observable expression of protein detected in cells. Scale bar = 200 µm.
G) Immunostaining of MRC-5 fibroblast cells cultured in monolayer using same methods described above for positive control targets specifically: β-actin, TBX3, and α-SMA. Immunostaining of MRC-5 fibroblast cells for β-actin. Images are taken at 5x. From left to right: left (DAPI), middle (β-Actin), right (merged images). Observable expression of protein detected in cells. Scale bar = 500 µm. Immunostaining of MRC-5 fibroblast cells for T-box transcription factor 3 (TBX3). Images are taken at 10x. From left to right: left (DAPI), middle (TBX3), right (Merged images). Observable expression of protein detected in cells. Scale bar = 200 µm. Immunostaining of MRC-5 fibroblasts cells for α-SMA. Images are taken at 10x. From left to right: left (DAPI), middle (α-SMA), right (Merged images). Observable expression of protein detected in cells.
H) Immunostaining of HUVEC cells cultured in monolayer using same methods described above for negative and positive controls specifically: 2Ab° and CD31. Negative immunostaining of Human umbilical vein endothelial cells (HUVEC) for platelet endothelial cell adhesion molecule (CD31) using secondary antibody only. Images are taken at 10x. From left to right: left (DAPI), middle (2 Ab° only), right (Merged images). Positive immunostaining of HUVEC cells for CD31. From left to right: left (DAPI), middle (CD31), right (Merged images). Observable expression of protein detected in cells.
I) Immunostaining of HepG2 hepatoblastoma cells cultured in monolayer using same methods described above for negative control targets specifically: 2Ab only, Rabbit IgG, OCT3/4, α-SMA, and CD31. Immunostaining of HepG2 hepatic carcinoma epithelial cells using secondary antibody (goat anti-rabbit) alone. Images are taken at 10x. From left to right: left (DAPI), middle (2 Ab°), right (Merged images). No observable expression of protein detected. Scale bar = 200 µm. Immunostaining of HepG2 hepatic carcinoma epithelial cells for Rabbit IgG. From left to right: left (DAPI), middle (Rabbit IgG), right (Merged images). No observable expression of protein detected. Scale bar = 200 µm. Immunostaining of HepG2 hepatic carcinoma epithelial cells for octamer-binding transcription factor 3/4 (OCT3/4). Images are taken at 5x. From left to right: left (DAPI), middle (OCT3/4), right (Merged images). Low observable protein expression detected in cells. Scale bar = 200 µm. Immunostaining of HepG2 hepatoblastoma cells for alpha smooth muscle actin (α-SMA). Images are taken at 5x. From left to right: left (DAPI), middle (α-SMA), right (Merged images). Low observable protein expression detected in cells. Immunostaining of HepG2 hepatic carcinoma cells for (CD31). Images are taken at 10x. From left to right: left (DAPI), middle (CD31), right (Merged images). No observable protein expression detected in cells. Scale bar = 200 µm.
J) Immunostaining of HepG2 hepatoblastoma cells cultured in monolayer using same methods described above for positive control targets specifically: β-actin, AFP, ALB, FOXA1, FOXA2, FOXA3, HNF4α, TBX3. Immunostaining of HepG2 for Beta actin (β-Actin). Images are taken at 5x. From left to right: left (DAPI), middle (β-Actin), right (Merged images). Immunostaining of HepG2 cells for alpha-fetoprotein (AFP), left (DAPI), middle (AFP), right (Merged images). Images are taken at 5x. Observable protein expression detected in cells. Scale bar = 500 µm. Immunostaining of HepG2 for albumin (Alb), left (DAPI), middle (Alb), right (Merged images). Images are taken at 5x. Observable protein expression detected in cells. Scale bar = 500 µm. Immunostaining of HepG2 hepatic cells for Forkhead Box A1 (FOXA1). Images are taken at 10x. From left to right: left (DAPI), middle (FOXA1), right (Merged images). Observable protein expression detected in cells. Scale bar = 200 µm. Immunostaining of HepG2 cells for Forkhead Box A2 (FOXA2). Images are taken at 10x. From left to right: left (DAPI), middle (FOXA2), right (Merged images). Immunostaining of HepG2 cells for Forkhead Box A3 (FOXA3). Images are taken at 10x. From left to right: left (DAPI), middle (FOXA3), right (Merged images). Observable protein expression detected in cells. Scale bar = 200 µm. Immunostaining of HepG2 cells for hepatocyte nuclear factor 4 alpha (HNF4α). Images are taken at 10x. From left to right: left (DAPI), middle (HNF4α), right (Merged images). Observable protein expression detected in cells. Scale bar = 200 µm.
K) Phase contrast morphology of replated D14 H+M derived hepatoblast cells for an additional ten days, to day 24. From left to right: Left (D18 H+M), middle (D21 H+M), right (D24 H+M). Immunostaining of iPSC D24 H+M derived hepatoblast (HB) cells cultured in monolayer using same methods described above for positive and negative targets: 2Ab°(neg), AFP(pos), ALB (pos), TBX3 (pos) and α-SMA (neg). Immunostaining of D14 H+M cells for 2Ab only. Images are taken at 10x. From left to right: Left (DAPI), middle (2Ab only), right (Merged images). No observable protein detected. Scale bar = 200 µm. Immunostaining of D14 H+M cells for AFP. Images are taken at 10x. From left to right: Left (DAPI), middle (AFP), right (Merged images). Observable protein detected in cells. Scale bar = 200 µm. Immunostaining of D14 H+M cells for Albumin. Images are taken at 10x. From left to right: Left (DAPI), middle (Alb), right (Merged images). Observable protein detected in cells. Scale bar = 200 µm. Immunostaining of D14 H+M cells for TBX3. Images are taken at 10x. From left to right: Left (DAPI), middle (TBX3), right (Merged images). Observable protein detected in cells. Scale bar = 200 µm. Immunostaining of D14 H+M cells for α-SMA. Images are taken at 10x. From left to right: Left (DAPI), middle (α-SMA), right (Merged images). No observable protein detected in cells. Scale bar = 200 µm.

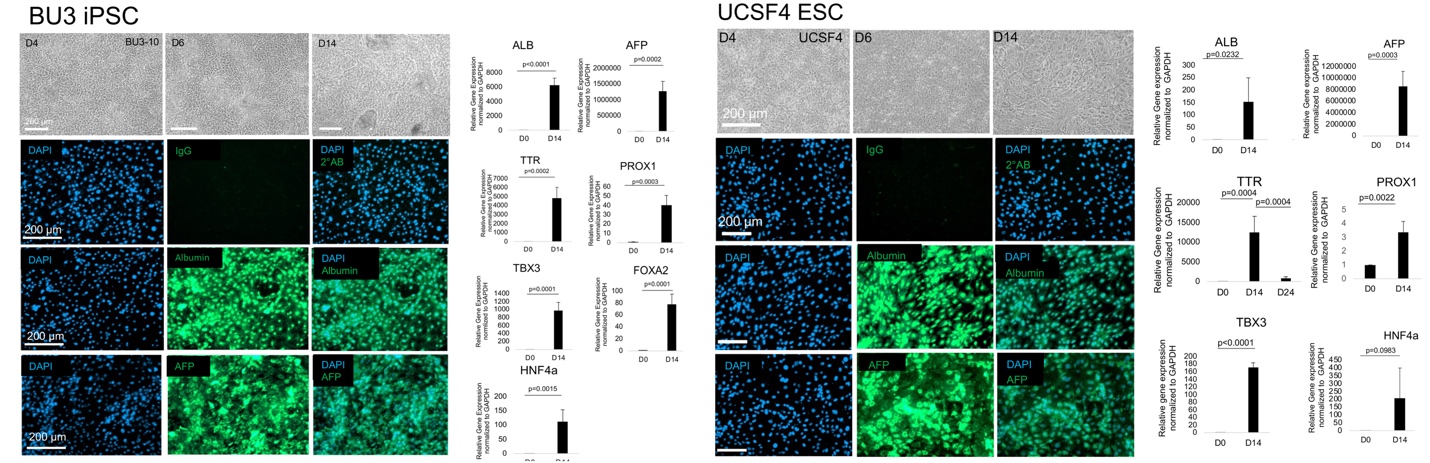

**Supplementary Figure 3.**

A) Phase contrast image for day 4, day 6, and day 14 differentiation using BU3 iPSC. Immunocytochemistry of day 14 BU3 iPSC. DAPI (UV filter) and staining of IgG (top), Albumin (middle), AFP (bottom). Bar graph of gene expression kinetics (qRT-PCR) of day 0 and day 14 for BU3 iPSC liver differentiation. N = 3 for each group compared. Plotted is mean ± SD.
B) Same as A) except for UCSF4 ESC differentiation.

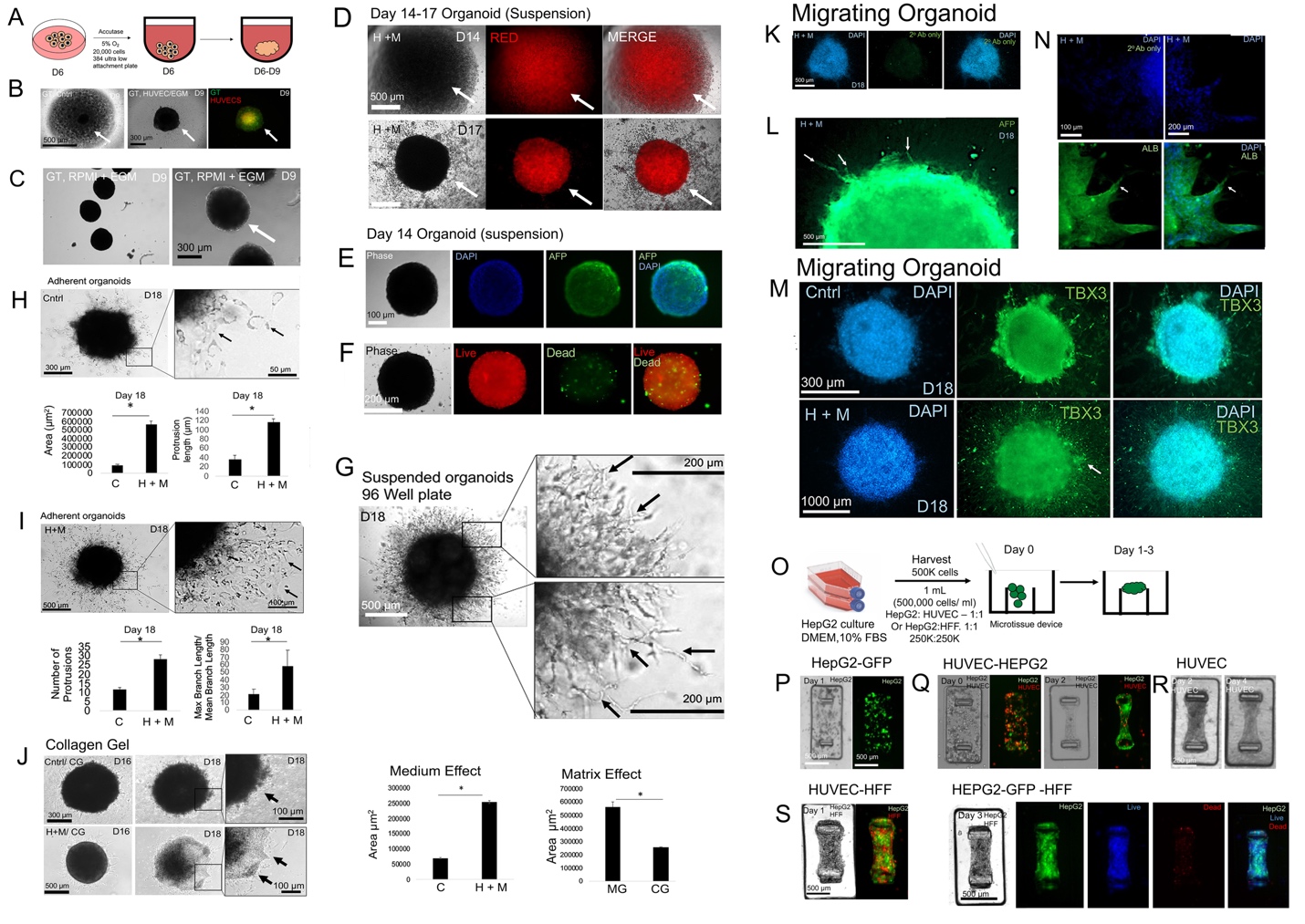

**Supplementary Figure 4.**

A) Schematic of 3D hPSC-derived organoid formation in 384 well ultra-low attachment plates using gut tube (GT) endoderm cells harvested on day 6.
B) Phase contrast image of showing medium effects upon organoid compaction/condensation of hPSC-derived GT endodermal cells. Left- Day 7 GT cells in basal medium, Middle-Day 9 organoids with 1:1 mixture of GT and HUVEC cells in 50% basal/50% EGM-2 medium Right- same as Middle panel except fluorescent image. GT- green, HUVEC- red.
C) Phase contrast image of showing medium effects upon organoid compaction/condensation of hPSC-derived GT endodermal cells. Left- Day 9 GT grown only in 50% basal/50% EGM-2 medium. Right- same as Left panel except higher magnification. No HUVEC are added.
D) Phase contrast images of day 14 and day 17 H + M treated, dye-labeled (red) cells during hPSC-HB organoid formation in 384-well ultra-low attachment plates. Day 14 cells at starting point (above). Day 17 after compaction (below). Organoids uniformly condense to form compact organoids.
E) Phase contrast and immunofluorescence staining of AFP on day 17 hPSC-HB organoids. Whole organoids were fixed and immunostained, with DAPI counterstaining. Cells are AFP positive throughout
F) Live/dead images analysis on day 15 of H + M treated hPSC-HB organoid. Data shows minimal cell death.
G) Phase contrast images of day 18 migrating hPSC-HBs treated in control and H + M medium. Control: cyst like structures (arrow), minimal CCM (arrowhead). H + M organoids (right) demonstrate CCM.
H) Phase contrast images of control (arrow: minimal CCM) and H + M treated (arrow: CCM) hPSC-HB an adherent organoid model. Bar graphs analysis of images in I-J. Area (P = 3 x 10^-4^), protrusion length (P = 2.8 x 10^-4^), number of protrusions (P = 1.4 x 10^-3^), and the max /mean branch length ratio (P = 1.4 x 10^-2^), mean ± SD.
I) Same as H except collagen gel (CG); c images of control (arrow: minimal CCM) and H + M treated (arrow: CCM)
J) Bar graphs analysis in adherent CG droplets. Left: Effects of medium for CG; Right- Effects of MG vs CG for H + M treated.
K) Immunocytochemistry of H + M treated day 18 whole organoids for 2°Ab; counterstained with DAPI and FITC.
L) Immunocytochemistry of H + M treated day 18 whole organoids in MG droplet culture for AFP. Cells were counterstained with DAPI and FITC channel was used. High magnification images taken of migrating cells.
M) Immunocytochemistry of Control (top) and H + M treated (lower) day 18 whole organoids in MG droplet culture for TBX3. Cells were counterstained with DAPI and FITC channel was used.
N) Same as L except for ALB.
O) Schematic of tissue self-assembly in microtissue array format. hPSC-HB, H + M treated day 14 monolayer cells are seeded onto PDMS array posts in collagen hydrogel for 3 days in H + M medium.
P) Phase contrast images of (left) and fluorescent (right) of HepG2-GFP cells (500 cells per microwell) on day 1 after seeding. No microtissue formed.
Q) Phase contrast images (left) and fluorescent images (right) of HepG2-GFP and dye-labeled HUVEC cells seeded in microdevices at seeding of cells (day 0, left) and after two days (right).
R) Phase contrast images of microtissue formed with HUVEC cells, on day 2 (left ) and day 4 (right).
S) Phase contrast images (left) and fluorescent images (right) of day 1 microtissues of HepG2-GFP and HFF (left) and day 3 microtissues (right). Live (blue) and dead (red) staining performed. Live, dead, and merged image shown.

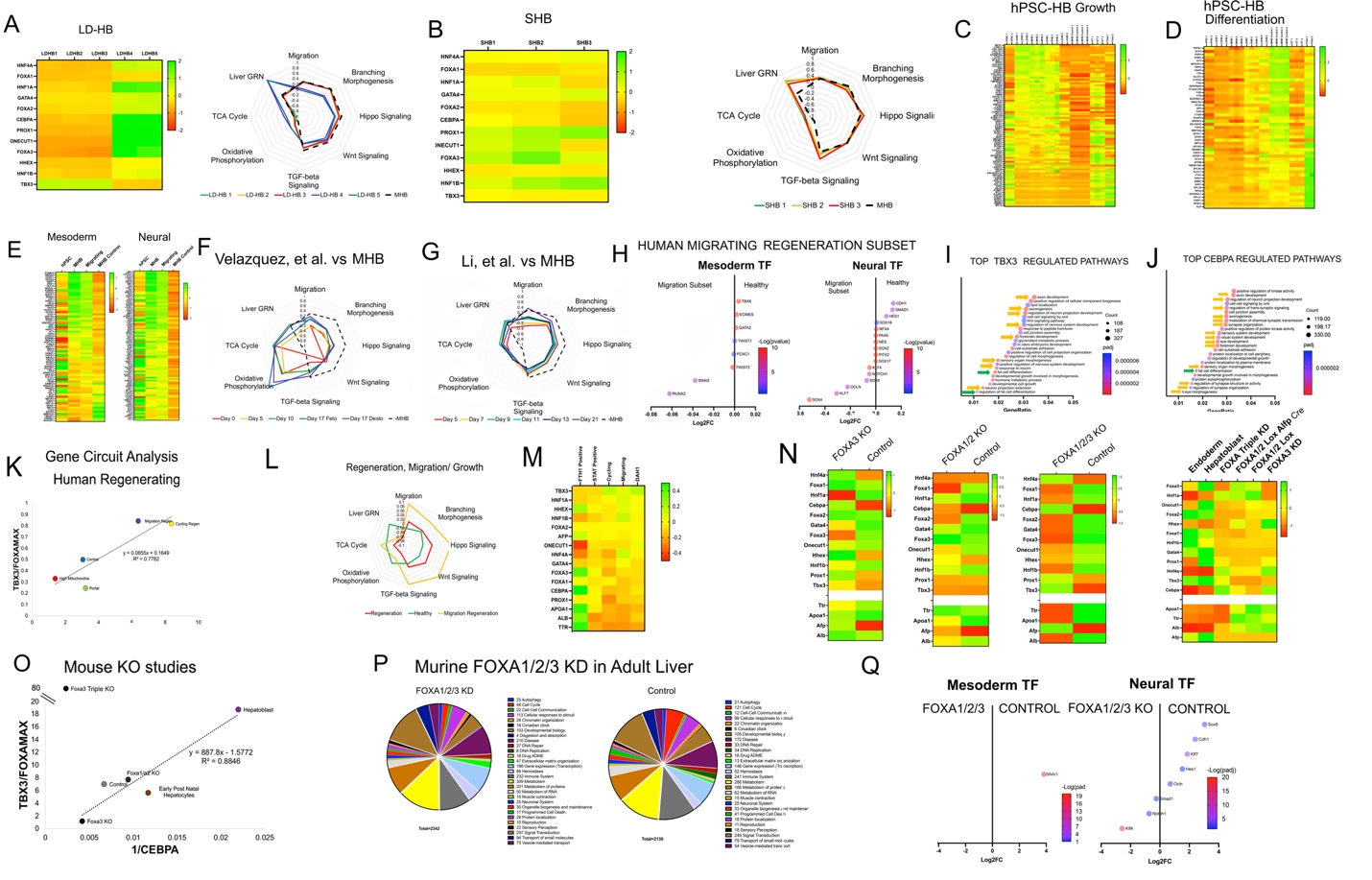

**Supplementary Figure 5.**

A) Heatmap comparing the average transformed expression values in our LD-HB samples (n = 5), and for select important TFs in liver organogenesis alongside a radar plot comparing the transformed expression scores between LD-HB samples for select GO pathways based on the differential expression analysis.
B) Same as A) except for SHB samples.
C) Heatmap comparing the transformed expression values for all our analyzed samples (hPSC (n = 2), LD-HB (n = 5), SHB (n = 3), MHB (n = 3), MHB Control (n = 4)) as well as gut tube and 12 week fetal hepatocyte controls for “developing or growth” gene list based on the overlap significantly upregulated genes (log2fc > 0.5, padj < 0.05) between E9.5 mouse MHB and E10.5 MHB, combined with genes up in MHB and down in MHB Control differential expression (log2fc >1.5, padj < 0.05).
D) Same as C) except for looking for “differentiation” gene list which are the genes downregulated in E9.5 MHB compared to E10.5 HB combined with genes in hPSC-derived MHB control and down in hPSC derived MHB.
E) Heatmap of transformed expression values for top 30 differential expressing mesoderm development genes (GO:0007498) and neuron development genes (GO:0048666) between MHB (n = 3) from MHB Control (n = 4). Control hPSC (n = 2) and migrating condition (n = 7).
F) Radar plot comparing the average transformed expression scores between Velazquez et al. time course samples for select GO pathways based on the differential expression analysis.
G) Same as F) except for Li et al.
H) Gene expression comparisons between ANXA2+ migrating subset cell population in regeneration to cell populations in healthy liver for select mesoderm and neuroectoderm TFs.
I) ChIP-seq gene pathway enrichment analysis for TBX3. Neural (yellow) and mesoderm (green) related pathways are labeled.
J) Same as I) except for CEBPA.
K) Correlation analysis for the ratio between TBX3 and largest normalized FOXA gene (FOXA1, FOXA2, FOXA3) with the inverse of CEBPA. Average gene expression values were calculated for all cells within each cluster population based on clustering done my Matchett et al. (High Mitochondria, Portal, Central, ANXA2+ Migration (Found overwhelmingly only in regeneration condition), Cycling (Found overwhelmingly only in regeneration condition).
L) Same as F) except based on average scores for clusters of regeneration and healthy liver cells based on the analysis by Matchett et al. Regeneration cells included cells in clustering only found in regeneration population while healthy hepatocytes included cells clustering in both regenerating and healthy liver tissue. ANXA2+ migration cluster cells were separately analyzed.
M) Heatmap comparing the average transformed expression values in regeneration exclusive clusters (F1H1+, STAT+, Cycling, ANAX2+ Migration, DAH1).
N) Heatmap comparing the average transformed expression values in FOXA3 KO, FOXA1/2 KO, FOXA1/2/3 KO mice compared to control as well as a combination of all three with endoderm and hepatoblast controls. Note that FOXA1/2/3 KO is not on the line.
O) Same as K) except including mouse KO studies.
P) Pie chart comparison for differentially expressed genes (log2fc > 1.5, padj < 0.05) between FOXA triple KO mouse liver with control mouse liver using Reactome Pathway categories to sort genes.
Q) Gene expression comparisons between FOXA1/2/3 KO and control mouse livers for select mesoderm and neuroectoderm TFs.

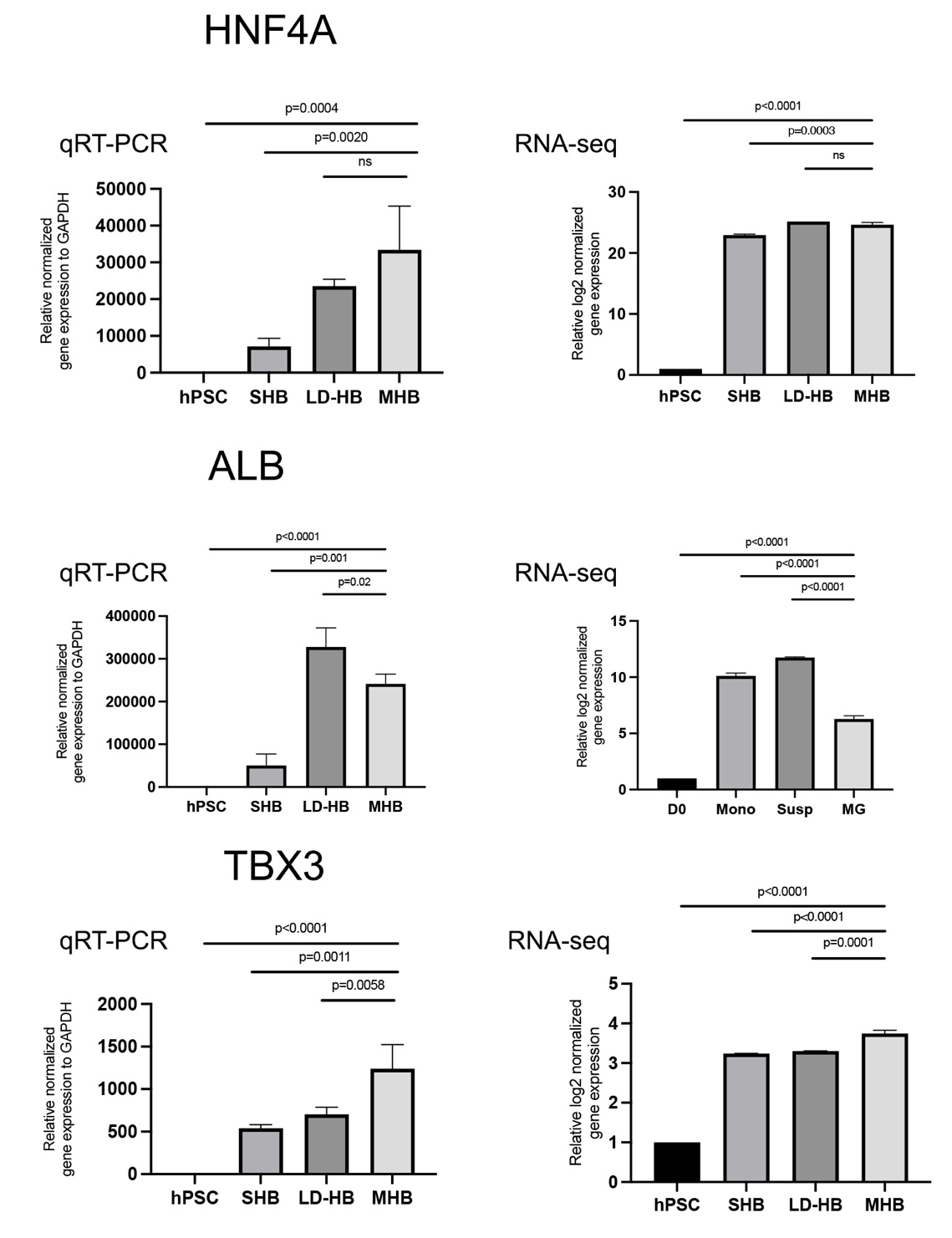

**Supplementary Figure 6:** Bar graph of gene expression kinetics (qRT-PCR and RNA-seq) for hPSC-derived hepatocyte populations (SHB (n = 3), LD-HB (n = 5), MHB (n = 3)) for HNF4A, ALB, and TBX3.

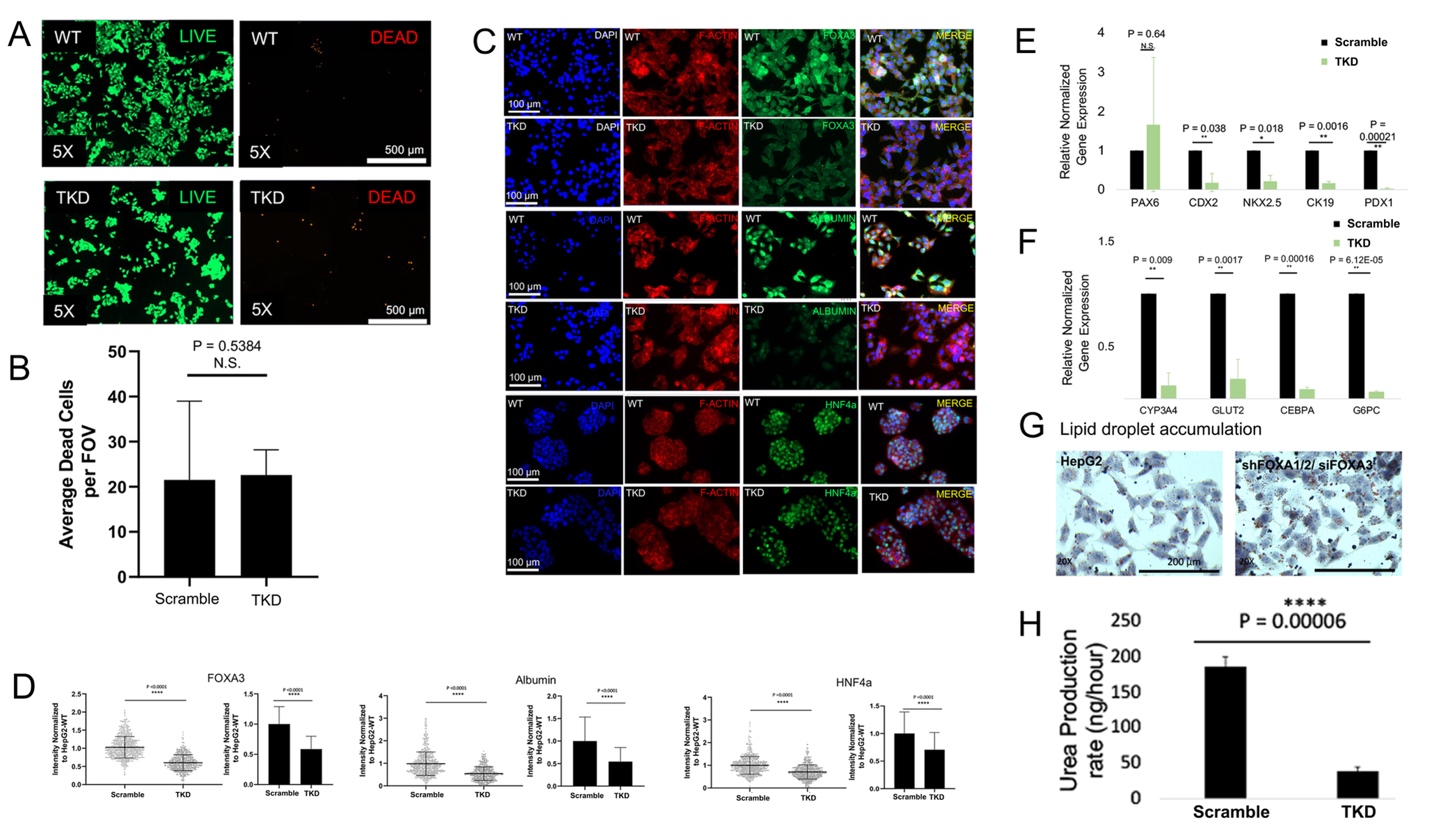

**Supplementary Figure 7**

A) Live/dead images analysis on HepG2 (WT) (top) and FOXA1/2/3 triple knockdown (TKD) cells (bottom).
B) Quantitative analysis of data in A). Mean ± SD. Significance (*) defined as P ≤ 0.05.
C) Immunocytochemistry of HepG2 (WT) (top) and FOXA1/2/3 triple knockdown (TKD) cells. DAPI (UV filter), F-ACTIN (red), FOXA3 (green), and merged (UV filter, F-ACTIN, FOXA3) are shown.
D) Quantitative analysis of data in immunocytochemistry data for FOXA3, Albumin, and HNF4A between HepG2 (WT) (top) and FOXA1/2/3 triple knockdown (TKD) cells. Mean ± SD. Significance (*) defined as P ≤ 0.05. P<0.0001 (****).
E) Bar graph of gene expression kinetics (qRT-PCR) of FOXA1/2/3 KD (TKD) HepG2 compared to shScrambled HepG2 for alternative lineage markers. N = 3 for each group compared. Plotted is mean ± SD. Significance (**) defined as P ≤ 0.05.
F) Same as E except for metabolic related genes CYP3A4, GLUT2, CEBPA, G6PC.
G) Images for HepG2 (left) and FOXA1/2/3 triple knockdown (TKD) (right) cells with staining (red) for lipid droplets.
H) Quantitative analysis of urea production between HepG2 (WT) (top) and FOXA1/2/3 triple knockdown (TKD) cells. Mean ± SD. Significance (*) defined as P ≤ 0.05. P<0.0001 (****).

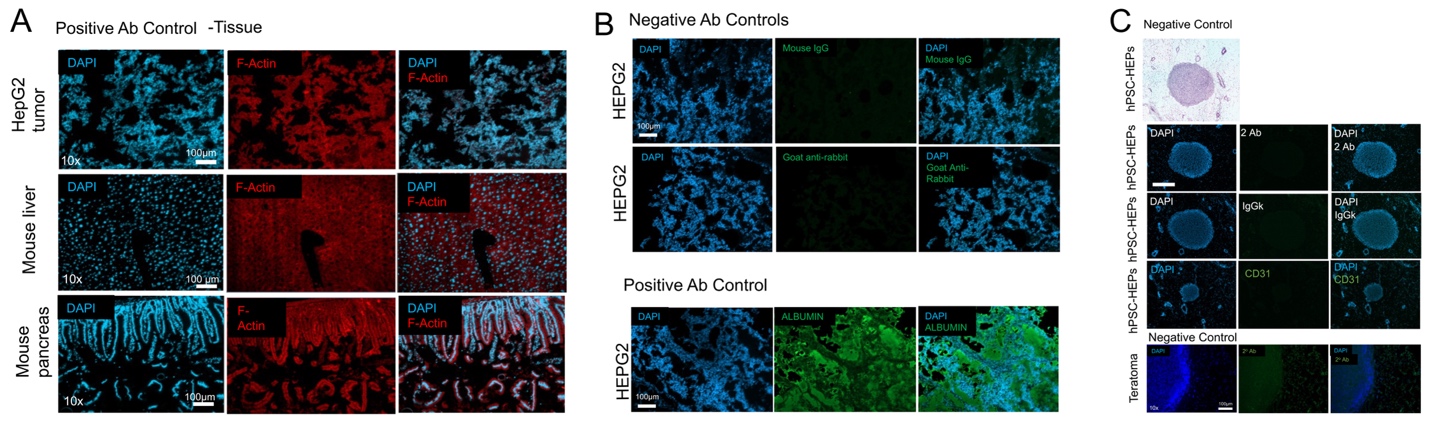

**Supplementary Figure 8**

A) Immunocytochemistry of Ab control (positive) tissues, HepG2 tumor (top), mouse liver (middle), mouse pancreas (bottom). DAPI (UV filter), F-Actin, and merged (UV and F-Actin) are shown.
B) Immunocytochemistry of Ab control (negative and positive) HepG2 tumor. DAPI (UV filter), Mouse IgG (top) or Goat anti-rabbit (bottom), and merged (UV and Mouse IgG (top), UV and Goat anti-rabbit (bottom)) are shown. Positive Ab HepG2 (bottom) stained for DAPI (UV filter), Albumin, and merged (UV and Albumin).
C) Immunocytochemistry controls for 8 week DE transplant. DAPI (UV filter), 2**°** Ab, and merged (UV and 2**°** Ab) are shown (top). DAPI (UV filter), IgGk, and merged (UV and IgGk) are shown (middle). DAPI (UV filter), CD31, and merged (UV and CD31) are shown (bottom). Immunocytochemistry of teratoma. DAPI (UV filter), 2**°** Ab, and merged (UV and 2**°** Ab) are shown.

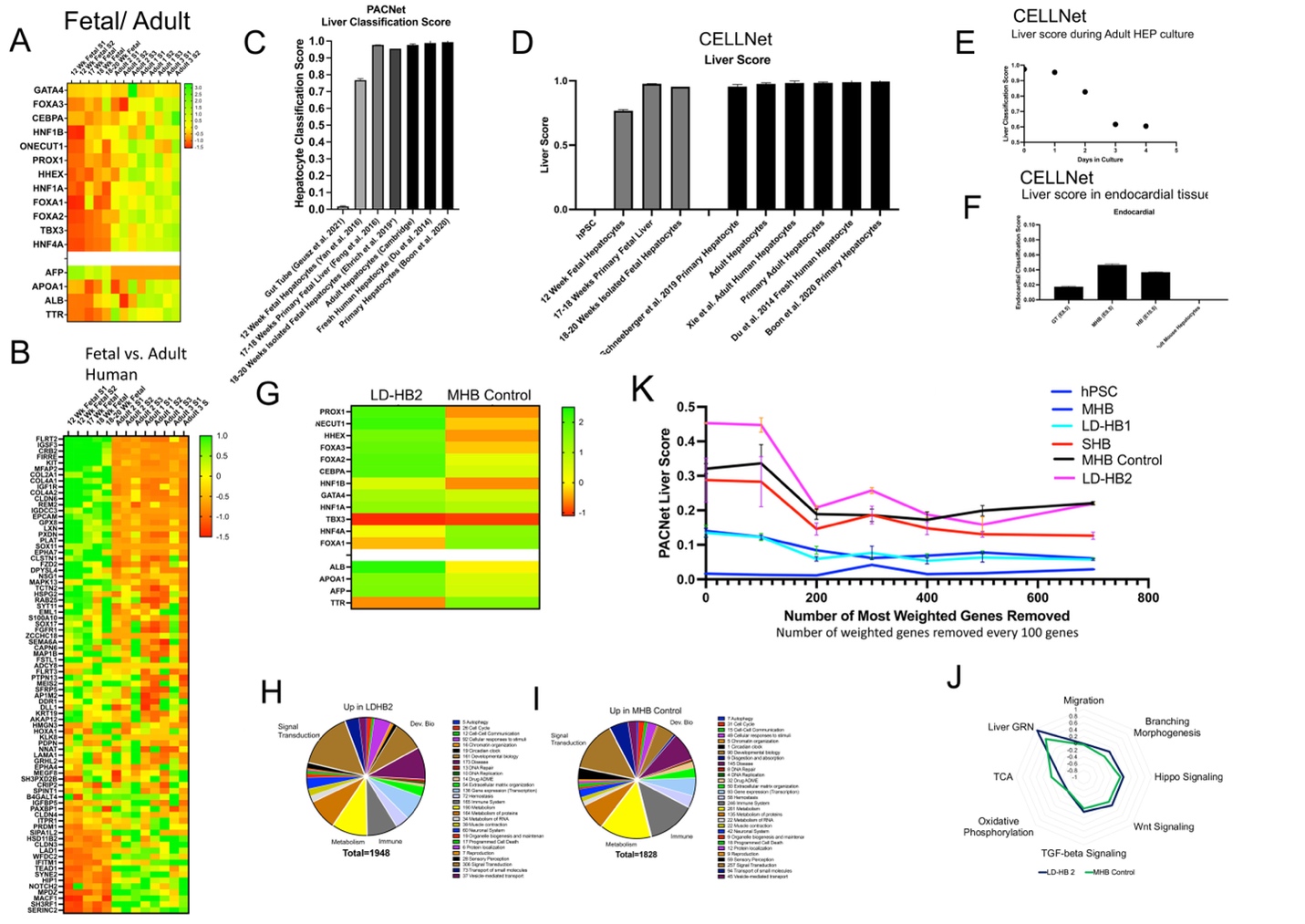

**Supplementary Figure 9**

A) Heatmap comparing the average transformed expression values for human fetal liver (Yan et al. (12 week, n = 2), Feng et al. 2016 (17-18 week, n = 2), Ehrlich et al. (18-20 week, n = 2)) and human adult hepatocytes (University of Cambridge (n = 2), Du et al. 2014 (n = 3), Boon et al. 2020 (n=2)) for select important TFs in liver organogenesis (upper) and maturation genes (lower).
B) Heatmap comparing the average transformed expression values for human fetal liver (Yan et al. (12 week, n = 2), Feng et al. 2016 (17-18 week, n = 2), Ehrlich et al. (18-20 week, n = 2)) and human adult hepatocytes (University of Cambridge (n = 2), Du et al. 2014 (n = 3), Boon et al. 2020 (n = 2)) for gut development genes sorted by difference in expression between 12 week fetal and adult.
C) Comparison of PACNet Liver Classification scores for human gut tube (Geusz et al. 2021, n = 3) , human fetal liver (Yan et al. (12 week, n = 2), Feng et al. 2016 (17-18 week, n = 2), Ehrlich et al. (18-20 week, n = 2)) and human adult hepatocytes (University of Cambridge (n = 2), Du et al. 2014 (n = 3), Boon et al. 2020 (n = 2)).
D) Same as C) except using CellNet classification scores, and additionally including adult hepatocytes from Schneeberger et al. 2019 (n = 3), and Xie et al. 2019 (n = 9)
E) CellNet classification score for adult hepatocytes at different numbers of days in culture (n =1 for each timepoint)
F) Endocardial classification score (negative control) for mouse E8.5 gut tube, E9.5 hepatoblasts, E10.5 hepatoblasts (Lotto et al.) determined by SingleCellNet and the tabulaMuris classifier. Plotted is mean ± SE.
G) Heatmap comparing the average transformed expression values for Heatmap comparing the average transformed expression values in our more mature hPSC-derived hepatocytes, LD-HB2 (n = 2) and MHB Control (n = 4) for select important TFs in liver organogenesis (upper) and maturation genes (lower).
H) Pie chart comparison for differentially expressed genes (log2fc > 0.5, padj < 0.05) up in LD-HB2 (n = 2) compared to MHB Control (n = 4) using Reactome Pathway categories to sort genes.
I) Same as H) except using genes up in MHB Control relative to LD-HB2.
J) Radar plot comparing the average transformed expression scores between MHB Control (n = 4) and LD-HB2 (n = 2) for select GO pathways based on the differential expression analysis.
K) PACNet classification scores for our hPSC-derived hepatocyte conditions (hPSC (n = 2), MHB (n =3), LD-HB1 (n = 3), SHB (n = 3), MHB Control (n = 4), LD-HB2 (n = 2) recalculated after removal of the 100 gene pairs used for the previous classification. Each iteration removes an additional 100 gene pairs from the analysis. Plotted is mean ± SD.

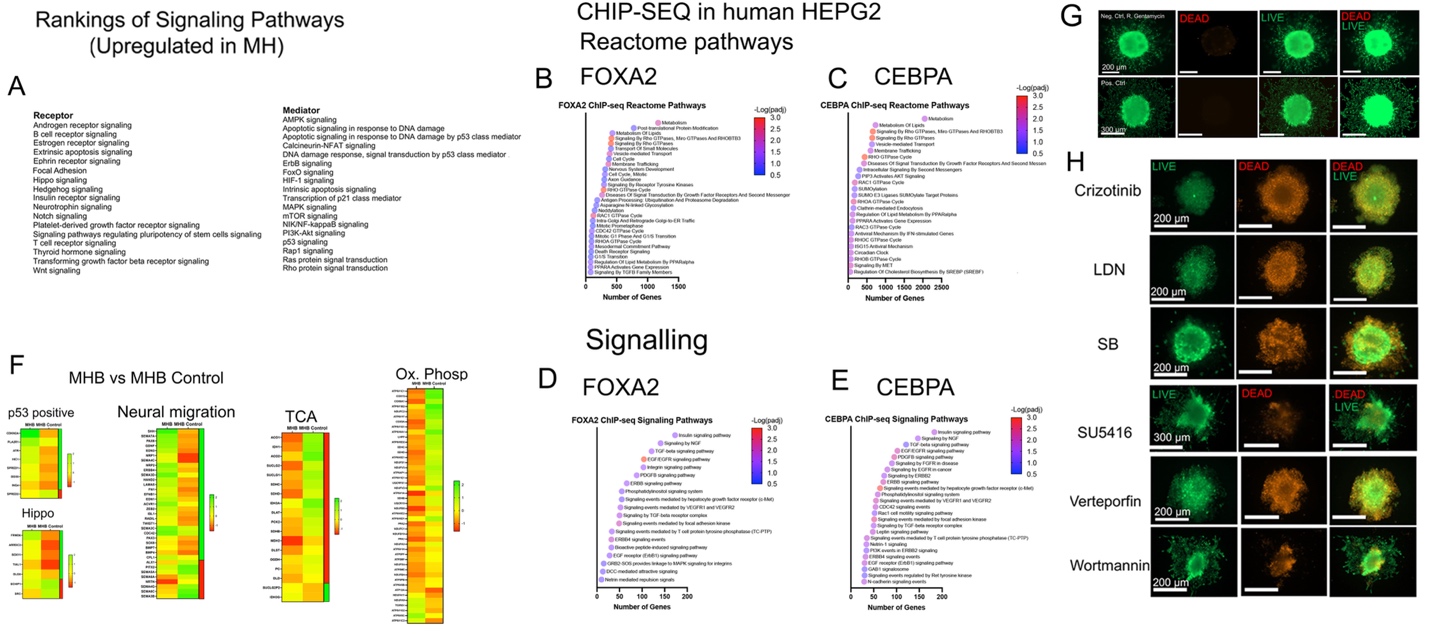

**Supplementary Figure 10**

A) List of Gene Ontology and Kegg signaling pathways found to be significantly enriched in mouse MHB (E9.5) when compared to other timepoints (E7.5, E8.5, E10.5). Lists sorted based on receptor and mediator pathways.
B) ChIP-seq gene pathway enrichment analysis for FOXA2 showing the most significantly enriched (padj < 0.05) pathways within the Reactome database sorted based on the number of genes impacted by the transcription factor.
C) Same as B) except for CEBPA
D) ChIP-seq gene pathway enrichment analysis for FOXA2 showing the most significantly enriched (padj < 0.05) signaling pathways within the Kegg database sorted based on the number of genes impacted by the transcription factor.
E) Same as D) except for CEBPA
F) Heatmap comparing the average transformed expression values between MHB (n = 3) and MHB Control (n = 4) for select gene ontology signaling pathways (positive regulating p53 signaling genes (GO:0043517), positive regulating hippo signaling genes (GO:0035332), promoting neural crest migration genes (GO:0001755), and metabolic pathways (positive regulating genes in TCA cycle (GO:0006099) and positive regulating genes in oxidative phosphorylation (GO:0006119).
G) Images of live (green) /dead (red) assay for cell viability after chemical treatment for the negative control. Enhanced images (green) shown.
H) Images of live (green) /dead (red) assay for cell viability after chemical treatment. All treatments cause cell death except SU5416.

**SUPPLEMENTAL METHODS**

**Reagents/Materials**

RPMI 1640 medium with GlutaMAX (Cat. #: 61870036), Knockout Serum Replacement (KOSR) (Cat. #: 1082810), DMEM, high glucose, GlutaMAX™ Supplement, pyruvate (Cat. #: 10569010), IMDM medium, Cat. #: 31980030, Ham’s F-12, Cat. #: 11765054, N-2 supplement, Cat. #: 17502048, L-Glutamine, Cat. #: 25030081, L-Glutamine, (Catalog #: 25030081), Fetal Bovine Serum (Cat. #: A3160701), Penicillin-Streptomycin (P/S) (10000 U/ml) (15140122), B27™ Supplement (50x), serum-free (Cat. #:17504044), 0.05% Trypsin-EDTA (Cat. #: Cat. #: V22887), Vybrant DiO Cell-Labeling Solution (Cat. #: V22886), Vybrant Dil Cell-Labeling Solution (Cat. #: V22885), L-Ascorbic acid, (Cat. #: A61-25), were purchased from Thermofisher. mTESR1 medium (Cat. #: 85850), Accutase (Cat. #: 07920), Dispase (Cat. #: 07923), Gentle dissociation reagent (Cat. #: 100-0485), Y27632 (ROCK) Inhibitor (Cat. #: 72304), Human Recombinant Activin (Cat. #: 7800.1). Dexamethasone (Cat. #: 72092), Human Recombinant Growth Factor (HGF), (Cat. #: 78019.1), Human Recombinant Oncostatin (M)), (Cat. #: 78094), KGF (Human Recombinant FGF-7), (Cat. #: 78046), bFGF (Human Recombinant), (Cat. #: 78003.1) were purchased form StemCell Technologies. N2 Supplement (100x) (GIBCO, Cat. #: 17502001), CHIR99021 (Cat. #: SML1046-5MG), EGM™- 2 Endothelial Cell Growth Medium-2 Bulletkit™ (Lonza, Catalog #: CC-3162), MG (Growth factor-free) (Cat. #: 354230), Collagen, rat tail, (Cat. #: 354236) was purchased from Corning. Aurum Total RNA Mini Kit (Cat. #: 7326820), DNase I (Cat. #: 7326828), iTaq Universal SYBR Green Supermix (Cat. #: 1725121), and iScript cDNA Synthesis Kit (Cat. #: 1708891), 96-well PCR plates (Cat. #: L223080) were purchased from Bio-Rad Vybrant DiD (red, Invitrogen, Cat. #V22887), Dil (yellow, Invitrogen, Cat. # V2885), or DiO (green, Invitrogen, Cat. # V22886), Tissue Culture Treated 24-well plate (LPS, Cat. #: 702001), 75 cm^2^ Polystyrene tissue culture-Treated Flasks (LPS, Cat. #:708003), 60 mm tissue culture treated dishes (LPS, Cat. #: 705001), 384-well round bottom, ultra-low attachment spheroid microplates (Corning, Cat. #: 3830), 96-well Cell Culture Plate (LPS, Cat. #: 701001), 6-well Cell Culture Plate (LPS, Cat. #: 703001), 96-well PCR plates (LPS, Cat. #: L223080), PCR Plate Covers (LPS, Cat. #: HOTS-100). Shifferdecker Staining Jar (Catalog No: 70314-05, Electron Microscopy Services), Slide rack (Cat. #: 70312-24, Electron Microscopy Services), Tissue Tek Base Molds for Embedding Rings (Cat. #: 62527-38,4124; Electron Microscopy Services), Millenia 2.0 Adhesion Slides, 75 x 25 mm (Cat. #: 71863-01, Electron Microscopy Services), Microprocessing/Embedding Cassettes (Cat. #: 70073-B, Electron Microscopy Services), Eosin Y (Cat. #: DcE-40, Electron Microscopy Services), Hematoxylin (Catalog No: DcH-48, Electron Microscopy Services), Permount™ Mounting Medium (Cat. #: 17986-01, Electron Microscopy Services), Cover-slips (Cat. #: 12-542-AP, Fischer Scientific). All primers for qRT-PCR were purchased from either Integrated DNA technologies (IDT), Sigma Aldrich, or Thermofisher. The following small molecule pathway inhibitors were ordered: LDN193189 Hydrochloride, Sigma, Cat.#: SML0559-5MG, SB431542, StemCell Technologies Cat. No: 72232, Y-27632 2 inhibitor (ROCK Inhibitor), MyBiosource, Cat. #: MBS577605; Crizotinib (PF-02341066), Selleck Chem Cat. #: S1068, Wortmannin, Selleck Chem, Cat. #: S2758, SU5416, Cayman Chemicals, Cat. #: 13342; A83-01, Stemgent, Cat. #:  04-0014, Verteporfin, Sigma, Cat. #: SML0534-5MG. Bovine serum albumin (BSA), Cat.# A9576-50ML, and 1-Thioglycerol, Cat#: M1753-100ML were purchased from Sigma.

**Cell lines**

iPSC (induced pluripotent stem cell) cell line: ATCC-BXS0114 Human (African American Female) Induced Pluripotent stem cells (iPSC, ACS-1028™). Embryonic stem cell line: UCSF4 (human embryonic stem cell (hESC), female, NIH Registry (0044), University of California San Francisco (UCSF). BU3-10-Cre02 Human (Caucasian Male) induced pluripotent stem cells (WiCell Cat. #: CREM003i-BU3C2) were obtained as a kind donation from Dr. Laertis Ikonomou (University at Buffalo). HepG2 liver carcinoma cells (ATCC®, Cat. #: HB-8065). Human umbilical vein endothelial cells (HUVEC) (Lonza®, Cat. #CC-2935). Human foreskin fibroblasts (HFFs) were obtained as a kind donation from Dr. Stelios Andreadis (University at Buffalo).

**Antibodies**

Mouse anti-human AFP monoclonal antibody (Cat. #: sc-130302, Santa Cruz Biotechnology). Rabbit anti-human albumin (Alb) monoclonal antibody (Cat. #: 109-4133, Rockland). Mouse anti-human CDX2 monoclonal antibody (Cat. #: sc-393572, Santa Cruz Biotechnology). Mouse anti-human SOX2 monoclonal antibody (Cat. #: sc-365823, Santa Cruz Biotechnology). Mouse anti-human CD31 (PECAM-1) monoclonal antibody (Cat. #: sc-71872, Santa Cruz Biotechnology). Mouse anti-human Foxa2 monoclonal (Cat. #: MA5-15542, Thermo Fisher). Mouse anti-human HNF4a monoclonal antibody (H-1, Cat. #: sc-374229, Santa Cruz Biotechnology). Mouse anti-human SMA monoclonal antibody (Cat. #: sc-53015, Santa Cruz Biotechnologies), Mouse anti-human TBX3 monoclonal antibody (Cat. #: sc-166623, Santa Cruz Biotechnologies). Rabbit IgG (Cat. # PI31235, Thermofisher). Mouse IgG2a kappa (Cat. # 5013049, Thermofisher). Mouse IgG2b kappa (Cat. # 5013052, Thermofisher).

**Feeder-free culture (maintenance), harvesting, and collecting of hPSC**

Methods are as described previously (Ogoke, Guiggey et al. 2021).

**Culture of cell lines (HepG2, HUVEC, HFF)**

HepG2 hepatoblast carcinoma cells (ATCC HB-8065) and human foreskin fibroblast (HFF) cells (ATCC SCRC-1041) were passaged at a 1:10 dilution on tissue culture-treated T-75 flasks in high glucose DMEM (Thermofisher) containing 10% FBS (ThermoFisher) and 1% P/S. Medium was changed every other day.

HUVEC (Lonza) were cultured in complete EGM-2 (Lonza) medium containing 1% P/S, with medium changed every other day. Modified EGM2, as described below, was used for stem cell experiments.

***In vivo* transplantation assay**

The transplant procedure was based upon prior studies (Parashurama, Nahmias et al. 2008). All animal procedures were approved by the Institutional Administrative Panel on Laboratory Animal Care at University a Buffalo, State University of New York. 1 x 10^6^ control hPSC and hPSC-derived DE combined with Human foreskin fibroblasts (HFF) (4:1 ratio of DE:HFF), and HEPG2 (human liver cancer), was harvested from *in vitro* monolayer culture using the appropriate dissociation reagent, washed in cell-appropriate medium, and collected in microcentrifuge tubes. hPSC-derived cells were transplanted in 8-10 week-old female immunodeficient NOD-SCID mice (Jackson Laboratories). Mice were anesthetized and maintained with inhaled isoflurane (1-3%). hPSC-derived cells were directly mixed on ice with 50 µL of growth factor free Matrigel (MG), or mixed with HFF and then mixed with MG. The MG/ cell mixture was then transplanted subcutaneously in the left hindlimb of NOD-SCID female mice using a 24-gauge needle. After cell transplantation, mice were kept under isoflurane for a few minutes to allow the cell/gel combination to solidify *in vivo*. Animals were recovered, and resumed spontaneous respiration on temperature-controlled heating pads. Mice were monitored carefully. After 4 weeks, mice were sacrificed for tissue assessment as described below.

**Design of culture system**

Since our unique transplantation model supported the hLD-MESC hypothesis, we wanted to develop an *in vitro* system based on this (**Fig. 1A-C**). Our focus on early LO precluded the use of traditional instructive/maturating factors like HGF, oncostatin, dexamethasone. Further, we noted that sequential instructive factors can provide mixed signals to a population of differentiating cells, for example, potentially signaling proliferation in some cells and alternate cell fates in others, and thus we chose to eliminate sequential factor design. Instead, we chose a single medium formulation, without any changes, which was employed for the entirety of differentiation under hypoxic conditions ^82,83^. In the first condition, we chose a nutrient rich medium containing, SFD previously used for maintaining hPSC-derived endoderm progenitor cells, the precursor to HBs, with minimal growth factors ^84^. We retained KGF (FGF7) in the medium, since KGF has been shown to induce hPSC-differentiation of endoderm towards GT ^80^. For the second strategy, we used data from ENRICHR pathway analysis (**Table 7, Sup. Tables 8-9)** to identify that VEGF, EGF, FGF2, and IGF-1 signaling were upregulated in the MH population. Each of these factors has been linked to liver differentiation and growth ^44, 85-89^. We found that EGM-2 medium, a commercially available medium employed VEGF, EGF, IGF-1, and FGF (**Fig. 2G**), and therefore we employed EGM-2 medium, but removed both hydrocortisone and serum (**Fig. 2G**). We also applied our knowledge of the molecular control of liver development which suggests that liver genes are primed to activate even within day 6 hPSC-derived GT endoderm ^80^. This molecular control concept supports the idea that spontaneous hepatic differentiation can occur from endoderm in the absence of formal instructive factors, which we and others have reported ^90, 91^.

**Preparation of SFD medium**

Methods are as described previously (Ogoke, Guiggey et al. 2021).

**Preparation of H + M medium**

Base (control) medium contained RPMI, 1% B27, 0.2% Knockout serum (KOSR), and 1% P/S. The modified EGM-2 medium contained EGM-2 basal medium (Lonza), 2% KOSR, 0.5% B-27, 0.05% long R3 insulin growth factor (R3-IGF), 0.05% epidermal growth factor (EGF), 0.2% fibroblast growth factor 2 (FGF2), 0.05% vascular endothelial growth factor (VEGF), 0.05% ascorbic acid, 0.05% heparin, 0.05% gentamycin, 1% P/S. The absolute concentrations of growth factors and heparin within EGM-2 are not known. Hydrocortisone and fetal bovine serum (FBS) were not added to the modified EGM-2, medium because of their complex effects on cells, and the FBS was replaced with KOSR (Stemcell technologies) in the modified EGM-2 medium, which is a defined serum. We mixed the base medium and the modified EGM-2 medium at a ratio of 1:1 and called this Hepatic and Mesenchymal medium (H + M medium).

**DE induction from human stem cells under hypoxic (5% O_2_) conditions**

Methods are as described previously (Ogoke, Guiggey et al. 2021). SFD medium contains RPMI supplemented with 75 % IMDM, 25% Ham’s F12, 0.5% N2 Supplement, 0.5% B27 supplement, 2.5 ng/ml FGF2, 1% Penicillin + Streptomycin, 0.05% Bovine Serum Albumin, 2mM Glutamine, 0.5mM Ascorbic Acid, and 0.4mM Monothioglycerol (MTG).

**HB induction using growth factor GF (+) differentiation protocol**

We adopted a hepatic differentiation protocol from (Takebe, Sekine et al. 2013) but we added an additional GT endoderm stage. Under hypoxic conditions (5% O_2_) until day 14, 2 x 10^5^ DE cells were seeded per well of a 24-well plate. Following DE induction by day 4, cells were induced to form GT endoderm using SFD media supplemented with 25 ng/mL (KGF/FGF-7). On day 6 of culture after GT induction, cells were treated in SFD medium with bone morphogenetic protein 4 (BMP4, 10 ng/ml), and fibroblast growth factor 2 (FGF2, 20 ng/ml) for specification of liver cell fate. On days 10-14, cells were treated with hepatocyte growth factor (HGF, 10 ng/ml), oncostatin (20 ng/ml), and dexamethasone (100 nM). 500 µL of culture media was used for daily medium changes.

**HB cell induction using GF (-) containing SFD medium**

Under hypoxic conditions (5% O_2_) until day 14, hPSC-derived DE progenitor cells derived on day 4 were induced to early HBs using a protocol in which no additional GF are added, to enable spontaneous hepatic differentiation. Briefly DE progenitor cells were incubated with SFD medium supplemented with (KGF/FGF7) at 25 ng/mL (in addition to FGF2 (2.5 ng/mL), from day 4-14. Differentiation throughout this protocol utilized 24-well plates and 500 µL of culture medium for daily medium changes.

**HB cell induction using hepatic and mesenchymal (H + M) medium**

hPSC-derived DE were differentiated towards HBs from day 4 to day 14 using H + M medium (defined above), under hypoxic conditions (5% O_2_). 2 x 10^5^ DE cells were seeded per well of a 24-well plate. The protocol consisted of daily medium changes (500 µL per well). 500 µL of culture medium was used for daily medium changes in each well of a 24-well plate.

**Fluorescent dye-labeling of stem cell-derived progenitors**

hPSC-derived progenitor cells at different stages of differentiation were harvested (500 µL/ well Accutase, 5 minutes, 37^o^C) and adjusted to a final concentration of 1 x 10^6^ cells/ mL appropriate medium. Subsequently, 5 µL of either Vybrant DiD (Invitrogen), Dil (Invitrogen), or DiO (Invitrogen) cell-labeling solution was added to 1 ml of cell suspension in a microcentrifuge tube, incubated for 20 minutes at 37°C, centrifuged at 1500 RPM for 5 minutes. The remaining cell pellet was then washed twice in fresh culture media before being seeded for liver organoid formation.

**Live/dead assay for thin cell sheet and liver organoids**

Organoids were assayed for cell viability using a live/dead reagent kit (Biotium, Catalog #30002-T). Briefly, kit reagents consisted of stock reagents: Calcien AM (live) and Ethidium bromide (EthD-III, dead) which were warmed to room temperature before being diluted to 2 µM and 4 µM respectively in serum-free DMEM containing 1%P/S. Organoids were rinsed once with PBS and then incubated for 3 hours in live/dead media reagents (37°C, 5% O_2_). Tissues, cultivated in incubation medium are then imaged for viability under green (live cells) and red fluorescence (dead cells) which will indicate cell viability. Images are obtained under fluorescence using Zeiss Axiovision fluorescence microscope (Observer Z.1).

**GT organoids**

***GT endoderm organoid formation***

To obtain GT organoids, hPSCs-derived DE (day 4) cells were differentiated on feeder-free, MG-coated plates (1:15 dilution in high glucose DMEM) under hypoxic conditions (5% O_2_) until day 6 in SFD medium. Next, differentiation medium was aspirated from each well and 500 µL of Accutase (Stemcell technologies) was added to each well and incubate for 5 minutes at 37^o^C, collected into a 15 mL tube, and centrifuged at 1000 RPM for 5 min. The medium containing Accutase was then aspirated, and cells were washed in fresh medium, and re-suspended in SFD medium containing ROCK inhibitor (Stemcell technologies) (1:1000) and counted. The total number of cells needed per well was 2 x 10^4^ so adjustments were done to have an appropriate number of cells for seeding. Prior to seeding cells for each organoid within each well, cells were mixed to maintain an appropriate cell distribution. GT cells were collected 50 µL and seeded into a number of wells of a sterile 384-well round bottom ultra-low attachment plate (Corning). The plate was centrifuged at 1000 RPM for 10 minutes in order to properly collect cells at the bottom center for improved compaction and organoid formation. Plates were incubated in 37°C under hypoxic conditions (5% O_2_) for 24 hours until organoids formed and imaged after each 24 hours using phase contrast and fluorescent microscopy. Organoids were further cultured and processed depending on the application.

***GT endoderm/HUVEC organoid formation***

On day 6, GT cells were harvested as described in the *GT endoderm organoid formation* section. HUVECS (previously cultured in complete EGM-2 medium) were dissociated using trypsin (0.25%) at 37^o^C for 5 minutes. Both cell types were separately collected into a 15 mL tube, centrifuged at 1000 rpm for 5 min, counted, and resuspended in SFD. The ratio of GT to HUVEC cells 1:1, therefore 1 x 10^4^ cells, of each cell type per well were used for organoid formation in a 1:1 mixture of SFD: EGM-2 culture medium. The mixture of cells was seeded as described in the *GT endoderm organoid formation* section, and organoids formed within 24 hours.

***GT organoids in EGM-2 medium***

Day 6 GT endoderm cells were obtained as described in GT organoid formation section. After Accutase digestion and harvest, the pellet was washed in fresh medium and resuspended in full EGM-2 (Lonza) medium. The collected cells were seeded (50 µL/ well) at a density of 2 x 10^4^ cells/well into a 384-well round bottom ultra-low attachment plate, and centrifuged to compact the cells into organoids, which formed in 24 hours, and further cultured in EGM-2 medium under hypoxic conditions (5% O_2_).

**Generation of microfabricated device (micropillar device) for functional analysis of mesenchymal components**

Methods are adopted from (Asmani, Velumani et al. 2018). Micropillar device was fabricated using a multi-layer microlithography technique that was previously defined. SU-8 masters were generated via spin coating, alignment, and then subsequent exposure and baking of multiple layers of SU-8 photoresists. To obtain the cross-sectional difference between the micropillar head and leg sections, a thin layer of SU-8 doped with S1813 was deposited to prevent UV light penetration to the leg section. Following this step, UV exposure was performed on an OAI mask aligner that encompassed a U-369 band pass filter. PDMS (Sylgard 184, Dow Corning) stamps were casted over the SU-8 master at 10:1 mixing ratio to demold the micropillar pattern. Additional micropillar devices were produced through replica molding from stamps in P35 petri dishes or 12 well plate. The devices were then prepared for appropriate cell seeding.

**Generation of thin organoids (microtissues) in microfabricated device for functional analysis of mesenchymal components**

Micropillar devices (Asmani, Velumani et al. 2018) were sterilized and subsequently treated with Pluronic F-127 to minimize non-specific cell adhesion to a PDMS surface. For HepG2 cells alone, a total of 1 x 10^6^ cells were used in complete DMEM (cDMEM) containing high glucose DMEM (Thermofisher) 10% FBS, 1% P/S. For HepG2: HUVEC microtissues, a 1:1 mixture of HepG2 cells and HUVEC cells was used (total cell number 1 x 10^6^), in a 1:1 mixture of cDMEM: EGM-2 medium. In the case of HUVEC cells, a total of 1 x 10^6^ cells were used in complete EGM-2 medium. For the hPSC-derived H+M cell cultured organoids were collected from 24 well plates (500 µL/well Accutase digestion, 5 minute, 37^o^C) and adjusted for 1 x 10^6^ cells per tissue device. All cell compositions were then mixed with collagen type-I at a final concentration of 3 mg/mL and introduced to the microwells via centrifugation. The collagen solution was then allowed to crosslink and maintained in 3 mL of complete H+M media in a CO_2_ incubator. The contraction force generated individual microtissues is as determined by the cantilever bending theory, $F=\delta\frac{3El}{L^{3}}$ , where El is the elastic modulus, L is the length of the cantilever, and δ is the distance the cantilever top moved. Micro-pillar deflection was measured using phase contrast microscopy by comparing the deflected position of the centroid of each pillar top to the centroid of its base. HepG2 or HepG2: HUVEC tissue devices, hPSC-derived tissues were cultivated at in 37°C, and 5% CO_2_, 5% O_2_.

**Preparation of agarose-coated microwells**

Sterile, 1 wt % agarose (1g/ 100 mL distilled H_2_0) was prepared, heated to liquid form, and pipetted (50 µL/ well) into a 96-well plate (Corning). The plate was then allowed to be cooled (25^o^C for 20 minutes) prior to cell seeding.

**Collagen type I gel formation**

Rat tail collagen type I was provided at a concentration between 3-4 mg/mL in 0.02N acetic acid (Corning; Catalog #: 354236). Current available lot of Rat tail collagen Type I (Corning; Lot# 0048003) was constituted at a concentration of 3.78 mg/mL. A working concentration of 2 mg/mL Rat tail collagen type I was utilized for downstream experiments. Adhering to manufacturer instruction, a total desired volume of 5 mL of 2mg/ml collagen gel (CG) was prepared with pre-calculated amounts of de-ionized water, 10X PBS, and 1N NaOH. For the 3.78mg/ml of stock rat tail collagen type I to polymerize, equivalent moles of NaOH are needed to neutralize the 0.02N acetic acid solvent (determined to be 60.8 µL of 1 N NaOH for 2.64 ml of 3.78 mg/mL rat tail collagen in 0.02N acetic acid needed to make 5 mL of 2 mg/mL CG). The osmolarity of the CG gel solution was adjusted with 0.5 mL of 10X PBS (pH 7.4) resulting in a final 1X concentration after gel preparation. De-ionized water was used to adjust the remaining solution to the total desired volume (calculated to be 1.80 mL for 5 mL of 2mg/mL CG). The reagents were sterilely prepared and added to a 15 mL centrifuge tube on ice in the following order: 1.80 mL de-ionized water, 0.5 mL 10X PBS, 60.8 µL of 1 N NaOH followed by 2.64 mL of 3.78mg/mL stock rat tail collagen type I in 0.02N acetic acid. The contents were briefly vortex before use in downstream applications. Prepared collagen can be stored at 4°C for 4 days.

**Organoid culture in MG droplets**

Twenty, day 15 control and H + M (twenty organoids, 300-400 µm in diameter) were collected from suspension culture and seeded onto 60 mm tissue culture treated dishes. Organoids were first collected in a 15 mL tube on ice from organoid suspension culture in well plates. The organoids were allowed to settle and the medium in the tube was aspirated (on ice). Separately, 1 mL of ice-cold diluted MG (growth factor free) mixed with RPMI control medium,1:1) was mixed with the organoids. The organoid/MG suspension was then mixed and distributed evenly inside the MG solution. To form MG droplets containing organoids, using a 200 µL pipette, a 15 µL volume of one organoid/MG solution is collected and seeded onto a 60 mm dish, for a total of 15-20 droplets. If more than organoid is seeded per droplet, it is removed and reseeded appropriately. Organoid/MG solutions are pipetted slowly onto the 60 mm dish to avoid air bubbles. Then, the droplets were incubated at 37°C, 5% O_2_ for 30 minutes to allow the MG to solidify before culture in H + M medium, of which 5 mL is slowly added to the dish and changed every 3 days. Organoids were imaged daily on the 60 mm dish using a (EVOS fluorescent, phase contrast microscope, #AMEFC4300R) at 4x, 10x, and 20x magnification.

**Organoid culture in collagen droplets**

Organoid droplet culture is the similar to the *“Organoid culture in MG droplets”* above except 1 mL of ice-cold rat tail collagen Type 1 (2 mg/ml) is employed instead of MG.

**Harvesting spheroids embedded in MG or collagen**

Spheroids embedded in gel were harvested and placed in 15 mL tubes. Accutase was then added to the tubes. The tube was placed in the cell incubator to allow the spheroids to detach from gel for 1 hour. If processing for protein or gene expression, spheroids were washed with PBS and then resuspended in 0.25% trypsin for an additional hour to break down spheroids into cells. Cells were then washed with PBS again before being processed for protein or mRNA isolation.

**Histology assay**

Organoids were fixed in 10% neutral buffered formalin for 30 minutes before being washed once in 70% ethanol and then processes for paraffin embedding, embedded in agarose (2 wt %) prior to paraffin embedding. Paraffin embedded blocks were then sectioned at 10 µm per tissue section. Antigen retrieval was completed by heating rehydrated section in 1x Tris-EDTA buffer solutions for 20 mins in a microwave. In addition, paraffin embedded, 10 µm sections, were also stained with Eosin and Hematoxylin (Electron Microscopy Services) ­­and mounted with medium before microscopy (Zeiss Axiovision fluorescence microscope (Observer Z.1)).

**Histological analysis of organoid and tissues**

Organoids were fixed in 10% neutral buffered formalin for 30 minutes before being washed once in 70% ethanol and then processes for paraffin embedding, embedded in agarose (2 wt %) prior to paraffin embedding. Paraffin embedded blocks were then sectioned at 10 µm per tissue section. Antigen retrieval was completed by heating rehydrated section in 1x Tris-EDTA buffer solutions for 20 mins in a microwave. In addition, paraffin embedded, 10 µm sections, were also stained with Eosin and Hematoxylin (Electron Microscopy Services) ­­and mounted with medium before microscopy (Zeiss Axiovision fluorescence microscope (Observer Z.1)). Harvested tissues were collected and immediately fresh frozen in dry ice. Samples were then quickly thawed before being formalin fixed (neutral buffered formalin 10% overnight). Samples were then processed for embedment in paraffin wax using standard techniques. (Histology Core, Jacobs School of Medicine and Biomedical Sciences). Tissues were then sectioned at 5 µm and stored until further processing. For hematoxylin and eosin staining, tissue sections were deparaffinized and stained using standard procedures. For organoid histological analysis, samples were fixed in 10% neutral buffered formalin for 30 minutes before being washed once in 70% ethanol and then embedded in agarose (2 wt %) prior to paraffin embedding (Histology Core, Jacobs School of Medicine and Biomedical Sciences). Paraffin embedded blocks were then sectioned at 10 µm per tissue section (Histology Core, Jacobs School of Medicine and Biomedical Sciences). In addition, paraffin embedded, 10 µm sections, were also stained with Eosin and Hematoxylin (Eosin Y Catalog Number (DcE-40), Hematoxylin Catalog Number (DcH-48)) in a similar method to whole tissues and covered with mounting medium before microscopy (Zeiss Axiovision fluorescence microscope (Observer Z.1)).

**Microscopy of organoids**

For cellular imaging, organoids placed inside 384-well plates were imaged within droplets for both phase contrast and fluorescence microscopy using a Zeiss Axiovision fluorescence microscope (Observer Z.1) using 4X, 10X objectives.

**Immunofluorescence microscopy**

For determination of intracellular localization of ALB, AFP, FOXA2, or other proteins, cells cultured on tissue culture treated dishes were washed once with PBS and then fixed with 10% neutral buffered formalin, permeabilized with 0.1% Triton X-100 in PBS, and blocked with 1% BSA in PBS for 30 minutes. Plates were then incubated with primary antibodies overnight, rinsed with PBS and incubated with secondary antibody (AlexaFlour 488, Thermofisher,1:1000 dilution) for one hour, before co-nuclear detection with DAPI (4,6-diamidino-2-phenylindole) and followed by fluorescence microscopy. Controls were non-specific antibody and secondary antibody only.

**Whole organoid fixation and immunofluorescence**

An additional staining protocol was developed for whole mount liver organoids in suspension. Briefly, liver organoids were fixed in 10% neutral buffered formalin for 1 hour and then blocked for 2 hours in 1% BSA as explained in detail (Ogoke, Yousef et al. 2021). The organoids were then incubated with primary antibody (1:100) at 4°C overnight. The following day, organoids are washed 3 times with 1% PBST (each wash 20 mins) under gentle agitation at room temperature. Secondary antibody (AlexaFlour 488, Thermofisher,1:1000 dilution) was then added for incubation at 4°C overnight and washed out as described above. DAPI incubation (10 minutes) was used to counterstain before images were obtained. Controls were secondary only and nonspecific IgG subtype with secondary antibody. Phase contrast and fluorescence microscopy were obtained using a Zeiss Axiovision fluorescence microscope (Observer Z.1).

**Urea assay**

Urea metabolite concentration analysis was done via a spectrophotometric urea assay kit (Bioassay systems, Catalog No: DIUR-100). Urea metabolite reacts directly with assay kit containing urease. Urease catalyzes the breakdown of urea into ammonia and carbon dioxide. The generate ammonia reacts turns blue in the presence of the assay reagent kit containing hypochlorite. The more urea, the bluer the sample. Per assay guidance for *in vitro* cell culture media extract, 50 µL of each cell culture supernatant was seeded into a 96 well tissue culture treated plate. For accurate analysis, a correlating blank of 50 µL of fully supplemented H+M cell culture media was used. In addition, the standard (50 µL) was also pre-diluted in H+M media for low urea concentration analysis. All analyzed samples including standard and blank were seeded in triplicate. Then 200 µL of working regent (Reagent A + Reagent B, 1:1) was introduced into each well. Subsequently, the plate was incubated at room temperature for 50 minutes before absorbance was read at 450 nm for optical density (O.D.) in a Biotek Synergy 4 Multiplate reader. The standard and the blank were used in an interpolation equation to determine urea concentrations from sample absorbance values obtained from the spectrophotometer analysis.

**ELISA assay**

To determine secreted ALB protein, sandwich ELISAs were performed using cell culture medium samples that were collected at different time points, in addition to standard samples of known concentrations. To coat the plate, captured antibody (monoclonal mouse anti-human albumin antibody – Abcam, Inc) was diluted to 2 µg/mL in 1x PBS. 100 µL of the diluted antibody was added separately to each well of a 96 well EIA/RIA high binding plate (Laboratory product sales (LPS)), and the plate was incubated overnight at 4°C with gentle shaking. The next day, the wells were washed three times with 100 µL of PBST (PBS containing 0.5% Tween-20) for 5 minutes each wash. Then 200 µL of blocking buffer (1% bovine serum albumin (BSA)) was added to each well to block residual protein-binding sites, the plates being incubated for 1 hour at 37 °C. Next, the wells were washed three times with PBST for 5 minutes each time. After that, 100 µL of the culture medium samples and controls (DMEM medium only and diluted albumin standards) were added to the appropriate wells before incubating for 1 hour at 37°C. Then the wells were washed three times with 100 µl of PBST for five minutes each time. Then, 100 µl of the detection antibody solution (dilution 2µg/ml of the biotinylated polyclonal goat anti human albumin antibody in 1X PBS, Abcam, Inc) was added to each well before incubating at 37 °C for an hour. The wells were washed three times with 100 µl of PBST for 5 mins each and 100 µl of the secondary antibody solution (dilution 2.5 µg/ml of the HRP-Streptavidin conjugated antibody in 1X PBS, Abcam, Inc.) was added to each well before incubating at 37°C for 1 hour. The wells were washed three times with PBST then 100 µl of TMB substrate was added to each well before wrapping the plate in foil and incubating it at room temperature for 30 minutes in the dark. Finally, 50 µl of stop solution was added to each well and the absorbance values were measured at 450nm (Biotek Synergy 4 Multiplate reader). ALB concentration was determined by creating a standard curve from standard samples absorbance and their respective concentrations. An equation of best fit was then created from the generated curve and used to determine sample concentrations based on respective absorbance values.

**Reverse transcriptase reaction and real-time polymerase chain reaction (RT-PCR)**

Primers are listed in table format (**Sup. Table 2**). For each experimental condition, cell lysates were collected using the Aurum Total RNA Mini Kit (Biorad). Total RNA was isolated from duplicate or triplicate samples and concentrations were measured using the NanoDrop One/One Microvolume UV-Vis Spectrophotometer, (Catalog #: ND-ONE-W). For reverse transcriptase cDNA was synthesized using the iScript cDNA synthesis kit (Biorad) and an Eppendorf 5331 MasterCycler Gradient Thermal Cycler with 5ng of RNA for each planned qRT-PCR reaction. The RT temperature protocol was 25°C for 5 min., 46°C for 20 min., 95°C for 1 min., and then either stored at 4°C or plated at 12°C prior to plating. For the PCR amplification reaction, each sample was plated in triplicate in a 10 µL per well reaction volume composed of 5µL of iTaq™ Universal SYBR® Green Supermix, and forward and reverse primers at a concentration of 300nM. The qRT-PCR reactions were run for 40 cycles (C1000 Touch Thermal Cycler, Biorad). Gene expression analysis was conducted utilizing the delta-delta-Ct method, with GAPDH used as a normalization. The PCR temperature amplification step was as follows: 98°C for 30 seconds, 98°C for an additional 15 seconds, 60°C for 30 seconds and then the process was repeated at steps 2 and 3 until 40 complete cycles had been reached. Subsequently, there was an incremental heating stage from 65°C to 95°C at an increment of 0.5°C for 5 seconds. The plate was then analyzed for relative cycle threshold (CT) value per gene of interest. All primers were purchased from either Integrated DNA technologies (IDT), Sigma Aldrich, or Thermofisher (18-22 bp in size). To quantify gene expression, CT values were calculated for the experimental and control conditions, for both the gene of interest and the housekeeping gene (GAPDH).

The 2^-ΔΔCT^ method was employed for quantification (Livak and Schmittgen 2001). For each gene, if the CT value after the PCR reaction was not detected, then 40 was chosen as the CT value reflecting the largest possible CT value using our PCR reaction. The delta-delta CT quantification method was then carried out.

**Analysis of organoid migration**

ImageJ was used to identify relative characteristics of the migrating organoids. Briefly, images were uploaded into Image J and a scale bar was set for each image uploaded before analysis. The length application in Image J was used to identify the protrusion length and thickness in the various experimental designs. The count plugin application was used to estimate the number of cords in different fields of view and over time. The trace plugin application in ImageJ identified the outline of the edge of the spheroids for overall growth kinetics. Lastly, Skeletonize3D application in image J analyzed the branching phenotype observed in migrating spheroids. With regards to Skeletonize3D plugin, images were first transformed into 8 bit grey-scale, before being thresholded to denote the peripheral edge of the organoid, and then endpoint image branching analysis performed.

**Filtering of lateral 3D migration for improved image visualization**

The raw images were loaded into the image editor GIMP 2.10.24. The brightness was increased by 20%-40% based on the brightness level of the spheroid and the contrast was reduced by 30% - 40% to decrease the visibility of the background cells and bring out the finer details of the spheroid. The Foreground Select Tool was then used to select the spheroid so the background can be completely removed.

**siRNA and shRNA knockdown of day 12 H + M treated hPSC-HEP cells**

Day 12 H + M cells were treated with shRNA FOXA1/2 and siFOXA3 using vectors, lentiviral production, transfection reagents and procedures described recently. ^39^ For siRNA transfections, separately, 6 pmol of each siRNA and 1 µL of Lipofectamine (Thermofisher) were diluted into 100 µL of serum free OptiMEM (Thermofisher) for each well in a 24-well tissue-culture-treated plate and was mixed gently. The siRNA mixtures were incubated at room temperature for 15 minutes. Subsequently, the Day 12 supernatant was aspirated, and siRNA mixture was added to the cells. Then, 600 µL of H+M medium (with no pen-strep) was added. Then, plates were placed in the incubator at 37 °C for 48 hours for RNA collection, or 72 hours for protein collection. Scramble siRNA was used as control. Medium changes post-transfection were done every 24 hours without the addition of Pen-Strep.

**Chemical screening of growth inhibition in MG culture**

Eight small molecule inhibitors were employed (**Sup. Table 1**). As mentioned above, D15 H + M (20 organoids, 300-400 µm in diameter) were collected from suspension culture and seeded onto 60 mm tissue culture treated dishes as done in the MG organoid droplet culture (one organoid per droplet, 20 droplets). After the MG gelling step (droplets were incubated at 37°C, 5% CO_2_, 5% O_2_ for 30 minutes at to allow the MG to solidify) the organoid droplets were cultured in 5 mL of H + M media supplemented with a pre-determined concentration of a small molecule inhibitor. All inhibitor/ H + M solutions were prepared the same way: the stock small molecule inhibitor was thawed from -20°C storage and diluted directly into H+M medium at room temperature. The inhibitor supplemented H + M media (5 mL) was then added directly to the 60 mm dish containing the MG/organoids and re-incubated (37°C, 5% CO_2_, 5% O_2_) for an additional 4 days. Organoids were imaged daily on the 60 mm dish using a (EVOS fluorescent, phase contrast microscope, #AMEFC4300R) at 4x, 10x, and 20x. Collective cell migration was then observed over the 4-day period. The readout of growth and migration was quantitatively assessed by light microscopy. A schematic of our screening approach to screen for growth/migration inhibitors is shown (**Fig. 5A**). Day 15 organoids in droplet culture were treated with a single small molecule inhibitor and cultured to day 18 compared to untreated controls. Phase contrast microscopy was performed on days 16-18, and growth/migration was quantified. The first criterion for determining inhibition was to demonstrate a block of growth/migration over time, by demonstrating no significant difference between day 16 to day 18. The second criterion was comparing day 18 levels of growth at a higher inhibitor concentration to a lower inhibitor concentration, to determine if there was a significant difference, indicating a block in the observed increase in growth, at the higher concentration. The third criterion was cell viability, since a loss of viability could lead to cell death, which would result in loss of growth (**Fig. 5A**), and therefore we removed conditions in which a loss of growth was due to a loss in viability. We tested A83-01 (Activin, TGF-β, Nodal, 2-200 µm), SB43152 (Activin, TGF-β, Nodal, 0.1-10 µm), Crizotinib (c-MET and ALK inhibitor, 2.5-250 nm), LDN 193189 (BMP, 50 nm-5 µm), SU5416 (VEGFR2, 0.1-10 µm), Verteporfin (VT) (Hippo (YAP inhibitor, 0.1-10 ug/mL), Wortmannin (PI3K inhibitor, 1-100 nm), and Y27632 (Rho-associated kinase (ROCK), 1-100 µm).

**Live/dead assay for biologically inhibited organoids in MG culture**

Hepatic organoids cultured in H + M medium and MG droplets until day 18 as mentioned above were assayed for cell viability using a live dead reagent kit (Biotium). Briefly, kit reagents consisted of stock reagents: Calcein AM (live) and Ethidium bromide (EthD-III, dead) which were previously warmed to room temperature before being diluted to 2 µM and 4 µM respectively in serum free culture media containing 1% pen/strep. The 60 mm dishes containing the day 18 H + M organoids are rinsed once with PBS and then incubated for 3 hours in live/dead media reagents (37°C, 5% CO_2_, 5% O_2_). Afterwards, organoid droplets kept in the incubation media are then imaged for viability under green (live cells) and red fluorescence (dead cells) which will indicate cell viability. Images are obtained under fluorescence using Zeiss Axiovision fluorescence microscope (Observer Z.1).

**Statistics**

For statistical assessment between two groups, Student’s t-test was done (type II with two tails) , and p-values equal to or less than 0.05 was considered to have statistical significance and are defined within the text and figures/figure legends. ANOVA was employed for comparison of multiple groups

**RNA-sequencing (RNA-seq)**

Total RNA integrity was determined with the Agilent Technologies Fragment Analyzer RNA assay system. The TruSeq Total RNA Stranded Library Preparation kit (Illumina) with rRNA removal was used for preparation of RNA sequencing libraries. Multiplexed libraries were individually quality controlled with the Fragment Analyzer and quantified using the Quant-iT ds DNA Assay Kit (Invitrogen). The libraries were pooled to 10 nM and diluted for qPCR using the KAPA Library Quantification Kit (Kapa Biosystems). Subsequently the pooled libraries were normalized to 4 nM based on qPCR values. All samples were sequenced on the NextSeq500 (Illumina) in midoutput mode producing 160 million paired end 75 bp reads, ~25 million per sample.

**Bulk RNA-seq Analysis**

The raw fastq files for our samples (2 hPSC, 3 SHB, 5 LD-HB, 3 MHB, and 4 MHB Control) along with WT gut tube (Geusz et al., SRR11518527, SRR11518528, SRR11518529) and fetal hepatocytes (Yan et al., SRR1663125, SRR1663126) were first aligned using the hisat2-2.1.0 GRCh38 genome and converted into sam files. The sam files were converted into bam files using samtools-1.6. FeatureCounts was used to count reads for genes from the GRCh38 genome to create a text file with the .  The text file was imported into R where it was analyzed with DESeq2. The gene expression data was normalized with Rlog normalization in DESeq2. The built in prcomp function was used on the normalized data to find the first and second principal component analysis (PCA1 and PCA2) for the Mu et al. liver and gut gene lists after inputting all the data. PCA1 and PCA2 were plotted using R plotting tools. The res function was used to obtain the adjusted p-value and log2-fold change for each gene.

**scRNA-seq Data Collection, Normalization and Filtering**

Normalized data files from the Lotto et al. (Lotto, Drissler et al. 2020) paper was downloaded from the Single Cell Portal of the Broad Institute (singlecell.broadinstitute.org/single_cell). Annotation data for the different cell types and forced-directed layout data were also downloaded. The data was previously size-factor normalized using the computeSumFactors function in R with account to library size. Feature selection, dimensionality reduction, and doublet identification were additionally performed using the scran and scater packages in R, and the data was log-transformed with the scater normalize function. Cells within the dataset were subsequently filtered based on key criteria for “unique features” (>1000 transcripts/cell) and “mitochondrial RNA content” (<20 per cell). These data were loaded into R with cell type data and further analyzed with the Seurat package. FindVariableFeature function was run with default dispersion function and mean functions was used. Further data was downloaded using a similar analysis.

We additionally downloaded scRNA-seq datasets from Mu, Xu et al. 2020 through the Gene Expression Omnibus (GEO) and further analyzed using a similar process. Processed single-nuclei RNA sequencing (snRNA-seq) data for human liver regeneration was additionally downloaded from the biorxiv preprint Matchett, Wilson-Kanamori et al. 2023, through the GEO accession GSE223561 and further analyzed using a similar process as above.

**snRNA-seq Analysis**

Data from 72,262 human nuclei was analyzed with snRNA-seq for the liver regeneration data from healthy samples (n=9), APAP-ALF samples (n=10), and NAE-ALF samples (n=12). The nuclei clusters were previously identified by Matchett et al. using the shared nearest neighbor (SNN) modularity optimization-based clustering in Seurat on the Healthy, APAP-ALF, and NAE-ALF cells variability for the purpose of constructing the SNN graph using Harmony-corrected principal components. This method resulted in nine identified clusters, which Matchett et al. named FTH1+, Central, MitoHigh, Portal1, Portal2, DAH1, STAT1+, Cycling, and ANXA2+. We further combined these clusters into two superclusters, “healthy adult human hepatocyte” and “regeneration specific clusters”. These two superclusters were based on if relative contribution of each individual cluster was <10% from the healthy hepatocytes, the cluster was considered a regeneration specific cluster. All other clusters were considered adult human hepatocyte clusters. The ANXA2+ cell cluster was identified to be a migration subset by Matchett et al. and was further analyzed.

***Cell Cluster Regrouping***

The Seurat function FindAllMarkers function was used to find globally enriched genes within each cell type. Default arguments were used except for a log2fc threshold of 0.5. Data was plotted ordered by log2fc with an adjusted-p-value cutoff of 1 x 10^-20^. Two highly significant genes (log2fc > 5, p-adj < 1E-20) were selected based on differential gene expression to represent Gut Tube (GT) (UBA52, RPL38) or the MH (DHX99, HNRNPU) cell populations. The previous cell clusters were readjusted based upon additional clustering of cell that either expressed or did not express with these genes. Cells found with expression values over a standard deviation above expected compared to the cell type for both markers (UBA52 and RPL38 for GT, DHX9 and HNRNPU for MH) were regrouped to either GT or MH respectively. For re-clustering, we used statistical analysis of 4 highly upregulated markers (two in the GT, two in the HB) to improve clustering, and performed a new heat map which had improved clustering (**Sup. Table 1)**. To determine marker expression across clusters, we employed the tools Harmony and Palantir (Lotto, Drissler et al. 2020) and re-created force-directed plots. Harmony (Korsunsky, Millard et al. 2019) groups cells by cell type, and Palantir (Setty, Kiseliovas et al. 2019) orders cells by pseudo-time and assigns probability for differentiation. The final force-directed plots demonstrate the DE to HB transition the four re-grouping markers, liver differentiation genes and major hepatic transcription factors (TFs). The hepatic TFs HEX, TBX3, and PROX1 were nearly exclusively upregulated in the MH population, as expected (Suzuki, Sekiya et al. 2008);(Sosa-Pineda, Wigle et al. 2000). EHT was visualized by comparing EPCAM (Epithelial) to DLK1 (Hepatic) expression. The MH population had high DLK1 expression and low EPCAM expression. Since the MH population was tied to growth, we analyzed cell cycle with a cell phase plot, and found that MH cells, compared to the GT and HB populations, were more actively cycling in the G2-M (mitosis) and S phases (DNA synthesis).

***T-distributed stochastic neighbor embedding (TSNE)***

Principal components were found for all cell types (DE, GT, MH, HB, HM) using the normalized log count data for all of the genes. The first 50 principal components were used to calculate the TSNE coordinates using the Seurat function, RunTSNE. The perplexity was set to 30. Data were graphed with a point size of 5 with Dimplot.

***Pathway Heatmaps***

Gene Set lists were downloaded from Mouse Genome Informatics (www.informatics.jax.org). The ScaleData function with a negative binomial model was used. The scale data was then imported into Graphpad Prism ( <https://www.graphpad.com>) where it was visualized as heatmaps using a green-yellow-red scale.

***Pathway validation***

To validate pathways in the MH cluster, we examined the expression of five liver differentiation genes (ALB, AFP, HEX, PROX1, TBX3), and these correlated in all three databases (DAVID, REACTOME, ENRICHR).

***Differential Expression Analysis-DAVID***

The FindMarkers function in Seurat was used to find differentially expressed genes between different cell types. These gene lists were able to be further filtered for genes with a log2fc > 0.5 and an adjusted-p-value less than 0.05. The Entrez gene symbols from these lists were loaded into the DAVID Bioinformatics Resources 6.8 Analysis Wizard. The Functional Annotation Tool was then used to find gene ontologies and pathways with significant enrichment. In DAVID, a Fisher’s Exact test is used to measure gene-enrichment for a specific gene set. DAVID produces a p-value from this test, and this p-value is adjusted based on the Benjamini-Hochberg method. Kegg Pathways and Gene Ontology (Biological Processes) were used, and only gene sets with an adjusted p-value < 0.3 were used in our analysis and plotted in bar graph format.

***Differential Expression-ENRICHR***

The same gene lists were used for the ENRICHR analyses as the DAVID analyses. The gene lists for the comparisons between the MH and the GT as well as the MH compared with the HB were used. Both downregulated and upregulated genes were tested separately. The Entrez gene symbols were loaded into ENRICHR. The ENRICHR gene list enrichment analysis tool was used to find significant transcription factors with the ENCODE and ChIP Enrichment Analysis (ChEA) Consensus TFs. Kegg 2021 Human, WikiPathway 2021, and GO Biological Process 2021 were the gene sets used for the pathway analysis. All data was combined into a single data table, with information about the source of the pathway and whether it was found for the upregulated or downregulated list. These data were then filtered to find gene sets with an adjusted p-value < 0.3.

***Comparison of DAVID and ENRICHR***

DAVID and ENRICHR are able to receive human and mouse genes as input. Both contain gene-set libraries from several sources (Gene Ontology, Kegg, Wiki Pathways, REACTOME, Biocarta, etc.). In addition to ontology and pathway libraries, ENRICHR additionally offers transcription, disease/drugs, cell type, and miscellaneous libraries to further analyze gene lists. Many of these pathways are exclusive to ENRICHR. For enrichment calculations, DAVID uses a modified Fisher Exact Test, called Expression Analysis Systematic Explorer (EASE), which is a more conservative test compared to the Fisher Exact Test. It calculates p-values after subtracting one gene from the List Hits (LH). These p-values were further adjusted with the linear step-up method of the Benjamini and Hochberg (DAVID: www.ncbi.nlm.nih.gov/pmc/articles/PMC2615629/). ENRICHR uses a Fisher exact test, which is corrected with a z-score permutation background correction. This process uses many random input gene lists to compute a mean rank and standard deviation from the expected rank. From this calculation, it is able to calculate a z-score, which is further combined with the p-value to score the pathways (ENRICHR Source: [www.ncbi.nlm.nih.gov/pmc/articles/PMC3637064/](http://www.ncbi.nlm.nih.gov/pmc/articles/PMC3637064/)).

***scRNA-seq Gene expression PCA plots***

Gene expression plots were plotted using the FeaturePlot function with the normalized expression data. A blue-red expression was used with blue indicating higher relative expression and red indicating lower relative expression.

***Developmental Mouse Pathway Ranking***

We performed a total of 15 comparisons involving single (e.g., MH to HB), double (e.g., MH to GT and HB), triple (e.g., MH to DE, GT, and HB) to develop a list of ranked pathways, focusing on for signaling and metabolism genes. We also performed single comparisons (e.g., MH to HB) only. For each comparison, we obtained an up-regulated and down-regulated list of pathways (KEGG and/or Biological Process). We then ranked the comparisons by the frequency in which each pathway appeared in each comparison performed, and calculated an averaged FDR value for all the comparisons.

**Average Pathway Score**

Average pathway scores were determined by running the ScaleData function with a negative binomial model to transform the expression values for all the genes within the dataset. This function adjusts the data to have a mean of 0 and a standard deviation of 1 in order to account for gene variability, so genes can be more easily compared. We then obtained curated gene lists from Kegg Pathway (https://www.genome.jp/kegg/pathway.html) and GEO (<http://geneontology.org>) biological process. The scaled expression data was then averaged for all the genes in the pathway for each individual cell or sample for scRNAseq and bulk RNAseq respectively across all the genes within each pathway. The result is a method to quickly summarize a pathway specific heatmap to quickly compare conditions and find significant differences within the data. We further created pathway heatmaps based on these scores with colors set based on a Red-Yellow-Green spectrum with a RGB color model. Listed in the scale in each figure, red indicates lower overall gene expression and green indicating higher gene expression. These scores were additionally used in radar plots as well as comparative bar graphs. These comparative bar graphs subtracted these average pathway scores between two conditions to find the change in average pathway score between the samples. In addition, the standard error was further calculated by adding the standard errors of both populations.

**Pie Chart analysis**

The FindMarkers function in Seurat was used with scRNA-seq data to find differentially expressed genes (log2fc>0.5,padj<0.01)between different cell types. For bulk RNA-seq, the results function in DESeq2 was used to find differentially expressed genes (log2fc > 1.5,padj < 0.05). These lists were found comparing MH vs. GT and HB in the developmental mouse, comparisons between several of our hPSC derived conditions, as well as human regeneration data, for both upregulated and downregulated genes. These Entrez gene lists were loaded separately into the REACTOME 3.7 Analysis Tool. From the REACTOME analysis, the number of genes can be found for each pathway group, to find the general types of genes making up the differentially expressed gene list. In order to further understand the changes in proportion of differentially expressed genes, we graphed these results using a pie chart.

**PACNet Classification**

The PACNet Classification Score was first run using the CellNet PACNET web client (http://ec2-3-89-75-200.compute-1.amazonaws.com/cl_apps/agnosticCellNet_web/), using normalized gene expression data for our samples from DESeq2. The classification score is calculated using a logistic regression model that estimates the probability of a cell belonging to a particular cell type based on the expression levels of marker genes. The classification score ranges from 0 to 1, with higher scores indicating a higher probability of a cell belonging to a particular cell type. The logistic regression model is trained on a reference cell type-specific gene expression dataset for adult tissue cells. The resulting liver classification scores for our samples were compared to the best bulk RNA-seq condition samples from 20 previously published studies for derived liver cells. We additionally wanted to look deeper at these classification scores by looking at the specific genes. From the PACNet github (https://github.com/CahanLab/PACNet) we were able to download the training data to understand the parameters that make up the score. Each gene is given a tissue specific training value (T-val). The training values are determined by an optimization procedure based on the training data. Genes that are the most highly expressed in the training data are assigned higher T-vals, whereas genes that are less informative or have no significant contribution to the cell type are assigned lower weights. Furthermore, PACNet additionally identified gene lists for positive and negative regulating genes involved in the GRN. We used these T-vals to rank each marker gene for liver and we first plotted a heatmap using only these marker genes by order of T-val as well as only the top 30 genes. We further ran the PACNet software after removing sets of 100 genes on the list to analyze the impact to the PACNet classification score.

**Pvalue Adjustment and Multiple Comparison Test**

To adjust pvalues between PACNet classification scores we calculated the pvalues for all comparisons and adjusted the pvalues utilizing the Bonferoni adjustment with an alpha equal to 0.05.

**Alternative PACNet GRN Analysis**

Since PACNet only utilizes 100 gene pairs to classify derived hepatocytes, we desired to further analyze all the genes within the PACNet GRN list. We went around accomplishing this by creating a combined dataset of samples with our conditions (n=17), gut tube (n=3), 12 week fetal liver (n=2), as well as from several different derived hepatocyte protocols including Li et al. (n=5), Velazquez et al. (n=10), and Tilson et al. (n=4). By combining all these data, we first utilized DESeq2 to take rlog normalization of all expression data, next we further transformed the expression data to fit a negative binomial model. This function adjusts the data to have a mean of 0 and a standard deviation of 1 to account for gene expression variability. We analyzed these data in a few different ways. We first found the distribution for transformed gene expression values for each condition, first creating a histogram with bin size every 0.25 from -1.25 to 2. Next in order to compare proportion of the genes relatively upregulated to genes downregulated, we used only four bins under -0.5, -0.5 to 0, 0 to 0.5, and over 0.5 and plotted these data with proportional bar graphs. Next to visualize all the genes, we first sorted the genes based on their PACNet T-vals and added the transformed expression values together one by one, which expectedly formed a line with decreasing slope. Further we found the derivative of the lines. To focus our analysis on comparing the derived hepatocytes to fetal hepatocytes instead of adult hepatocytes, we repeated this plot sorting the genes based on the average transformed expression values for fetal hepatocytes and once again found cumulative summation. In addition to find differences between fetal hepatocytes and the derived hepatocyte populations, we found the absolute cumulative expression difference between the average fetal hepatocyte and the other samples.

**Transcription Factor Correlation Analysis**

To identify important TFs correlating with maturation, we ran ordinary least squares (OLS) with the Lotto et al. murine early developmental data (E7.5-E10.5) to identify known liver TFs both positively and negatively correlating with liver maturation. We used ALB and AFP gene expression to represent liver maturation. This method identified CEBPA and FOXA3 as the most highly correlating TFs as well as TBX3 as the most negatively correlating out of the liver GRN TFs tested. We used our combined dataset of samples with our conditions (n=17), gut tube (n=3), 12 week fetal liver (n=2), Control HepG2 (n=3), FOXA1/2KD HepG2 (n=3) as well as from several different derived hepatocyte protocols including Li et al. (n=5), Velazquez et al. (n=10), and Tilson et al. (n=4). All these data were previously normalized in DESeq2 using rlog normalization. Normalized expression values were then further analyzed. We further developed a regression line by taking the ratio of TBX3 to FOXAMAX to further improve the correlation. FOXAMAX was used instead of FOXA3 due to the compensatory role between FOXA1, FOXA2, and FOXA3. All RNA-seq samples were plotted for this calculated ratio on the y axis and 1/CEBPA on the x axis. OLS was used to find the best fitting linear curve using excel. Samples for each condition were further averaged to find a single point for each condition. Further this analysis was repeated using the average ratios of cell clusters in the liver regeneration dataset (Matchett et al. 2023). An average normalized expression value was found for each cell cluster by taking the average of all the cells within the cluster. Additionally, samples in the mouse FOXA1/2/3 KO dataset (Reizel et al. 2020) using the mouse equivalents to the human genes and linear regression curves were found using the same method.

**Deep Learning Model**

For our analysis, we selected specific transcription factors (Foxa2, Foxa3, Hnf1a, Cebpa, Tbx3, Prox1, Hnf4a) as predictors, aiming to predict the RNAseq time points (mentioned earlier) variable. We partitioned our dataset, allocating 80% for training and 20% for testing using a stratified split. Prior to modeling, predictor variables were standardized to a mean of 0 and a standard deviation of 1 using the StandardScaler from sklearn. Subsequently, we constructed a feedforward neural network (also known as a multilayer perceptron) using the Keras Sequential API. This model comprised three layers: an input layer with 128 neurons (ReLU activation), a hidden layer with 64 neurons (ReLU activation), and an output layer with a single neuron employing a sigmoid activation, catering to our binary classification task. Training utilized the Adam optimizer with a binary cross-entropy loss function. The model was trained for 10 epochs, using batches of 32 samples, with performance monitored using the validation set.

**SUPPLEMENTAL TABLES**

**Sup. Table 1**: Top markers for the five major cell types (definitive endoderm, gut tube endoderm, migrating hepatoblasts, hepatoblasts, hepatomesenchyme) from E7.5 to E10.5 based on our revised clustering

| **Gene** | **Cluster** | **Average Log2FC** | **Pvalue** | **Pvalue Adj** |
| --- | --- | --- | --- | --- |
| *Spink1* | definitive endoderm | 63.36 | 1.10E-70 | 1.97E-66 |
| *Rps27* | definitive endoderm | 55.35 | 4.25E-80 | 7.62E-76 |
| *Rps29* | definitive endoderm | 50.05 | 8.13E-80 | 1.46E-75 |
| *Rplp1* | definitive endoderm | 43.11 | 1.23E-78 | 2.21E-74 |
| *Plk4* | definitive endoderm | 41.89 | 4.14E-61 | 7.43E-57 |
| *Hist1h2ap* | definitive endoderm | 40.63 | 1.17E-53 | 2.09E-49 |
| *Commd7* | definitive endoderm | 32.22 | 1.09E-25 | 1.96E-21 |
| *Tmem245* | definitive endoderm | 31.73 | 2.76E-52 | 4.94E-48 |
| *Slc2a3* | definitive endoderm | 25.38 | 2.79E-85 | 5.00E-81 |
| *Malat1* | gut tube endoderm | 195.24 | 7.74E-08 | 1.39E-03 |
| *Actb* | gut tube endoderm | 122.55 | 4.49E-65 | 8.05E-61 |
| *Rpl35* | gut tube endoderm | 102.60 | 6.36E-126 | 1.14E-121 |
| *Pyy* | gut tube endoderm | 56.89 | 1.49E-81 | 2.66E-77 |
| *Gm10076* | gut tube endoderm | 55.47 | 1.72E-117 | 3.08E-113 |
| *Tmsb10* | gut tube endoderm | 55.02 | 5.48E-120 | 9.82E-116 |
| *Comt* | gut tube endoderm | 47.79 | 8.06E-10 | 1.44E-05 |
| *Rps19* | gut tube endoderm | 42.03 | 1.79E-96 | 3.21E-92 |
| *Rpl41* | gut tube endoderm | 38.82 | 5.15E-126 | 9.22E-122 |
| *Rpl6* | gut tube endoderm | 36.09 | 1.14E-81 | 2.03E-77 |
| *H19* | migrating hepatoblasts | 130.24 | 1.77E-293 | 3.17E-289 |
| *Krt8* | migrating hepatoblasts | 83.86 | 2.44E-10 | 4.37E-06 |
| *Krt18* | migrating hepatoblasts | 78.65 | 1.33E-18 | 2.39E-14 |
| *AY036118* | migrating hepatoblasts | 59.98 | 1.49E-50 | 2.67E-46 |
| *Peg10* | migrating hepatoblasts | 54.61 | 6.50E-240 | 1.17E-235 |
| *Hist1h1e* | migrating hepatoblasts | 44.20 | 1.55E-294 | 2.79E-290 |
| *Ncl* | migrating hepatoblasts | 37.37 | 9.05E-116 | 1.62E-111 |
| *Scd2* | migrating hepatoblasts | 33.65 | 7.71E-247 | 1.38E-242 |
| *Peg3* | migrating hepatoblasts | 30.33 | 1.53E-158 | 2.74E-154 |
| *Hist1h1b* | migrating hepatoblasts | 29.60 | 1.68E-237 | 3.01E-233 |
| *Apoa1* | hepatoblasts | 208.43 | 0.00E+00 | 0.00E+00 |
| *Apoa2* | hepatoblasts | 190.83 | 0.00E+00 | 0.00E+00 |
| *Ttr* | hepatoblasts | 134.25 | 4.25E-131 | 7.61E-127 |
| *Afp* | hepatoblasts | 132.82 | 1.32E-261 | 2.36E-257 |
| *Trf* | hepatoblasts | 126.98 | 0.00E+00 | 0.00E+00 |
| *Apoe* | hepatoblasts | 125.02 | 0.00E+00 | 0.00E+00 |
| *Ambp* | hepatoblasts | 93.12 | 0.00E+00 | 0.00E+00 |
| *Serpina6* | hepatoblasts | 91.84 | 0.00E+00 | 0.00E+00 |
| *Alb* | hepatoblasts | 80.97 | 0.00E+00 | 0.00E+00 |
| *Mt1* | hepatoblasts | 80.36 | 2.51E-184 | 4.51E-180 |
| *Myl7* | hepatomesenchyme | 236.40 | 6.14E-10 | 1.10E-05 |
| *Myl4* | hepatomesenchyme | 150.45 | 5.65E-20 | 1.01E-15 |
| *Itm2a* | hepatomesenchyme | 98.68 | 1.16E-74 | 2.08E-70 |
| *Ptn* | hepatomesenchyme | 71.92 | 6.33E-112 | 1.14E-107 |
| *Serpinh1* | hepatomesenchyme | 51.34 | 1.66E-99 | 2.98E-95 |
| *Mdk* | hepatomesenchyme | 51.01 | 4.79E-59 | 8.58E-55 |
| *Tnni1* | hepatomesenchyme | 50.80 | 5.74E-20 | 1.03E-15 |
| *Tpm1* | hepatomesenchyme | 49.59 | 1.75E-11 | 3.14E-07 |
| *Lgals1* | hepatomesenchyme | 48.68 | 3.34E-79 | 5.98E-75 |
| *Col1a1* | hepatomesenchyme | 44.48 | 3.45E-107 | 6.18E-103 |

**Sup. Table 2**: Reactome pathways sorted by the difference in proportion between the upregulated and downregulated significantly expressed genes for E9.5 MHB and E10.5 HB cells

| **Migrating Hepatoblasts (MHB)** | | | | | |
| --- | --- | --- | --- | --- | --- |
| **Reactome Pathway Groups** | **MHB upregulated differentially expressed genes compared GT and HB (2102 genes)** | **Percentage of upregulated genes within each pathway group** | **MHB downregulated differentially expressed genes compared with GT and HB(1101 genes)** | **Percentage of downregulated genes within each pathway group** | **Difference upregulated genes compared to downregulated genes** |
| *Signal Transduction* | 504 | 12.72% | 216 | 7.41% | 5.31% |
| *Gene Expression (Transcription)* | 369 | 9.31% | 171 | 5.86% | 3.45% |
| *Immune System* | 360 | 9.09% | 204 | 7.00% | 2.09% |
| *Chromatin Organization* | 99 | 2.50% | 24 | 0.82% | 1.68% |
| *Cell Cycle* | 225 | 5.68% | 125 | 4.29% | 1.39% |
| *Vesicle-Mediated Transport* | 148 | 3.74% | 75 | 2.57% | 1.16% |
| *DNA Repair* | 89 | 2.25% | 39 | 1.34% | 0.91% |
| *Organelle Biogenesis and Maintenance* | 81 | 2.04% | 43 | 1.47% | 0.57% |
| *Cell-Cell Communication* | 32 | 0.81% | 7 | 0.24% | 0.57% |
| *Muscle Contraction* | 28 | 0.71% | 8 | 0.27% | 0.43% |
| *Circadian Clock* | 25 | 0.63% | 6 | 0.21% | 0.43% |
| *Reproduction* | 27 | 0.68% | 11 | 0.38% | 0.30% |
| *Digestion and Absorption* | 3 | 0.08% | 1 | 0.03% | 0.04% |
| *Neuronal System* | 38 | 0.96% | 27 | 0.93% | 0.03% |
| *Extracellular matrix organization* | 30 | 0.76% | 23 | 0.79% | -0.03% |
| *Programmed Cell Death* | 57 | 1.44% | 46 | 1.58% | -0.14% |
| *Sensory Perception* | 18 | 0.45% | 19 | 0.65% | -0.20% |
| *Hemostasis* | 90 | 2.27% | 72 | 2.47% | -0.20% |
| *Disease* | 339 | 8.56% | 259 | 8.88% | -0.33% |
| *DNA Replication* | 47 | 1.19% | 49 | 1.68% | -0.49% |
| *Autophagy* | 15 | 0.38% | 30 | 1.03% | -0.65% |
| *Developmental Biology* | 215 | 5.43% | 179 | 6.14% | -0.71% |
| *Transport of Small Molecules* | 84 | 2.12% | 86 | 2.95% | -0.83% |
| *Protein Localization* | 19 | 0.48% | 41 | 1.41% | -0.93% |
| *Metabolism of RNA* | 204 | 5.15% | 199 | 6.82% | -1.68% |
| *Metabolism of Proteins* | 366 | 9.24% | 343 | 11.76% | -2.52% |
| *Cellular Responses to Stimuli* | 189 | 4.77% | 236 | 8.09% | -3.32% |
| *Metabolism* | 261 | 6.59% | 377 | 12.93% | -6.34% |
| **Hepatoblast (HB)** | | | | | |
| **Reactome Pathway Groups** | **HB upregulated differentially expressed genes compared with GT and MH (602 genes)** | **Percentage of upregulated genes within each pathway group** | **HB downregulated differentially expressed genes compared with gut tube and migrating hepatoblast**  **(2550 genes)** | **Percentage of downregulated genes within each pathway group** | **Difference upregulated genes compared to downregulated genes** |
| *Metabolism* | 243 | 16.51% | 354 | 7.09% | 9.42% |
| *Metabolism of Proteins* | 187 | 12.70% | 467 | 9.35% | 3.36% |
| *Transport of Small Molecules* | 57 | 3.87% | 106 | 2.12% | 1.75% |
| *Hemostasis* | 54 | 3.67% | 104 | 2.08% | 1.59% |
| *Protein Localization* | 27 | 1.83% | 28 | 0.56% | 1.27% |
| *Cellular Responses to Stimuli* | 106 | 7.20% | 297 | 5.94% | 1.26% |
| *Autophagy* | 18 | 1.22% | 30 | 0.60% | 0.62% |
| *Extracellular matrix organization* | 15 | 1.02% | 37 | 0.74% | 0.28% |
| *DNA Replication* | 23 | 1.56% | 65 | 1.30% | 0.26% |
| *Programmed Cell Death* | 23 | 1.56% | 70 | 1.40% | 0.16% |
| *Sensory Perception* | 9 | 0.61% | 24 | 0.48% | 0.13% |
| *Neuronal System* | 17 | 1.15% | 53 | 1.06% | 0.09% |
| *Digestion and Absorption* | 1 | 0.07% | 2 | 0.04% | 0.03% |
| *Immune System* | 119 | 8.08% | 409 | 8.19% | -0.10% |
| *Organelle Biogenesis and Maintenance* | 26 | 1.77% | 95 | 1.90% | -0.14% |
| *Circadian Clock* | 5 | 0.34% | 25 | 0.50% | -0.16% |
| *Reproduction* | 6 | 0.41% | 31 | 0.62% | -0.21% |
| *Vesicle-Mediated Transport* | 46 | 3.13% | 167 | 3.34% | -0.22% |
| *Muscle Contraction* | 3 | 0.20% | 26 | 0.52% | -0.32% |
| *Cell-Cell Communication* | 3 | 0.20% | 36 | 0.72% | -0.52% |
| *DNA Repair* | 18 | 1.22% | 99 | 1.98% | -0.76% |
| *Cell Cycle* | 56 | 3.80% | 263 | 5.26% | -1.46% |
| *Developmental Biology* | 67 | 4.55% | 307 | 6.14% | -1.59% |
| *Disease* | 108 | 7.34% | 452 | 9.05% | -1.71% |
| *Chromatin Organization* | 4 | 0.27% | 116 | 2.32% | -2.05% |
| *Metabolism of RNA* | 57 | 3.87% | 306 | 6.12% | -2.25% |
| *Gene Expression (Transcription)* | 71 | 4.82% | 432 | 8.65% | -3.82% |
| *Signal Transduction* | 103 | 7.00% | 595 | 11.91% | -4.91% |

**Sup. Table 3**: DAVID results for Kegg Pathways using the upregulated and downregulated differentially expressed genes from E9.5 MHB compared to E10.5 HB and E8.5 GT comparison

| **Upregulated in MHB** | | | |
| --- | --- | --- | --- |
| **Term** | **Count** | **P-Value** | **Benjamini** |
| *RNA transport* | 48 | 2.8E-11 | 7.2E-09 |
| *Cell cycle* | 39 | 8.6E-11 | 0.000000011 |
| *Protein processing in endoplasmic reticulum* | 43 | 9.1E-09 | 0.00000078 |
| *Ubiquitin mediated proteolysis* | 38 | 0.000000022 | 0.0000014 |
| *Ribosome biogenesis in eukaryotes* | 27 | 0.000000051 | 0.0000026 |
| *mRNA surveillance pathway* | 29 | 0.000000088 | 0.0000037 |
| *Focal adhesion* | 46 | 0.00000024 | 0.0000089 |
| *Spliceosome* | 34 | 0.00000044 | 0.000014 |
| *Proteoglycans in cancer* | 44 | 0.00000096 | 0.000027 |
| *RNA degradation* | 23 | 0.0000096 | 0.00024 |
| *Pathways in cancer* | 67 | 0.000011 | 0.00026 |
| *Regulation of actin cytoskeleton* | 41 | 0.000047 | 0.001 |
| *Bacterial invasion of epithelial cells* | 20 | 0.00016 | 0.0031 |
| *Renal cell carcinoma* | 18 | 0.00025 | 0.0046 |
| *Transcriptional misregulation in cancer* | 32 | 0.00033 | 0.0055 |
| *Hippo signaling pathway* | 30 | 0.00034 | 0.0055 |
| *Signaling pathways regulating pluripotency of stem cells* | 28 | 0.0004 | 0.006 |
| *Small cell lung cancer* | 20 | 0.00044 | 0.0061 |
| *Pancreatic cancer* | 17 | 0.00046 | 0.0061 |
| *Adherens junction* | 18 | 0.00052 | 0.0063 |
| *Chronic myeloid leukemia* | 18 | 0.00052 | 0.0063 |
| *FoxO signaling pathway* | 27 | 0.00058 | 0.0067 |
| *ErbB signaling pathway* | 20 | 0.00071 | 0.0078 |
| *Prostate cancer* | 20 | 0.00082 | 0.0087 |
| *Endometrial cancer* | 14 | 0.0013 | 0.013 |
| *Oocyte meiosis* | 22 | 0.0029 | 0.029 |
| *Endocytosis* | 41 | 0.0033 | 0.03 |
| *Colorectal cancer* | 15 | 0.0034 | 0.03 |
| *Fanconi anemia pathway* | 13 | 0.0034 | 0.03 |
| Lysine degradation | 13 | 0.0041 | 0.035 |
| Axon guidance | 23 | 0.0077 | 0.061 |
| Non-small cell lung cancer | 13 | 0.0076 | 0.061 |
| HTLV-I infection | 41 | 0.0086 | 0.064 |
| TGF-beta signaling pathway | 17 | 0.0085 | 0.064 |
| Wnt signaling pathway | 24 | 0.011 | 0.081 |
| Long-term potentiation | 14 | 0.012 | 0.082 |
| PI3K-Akt signaling pathway | 49 | 0.012 | 0.086 |
| **Downregulated in MHB** | | | |
| **Term** | **Count** | **P-Value** | **Benjamini** |
| *Ribosome* | 94 | 1.4E-64 | 3.8E-62 |
| *Oxidative phosphorylation* | 69 | 1.3E-36 | 1.7E-34 |
| *Parkinson's disease* | 71 | 3.3E-36 | 2.9E-34 |
| *Huntington's disease* | 80 | 1.5E-34 | 9.8E-33 |
| *Alzheimer's disease* | 70 | 1.7E-29 | 9.1E-28 |
| *Non-alcoholic fatty liver disease (NAFLD)* | 59 | 1.7E-23 | 7.3E-22 |
| *Metabolic pathways* | 182 | 5.7E-14 | 2.1E-12 |
| *Spliceosome* | 39 | 1.1E-11 | 3.5E-10 |
| *Biosynthesis of antibiotics* | 48 | 8.6E-10 | 2.5E-08 |
| *Carbon metabolism* | 31 | 2.2E-08 | 5.6E-07 |
| *Proteasome* | 18 | 6.5E-08 | 1.5E-06 |
| *Citrate cycle (TCA cycle)* | 13 | 6.3E-06 | 0.00014 |
| *Cardiac muscle contraction* | 20 | 0.000018 | 0.00035 |
| *Protein processing in endoplasmic reticulum* | 32 | 0.00003 | 0.00056 |
| *Biosynthesis of amino acids* | 19 | 0.000053 | 0.00091 |
| *Glycine, serine and threonine metabolism* | 12 | 0.00038 | 0.0062 |
| *Glyoxylate and dicarboxylate metabolism* | 10 | 0.00049 | 0.0074 |
| *Complement and coagulation cascades* | 17 | 0.00057 | 0.0079 |
| *Glutathione metabolism* | 14 | 0.00058 | 0.0079 |

**Sup. Table 4**: Top differentially expressed genes (Padj<0.01, Top 25 by positive and negative log2fc) between normal hepatocyte and regenerating hepatocyte cell populations

| Regenerating Hepatocytes vs. Normal Hepatocytes | | |
| --- | --- | --- |
| Gene | Log2FoldChange  (Positive is up in normal hepatocytes compared to regenerating | Padj |
| AKAP12 | 1.70721267 | 6.792E-107 |
| BICC1 | 1.65452698 | 1.522E-149 |
| FP671120.1 | 1.58582402 | 1.028E-274 |
| ASPM | 1.39965022 | 8.468E-142 |
| FP236383.1 | 1.18404747 | 8.585E-162 |
| TNFAIP8 | 1.17216402 | 4.848E-230 |
| RELN | 1.1071955 | 2.9941E-98 |
| GMDS | 1.05578074 | 2.042E-235 |
| FMNL2 | 1.05473671 | 1.539E-174 |
| ITGAV | 1.05398081 | 5.775E-203 |
| TPM1 | 0.99810628 | 1.067E-138 |
| DTNA | 0.97749722 | 4.844E-257 |
| ASAP1 | 0.96841316 | 2.012E-252 |
| SMC4 | 0.95375761 | 2.706E-107 |
| HKDC1 | 0.9442079 | 3.142E-195 |
| CENPP | 0.92285237 | 3.8979E-78 |
| LINC00854 | 0.92149184 | 1.8007E-36 |
| CDK5RAP2 | 0.92008662 | 8.429E-161 |
| PTPRM | 0.91526182 | 6.607E-236 |
| SLC38A1 | 0.91268458 | 2.189E-196 |
| KLHL29 | 0.90759443 | 3.901E-173 |
| NDRG1 | 0.90283785 | 1.3926E-21 |
| PVT1 | 0.90177727 | 3.35E-119 |
| DIAPH3 | 0.89658977 | 3.522E-110 |
| ACSL4 | 0.89571929 | 1.5529E-84 |
| ADRA1A | -1.2084648 | 0 |
| FMO5 | -1.216798 | 2E-266 |
| INSIG1 | -1.222156 | 2.807E-296 |
| SERPINA1 | -1.2429267 | 2.3394E-34 |
| AASS | -1.2563292 | 2.67E-282 |
| TTR | -1.2684031 | 2.121E-206 |
| ADH4 | -1.273336 | 5.84E-179 |
| CYP2A7 | -1.2984887 | 1.01E-271 |
| FGA | -1.3176219 | 3.011E-209 |
| LINC00598 | -1.335324 | 1.378E-190 |
| KCNN2 | -1.3467242 | 7.721E-280 |
| MAT1A | -1.3684749 | 0 |
| APOA1 | -1.384985 | 1.3443E-37 |
| TAT | -1.4438627 | 0 |
| CTH | -1.502157 | 2.055E-273 |
| HAL | -1.5073445 | 4.563E-242 |
| CYP3A43 | -1.5943398 | 6.726E-245 |
| SAA1 | -1.6185724 | 1.05E-19 |
| APOC3 | -1.6424049 | 1.9552E-33 |
| APOE | -1.6482872 | 2.6663E-06 |
| CYP2A6 | -1.7210513 | 0 |
| HP | -1.7465084 | 2.021E-159 |
| CYP2B6 | -2.1559739 | 0 |
| CTNNA3 | -2.4560013 | 0 |
| CYP3A4 | -2.7856881 | 0 |

**Sup. Table 5**: Top differentially expressed genes (Padj<0.01, Top 25 by positive and negative log2fc) between our hPSC derived MHB population and MHB control cell populations

| MHB vs. MHB Control | | |
| --- | --- | --- |
| Gene | Log2FoldChange  (Positive is up in MHB Control compared to MHB) | Padj |
| RPL10P6 | 9.14035806 | 2.284E-26 |
| RPS3AP21 | 6.8107608 | 4.7861E-14 |
| GC | 6.71194789 | 4.8009E-44 |
| ORM1 | 6.53803685 | 4.13E-62 |
| LINC01713 | 6.39206448 | 7.2598E-13 |
| HEPACAM2 | 6.38529204 | 2.8253E-06 |
| PTPRC | 6.17862473 | 0.00048171 |
| MT1H | 6.12616057 | 9.0112E-21 |
| AKR1C1 | 6.09738727 | 0 |
| RBP3 | 6.06575029 | 2.925E-149 |
| ORM2 | 5.95448107 | 8.502E-08 |
| VNN3P | 5.93199065 | 2.9315E-05 |
| ALOXE3P1 | 5.90108511 | 5.561E-24 |
| VAV1 | 5.82762323 | 1.224E-10 |
| CCL20 | 5.81243066 | 3.9744E-16 |
| GLYAT | 5.78321916 | 6.2078E-76 |
| C3 | 5.76123196 | 0 |
| LINC01147 | 5.7313921 | 4.7292E-05 |
| FETUB | 5.6896291 | 7.403E-124 |
| AKR1B10 | 5.65455213 | 3.7303E-68 |
| TBC1D3F | 5.64913584 | 0.00093628 |
| MUCL3 | 5.61829936 | 0.000113 |
| F11 | 5.61448493 | 5.9853E-07 |
| TINAG | 5.50764013 | 8.3577E-12 |
| SERPINA6 | 5.48631886 | 0 |
| ALB | 5.48341328 | 0 |
| CER1 | -7.8549419 | 4.825E-161 |
| CD177 | -7.8871319 | 2.2295E-12 |
| CYP26A1 | -7.8914062 | 3.8033E-12 |
| RPL21P42 | -7.8960592 | 2.2852E-12 |
| PLPPR5 | -7.9643879 | 1.0878E-13 |
| STIP1P3 | -8.1012015 | 3.3669E-13 |
| EIF4A1P4 | -8.1116548 | 3.9241E-13 |
| SEZ6L | -8.208345 | 4.6065E-39 |
| EEF1A1P1 | -8.2092693 | 4.9258E-13 |
| RN7SKP9 | -8.2986936 | 1.692E-224 |
| GAPDHP63 | -8.3320041 | 8.8306E-14 |
| KRT18P68 | -8.4060481 | 4.8977E-14 |
| APLNR | -8.4593674 | 3.1203E-65 |
| GAPDHP71 | -8.6053142 | 6.832E-15 |
| CMKLR1 | -8.6141569 | 6.1123E-16 |
| EYA2 | -8.6259093 | 3.7853E-15 |
| FTH1P13 | -8.6456773 | 6.7449E-15 |
| RN7SL587P | -8.726555 | 1.0556E-80 |
| LEFTY2 | -8.8909002 | 9.4059E-25 |
| GAPDHP35 | -8.9053127 | 5.4182E-16 |
| RPS3P6 | -8.9332363 | 2.0415E-16 |
| NTRK3 | -8.9775358 | 4.2321E-25 |
| LDHAP5 | -9.0707059 | 9.0225E-17 |
| DKK2 | -9.1912717 | 6.0352E-26 |
| RN7SL665P | -9.2266917 | 1.9182E-17 |

**Sup. Table 6**: Top differentially expressed genes (Padj<0.01, Top 25 by positive and negative log2fc) between our hPSC derived LD-HB1 population and MHB cell populations

| LD-HB1 vs. MHB | | |
| --- | --- | --- |
| Gene | Log2FoldChange  (Positive is up in MHB compared to LD-HB1) | Padj |
| PRR9 | 6.820584 | 7.3622E-07 |
| TMEM207 | 6.64364221 | 3.8098E-09 |
| GPX5 | 6.5160227 | 3.7207E-06 |
| SERPINA7 | 6.18272826 | 5.5585E-41 |
| NFIA-AS2 | 6.02565651 | 1.5382E-05 |
| TAGLN3 | 5.96176673 | 6.4727E-05 |
| BHLHE22 | 5.80745222 | 0.00012441 |
| LRRIQ1 | 5.55828724 | 9.943E-05 |
| OR5BA1P | 5.39603888 | 0.00020114 |
| CHRNA1 | 5.38467032 | 0.00080755 |
| MARCHF4 | 5.22185902 | 3.885E-27 |
| MIR3689F | 5.21876397 | 0.00145452 |
| HS3ST2 | 5.10911791 | 1.9514E-07 |
| MANCR | 5.07732818 | 2.2432E-09 |
| ACTA1 | 5.00094571 | 1.8494E-58 |
| ACTG2 | 4.95535957 | 6.1791E-25 |
| FAP | 4.91263647 | 3.5739E-12 |
| LINC02488 | 4.79204024 | 0.00248519 |
| WNT10A | 4.76357767 | 6.7752E-17 |
| SI | 4.75710735 | 4.5014E-18 |
| PRODH2 | 4.71332378 | 1.0187E-31 |
| OR14J1 | 4.70194666 | 0.01037492 |
| LINC01629 | 4.70192864 | 2.1178E-09 |
| CXCL6 | 4.6913048 | 4.2758E-06 |
| SLC22A6 | 4.68487263 | 1.608E-12 |
| LINC01639 | -4.9487058 | 0.00134919 |
| NFILZ | -5.004619 | 0.00112824 |
| HSPA8P14 | -5.0330937 | 0.00391416 |
| UGT1A3 | -5.044241 | 0.00414289 |
| PSG9 | -5.0470636 | 3.3871E-13 |
| C3P1 | -5.0555491 | 0.00090355 |
| DIPK1C | -5.100792 | 0.00314401 |
| LYPD5 | -5.1025935 | 0.00327917 |
| BEND4 | -5.1124298 | 5.0214E-07 |
| PSG3 | -5.1508051 | 3.1652E-05 |
| XAGE2 | -5.1674923 | 0.00385944 |
| SPTA1 | -5.1929218 | 0.00057417 |
| FGFBP1 | -5.2648315 | 0.00160102 |
| INSYN1-AS1 | -5.360191 | 0.00119632 |
| RNU6-616P | -5.3857513 | 9.5628E-06 |
| HCK | -5.503984 | 0.00098709 |
| MRPL23-AS1 | -5.541832 | 0.00052826 |
| LINC02975 | -5.6426973 | 8.6902E-05 |
| PLAAT4 | -5.6629879 | 7.5818E-05 |
| PSG2 | -5.7095317 | 0.000283 |
| LINC00520 | -5.7874967 | 0.00025802 |
| LINC02154 | -5.8137115 | 1.6391E-18 |
| ECRG4 | -5.8580724 | 0.00012799 |
| PYHIN1 | -5.9530946 | 8.3609E-05 |
| LINC01300 | -7.3996192 | 3.5822E-08 |

**Sup. Table 7**: Top differentially expressed genes (Padj<0.01, Top 25 by positive and negative log2fc) between our hPSC derived LD-HB2 population and MHB Control cell populations

| LD-HB2 vs. MHB Control | | |
| --- | --- | --- |
| Gene | Log2FoldChange  (Positive is up in MHB Control compared to LD-HB2) | Padj |
| XIST | 11.8907675 | 6.7817E-16 |
| RPL10P6 | 9.53581194 | 1.6712E-10 |
| PTPRC | 9.24435887 | 0.0027717 |
| MT1H | 8.94948356 | 5.794E-09 |
| CXCL5 | 8.19747626 | 1.1597E-07 |
| HLA-DPA1 | 8.06908671 | 0.00115764 |
| CYBB | 7.98557033 | 0.00400009 |
| RNA5-8SN2 | 7.93806329 | 0.00078905 |
| LINC01335 | 7.70989621 | 8.5513E-07 |
| HMGB1P11 | 7.4971248 | 2.0427E-06 |
| MC5R | 7.42293705 | 2.56E-06 |
| DIO2 | 7.38230707 | 3.169E-06 |
| TUBBP5 | 7.3647785 | 3.2307E-06 |
| HS6ST3 | 7.28468685 | 2.0057E-06 |
| CFL1P3 | 7.26480152 | 2.1142E-06 |
| MUC19 | 7.16119058 | 4.9946E-42 |
| SERPINA7 | 7.1427929 | 7.369E-283 |
| LINC01438 | 7.08126207 | 1.0141E-05 |
| SPRR1B | 7.04563798 | 1.4191E-05 |
| MIR3648-1 | 7.04431101 | 6.5887E-06 |
| LINC02906 | 7.012937 | 1.2244E-05 |
| TMEM207 | 6.95489237 | 1.1272E-29 |
| BSND | 6.91824298 | 1.7084E-05 |
| DOCK10 | 6.88190971 | 0.02324176 |
| PNLIPRP2 | 6.85661109 | 9.1492E-06 |
| RPS24P7 | -8.1131609 | 3.0247E-11 |
| RPL7AP36 | -8.191345 | 1.2612E-11 |
| UBA52P5 | -8.1941502 | 1.1021E-13 |
| RPL23AP69 | -8.1950634 | 1.2521E-11 |
| LDHAP1 | -8.2036011 | 1.2728E-11 |
| BTF3P7 | -8.2375982 | 8.0134E-12 |
| RPL37P6 | -8.2859231 | 5.3744E-12 |
| RPSAP29 | -8.352996 | 1.6853E-20 |
| STIP1P3 | -8.4962316 | 5.7348E-13 |
| ASS1P14 | -8.7503265 | 4.2857E-14 |
| EEF1A1P29 | -8.817435 | 2.9967E-16 |
| RPS5P8 | -8.8538845 | 1.6148E-14 |
| RPL23AP31 | -9.0135403 | 7.0686E-15 |
| RPL15P20 | -9.0240068 | 2.2464E-15 |
| GAPDHP63 | -9.0589901 | 1.5565E-15 |
| GAPDHP71 | -9.2257589 | 3.7886E-16 |
| RPS3AP36 | -9.3643214 | 5.0914E-44 |
| RPL34P27 | -9.611835 | 4.0811E-18 |
| RPL21P42 | -9.7741431 | 7.9613E-19 |
| GAPDHP35 | -9.821962 | 3.8294E-19 |
| EEF1A1P1 | -9.95119 | 9.0331E-20 |
| EIF4A2P4 | -9.968701 | 1.6421E-50 |
| LDHAP5 | -10.174361 | 7.9984E-21 |
| RPS3P6 | -10.539777 | 3.9784E-23 |
| FTH1P13 | -10.883528 | 2.3461E-24 |

**Sup. Table 8**: Top differentially expressed genes (Padj<0.01, Top 25 by positive and negative log2fc) between our hPSC derived LD-HB1 population and LD-HB2 cell populations

| LD-HB1 vs. LD-HB2 | | |
| --- | --- | --- |
| Gene | Log2FoldChange  (Positive is up in LD-HB2 compared to LD-HB1) | Padj |
| IGFBP1 | 10.0255365 | 5.103E-122 |
| F11 | 9.06752428 | 1.6459E-12 |
| ALB | 9.01313325 | 0 |
| ITIH2 | 9.00142627 | 0 |
| SLC22A9 | 8.88729006 | 3.412E-32 |
| LINC01146 | 8.54565313 | 1.6229E-29 |
| LBP | 8.38531211 | 7.0335E-42 |
| ESM1 | 8.33217348 | 2.863E-14 |
| ONECUT1 | 8.25942545 | 2.644E-201 |
| AGXT | 7.93907549 | 8.7139E-37 |
| TM4SF4 | 7.93517111 | 1.427E-292 |
| F9 | 7.91775714 | 6.3137E-09 |
| SERPINA11 | 7.89776503 | 4.434E-113 |
| AZGP1 | 7.87511842 | 2.0237E-09 |
| ALOXE3P1 | 7.81121082 | 1.389E-08 |
| LINC01717 | 7.67306908 | 3.2379E-08 |
| AMBP | 7.55673806 | 0 |
| SLC38A3 | 7.54286005 | 3.746E-186 |
| ACSM2B | 7.49771164 | 2.1289E-11 |
| ST8SIA3 | 7.46772916 | 2.8435E-11 |
| HAMP | 7.3949028 | 5.6101E-16 |
| GC | 7.37126548 | 3.4496E-31 |
| NR1H4 | 7.36862546 | 4.7379E-61 |
| F13B | 7.33698465 | 4.2729E-08 |
| AGTR1 | 7.1787714 | 0 |
| LIPN | -6.7834621 | 5.18E-05 |
| TECRL | -6.858848 | 1.3471E-05 |
| ANKRD30B | -6.8703096 | 3.7309E-05 |
| U2 | -6.9433727 | 0.00119911 |
| CD177 | -6.9495032 | 2.59E-05 |
| HOXB7 | -6.9930787 | 2.17E-05 |
| PENK | -7.0069871 | 2.07E-05 |
| CLEC12A | -7.0079194 | 2.26E-05 |
| TUBBP5 | -7.012103 | 1.98E-05 |
| FOXC2 | -7.1014053 | 1.42E-05 |
| PSG8 | -7.123825 | 1.38E-05 |
| CCL2 | -7.1252236 | 6.23E-16 |
| MRAP2 | -7.2755909 | 2.86E-06 |
| GDF3 | -7.2885792 | 2.77E-06 |
| PSG1 | -7.4494228 | 1.50E-06 |
| FENDRR | -7.5627695 | 2.40E-06 |
| CYP1B1 | -7.6561625 | 1.81E-18 |
| TCF21 | -7.7334164 | 1.27E-06 |
| PSG5 | -7.7999045 | 9.86E-07 |
| PSG9 | -7.8771357 | 6.83E-07 |
| RSPO2 | -8.1009905 | 4.19E-14 |
| IGF1 | -8.1132777 | 2.66E-07 |
| DIO2 | -8.6511741 | 2.59E-08 |
| HOXA13 | -8.8460306 | 1.08E-08 |
| XIST | -13.146847 | 4.32E-19 |

**Sup. Table 9**: Binding site analysis of FOXA, HNF4, and HNF1 binding sites

| **Hepatic TF** | **Binding Sites to Neural TF Promoters/Enhancers** | **Binding Sites to Mesoderm TF Promoters/Enhancers** |
| --- | --- | --- |
| FOXA1 | SOX4, KLF7, NOTCH1, KLF4, NR5A2, SOX2 | SNAI1, RUNX2, TWIST2, FOXC1 |
| FOXA2 | KLF7, OCLN, PAX6, NES, SOX2 | SNAI1, RUNX2, TWIST2, FOXC1 |
| FOXA3 | SOX4, KLF7, OCLN, SOX9 | SNAI1, RUNX2, TWIST2, FOXF1 |
| HNF4A | KLF7, OCLN, NOTCH1, SOX17, NR5A2 | SNAI1, RUNX2, TWIST2 |
| HNF1A | SOX9 |  |

| **GENE** | **FORWARD** | **REVERSE** |
| --- | --- | --- |
| AFP | CTTTGGGCTGCTCGCTATGA | GCATGTTGATTTAACAAGCTG |
| ALBUMIN | ACCCCACACGCCTTTGGCACA | CACACCCCTGGAATAAGCCGA |
| CDX2 | GGGCTCTCTGAGAGGCAGGT | CCTTTGCTCTGCGGTTCTG |
| CD31 | CAGAGGTCTTGAAATACAGG | ATGGAGATTACTGATGGTCC |
| CK19 | AACCATGAGGAGGAAATCAG | CATGACCTCATATTGGCTTC |
| EPCAM | GTATGAGAAGGCTGAGATAAAG | CTTCAAAGATGTCTTCGTCC |
| FOXA2 | GGGAGCGGTGAAGATGGA | TCATGTTGCTCACGGAGGAGT |
| FOXF1 | AGCTGCAAGGCATCC | AGTGCATGGAAGAGGAC |
| HNF4A | CATGGCCAAGATTGACAACCT | TTCCCATATGTTCCTGCATCA |
| NKX2.5 | AAAGCCTGAAATTTTAAGTCAC | CAGCATTTGTAGAAAGTCAGG |
| PDX1 | GCGTTGTTTGTGGCTGTTGCG | AGCTTCCCCGCTGTGTGTGTT |
| PROX1 | AGTTCAACAGATGCATTACC | TCTCTGGTTATAGACAGCTC |
| RUNX2 | AAGCTTGATGACTCTAAACC | TCTGTAATCTGACTCTGTCC |
| SOX2 | TGGACAGTTACGCGCACAT | CGAGTAGGACATGCTGTAGGT |
| TBX3 | AGACACAAAAAGGAGAATGG | AATCTTTGAGGTTCGATGTC |
| TTR | GCTGGGAGCAGCCATCACAGA | CACTTGGATTCACCGGTGCCC |
| VEGFR2 | GTACATAGTTGTCGTTGTAGC | TCAATCCCCACATTTAGTTC |

**Sup Table 10. Primer list**
